# High-resolution functional description of vaginal microbiomes in health and disease

**DOI:** 10.1101/2023.03.24.533147

**Authors:** Johanna B. Holm, Michael T. France, Pawel Gajer, Bing Ma, Rebecca M. Brotman, Michelle Shardell, Larry Forney, Jacques Ravel

## Abstract

**Background:** A *Lactobacillus-*dominated vaginal microbiome provides the first line of defense against numerous adverse genital tract health outcomes. However, there is limited understanding of the mechanisms by which the vaginal microbiome modulates protection, as prior work mostly described its composition through morphologic assessment and marker gene sequencing methods that do not capture functional information. To address this limitation, we developed metagenomic community state types (mgCSTs) which uses metagenomic sequences to describe and define vaginal microbiomes based on both composition and function.

**Results:** MgCSTs are categories of microbiomes classified using taxonomy and the functional potential encoded in their metagenomes. MgCSTs reflect unique combinations of metagenomic subspecies (mgSs), which are assemblages of bacterial strains of the same species, within a microbiome. We demonstrate that mgCSTs are associated with demographics such as age and race, as well as vaginal pH and Gram stain assessment of vaginal smears. Importantly, these associations varied between mgCSTs predominated by the same bacterial species. A subset of mgCSTs, including three of the six predominated by *Gardnerella* mgSs, as well as a mgSs of *L. iners*, were associated with a greater likelihood of Amsel bacterial vaginosis diagnosis. This *L. iners* mgSs, among other functional features, encoded enhanced genetic capabilities for epithelial cell attachment that could facilitate cytotoxin-mediated cell lysis. Finally, we report a mgSs and mgCST classifier as an easily applied, standardized method for use by the microbiome research community.

**Conclusions:** MgCSTs are a novel and easily implemented approach to reducing the dimension of complex metagenomic datasets, while maintaining their functional uniqueness. MgCSTs enable investigation of multiple strains of the same species and the functional diversity in that species. Future investigations of functional diversity may be key to unraveling the pathways by which the vaginal microbiome modulates protection to the genital tract. Importantly, our findings support the hypothesis that functional differences between vaginal microbiomes, including those that may look compositionally similar, are critical considerations in vaginal health. Ultimately, mgCSTs may lead to novel hypotheses concerning the role of the vaginal microbiome in promoting health and disease, and identify targets for novel prognostic, diagnostic, and therapeutic strategies to improve women’s genital health.

## BACKGROUND

The vaginal microbiome plays a vital role in gynecological and reproductive health. *Lactobacillus* predominated vaginal microbiota constitute the first line of defense against infection. Protective mechanisms include lactic acid production by *Lactobacillus* spp., which acidifies the vaginal microenvironment and elicits anti-inflammatory effects [1-4]. This environment wards off non-indigenous organisms, including causative agents of sexually transmitted infections (STIs) like HIV and pathogenic bacteria associated with bacterial vaginosis (BV) [5-7]. However, vaginal *Lactobacillus* spp. are functionally diverse. For example, *L. crispatus*, and *L. gasseri* are capable of producing both the D- and L-isomers of lactic acid, *L. jensenii* produces only the D-isomer, and *L. iners* only the L-isomer [4, 8]. These key features have implications for susceptibilities to pathogens [9, 10].

The vaginal microbiota has been previously shown to cluster into community state types (CSTs) that reflect differences in bacterial species composition and abundance [1, 11]. *Lactobacillus* spp. predominate four of the five CSTs (CST I: *L. crispatus*; CST II: *L. gasseri*; CST III: *L. iners*, CST V: *L. jensenii*). In contrast, CST IV communities are characterized by a paucity of lactobacilli and the presence of a diverse array of anaerobes such as *Gardnerella vaginalis* and “*Ca.* Lachnocurva vaginae”. CST IV is found, albeit not exclusively, during episodes of BV, a condition associated with increased risk to sexually transmitted infections, including HIV, as well as preterm birth and other gynecological and obstetric adverse outcomes [12-20]. BV is clinically defined by observing 3 of 4 Amsel’s criteria (Amsel-BV; vaginal pH > 4.5, abnormal discharge, and on wet mount, presence of clue cells and fishy odor with 10% KOH) [21]. Patients presenting with symptoms and satisfying the Amsel’s criteria (symptomatic Amsel-BV) are treated with antibiotics, however, efficacy is poor, and recurrence is common [21-24]. In research settings, BV is often defined by scoring Gram stained vaginal smears (Nugent-BV) [25] or molecular typing of bacterial composition by sequencing marker genes (molecular-BV) [26]. There is no definition of BV that relies on both the composition and function of the microbiome.

Species-level composition of the vaginal microbiota may not suffice to accurately capture associations between the vaginal microbiome and outcomes of interest because functional differences exist between strains of the same species. For example, in the skin microbiome, strains of *Staphylococcus aureus* or *Streptococcus pyogenes* elicit different acute immune responses [27]. Similarly, genomic and functional analyses of *Lactobacillus rhamnosus* strains demonstrate distinct adaptations to specific niches (for example, the gut versus the oral cavity) [28]. While functional differences likely exist between strains of the same species in the vaginal microbiota, metagenomic studies show that combinations of multiple strains co-exist within a single vaginal microbiome [29, 30]. These strain assemblages are known as metagenomic subspecies or mgSs [29], and are important to consider as they potentially impact the functional diversity and resilience of a species in a microbiome. Determining the mechanistic consequences and health outcomes associated with metagenomic subspecies may improve precision of risk estimates and interventions.

To integrate the taxonomic composition and functional potential of vaginal microbiomes, we developed metagenomic community state types (mgCSTs). MgCSTs are composed of unique combinations of mgSs. We developed and validated a two-step classifier that assigns metagenomic subspecies and mgCSTs and is designed to work in concert with the vaginal non-redundant gene database, VIRGO [29]. This easy-to-use classifier will facilitate reproducibility and comparisons across studies.

## RESULTS

### Metagenomic community state types (mgCST) of the vaginal microbiome

We evaluated the within-species bacterial genomic diversity in 1,890 vaginal metagenomes of reproductive-age participants from 1,024 mostly North American women (98.7% of samples) (**Table 1 SUBJECT DEMOGRAPHICS**). Vaginal metagenomes derived from five cohort studies as well as metagenomes generated to build the vaginal non-redundant gene database (VIRGO, [29]) were used to construct mgCSTs (see **Methods**). In total, 135 metagenomic subspecies (mgSs) from 28 species were identified by hierarchical clustering of species-specific gene presence/absence profiles (**Table S1 Subspecies**). Subsequent clustering of samples based on mgSs compositional data produced 27 mgCSTs (**Table 2 mgCST)**. MgCSTs consisted of mgSs from commonly observed vaginal species including *L. crispatus* (mgCST 1-6, 19% of samples), *L. gasseri* (mgCST 7-9, 3% of samples), *L. iners* (mgCST 10-14, 23% of samples), *L. jensenii* (mgCST 15 and 16, 4.6% of samples), “*Ca.* Lachnocurva vaginae” (mgCST 17-19, 7.5% of samples), *Gardnerella* (mgCST 20-25, 36.3% of samples) and *Bifidobacterium breve* (mgCST 26, 0.74% of samples) (**Figure 1 mgCST Heatmap**). MgCST 27 (5.5% of samples) contained less-common species such as *Streptococcus anginosus* or had no predominant taxon. MgCST 2 (n=39 samples from 26 women), mgCST 14 (n=34 samples from 25 women), and mgCST 21 (n=37 samples from 21 women), were only comprised of samples from reproductive aged women in Alabama enrolled in the UMB-HMP cohort (**Table 2 mgCST**). Metagenomic CSTs expand amplicon-based CSTs as multiple mgCSTs are predominated by the same species, but a different mgSs of that species (**Supplemental Figure 1 Valencia, TABLE 2 mgCST**).

**Table 1.**
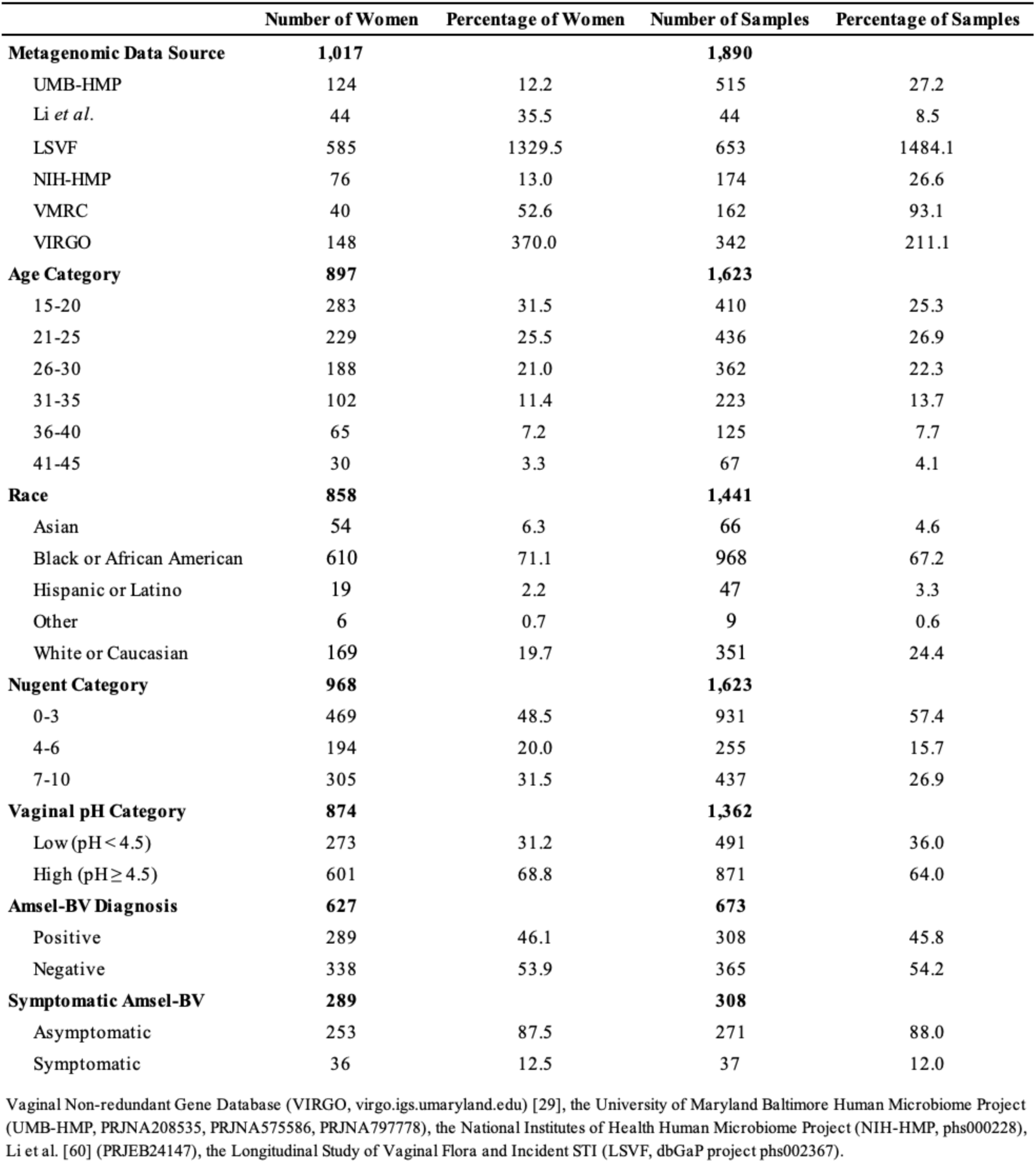
Demographic information for all women included in this study. Some women contributed multiple samples.

**Table 2.**
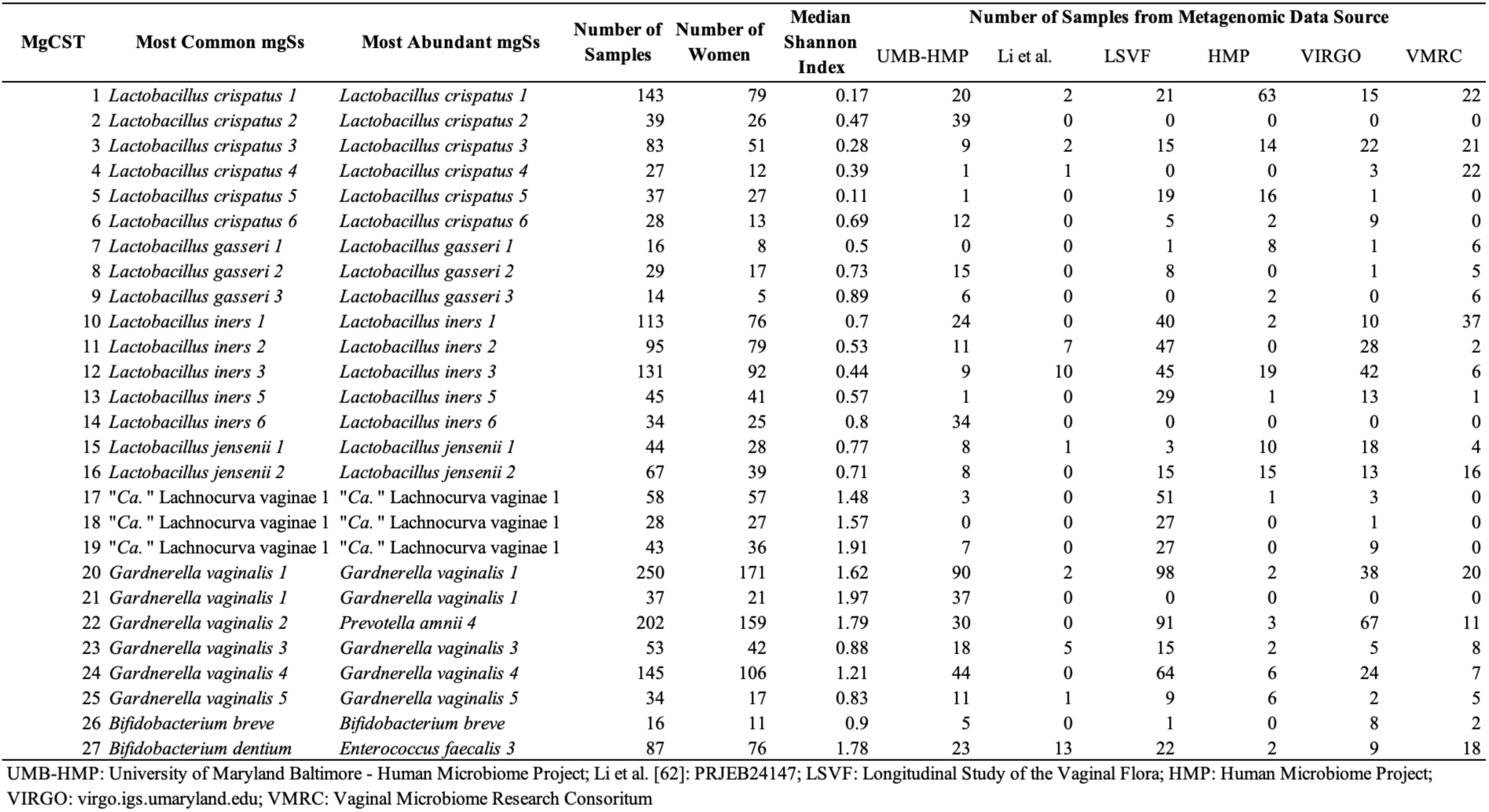
Metagenomic Community State Types (mgCSTs) of the vaginal microbiome are dominated by different metagenomic subspecies.

**Figure 1.**
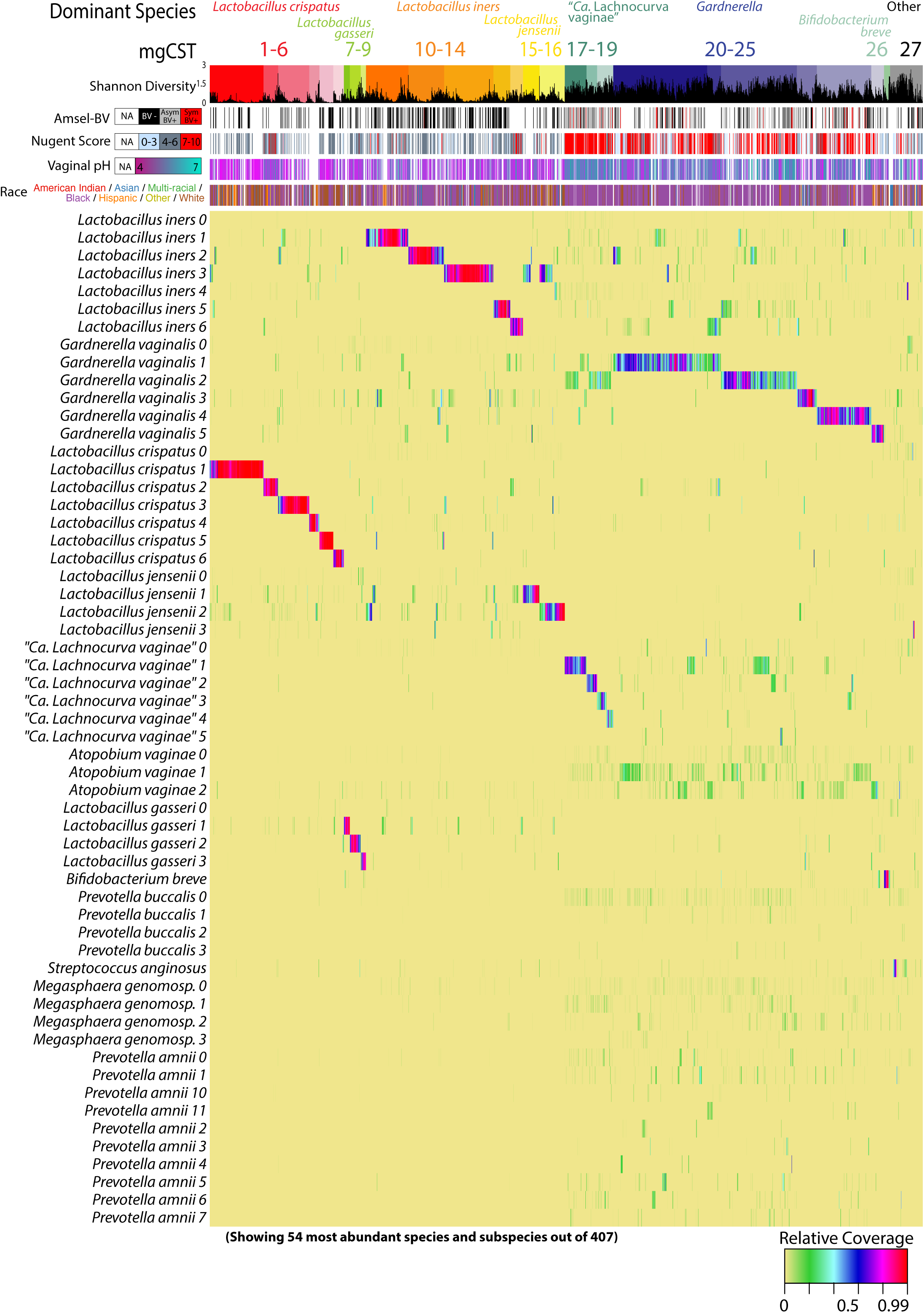
Vaginal Metagenomic Community State Types (mgCSTs). Using 1,890 metagenomic samples, 27 mgCSTs were identified: mgCSTs 1-16 are predominated by metagenomic subspecies of *Lactobacillus* spp., mgCSTs 17-19 by metagenomic subspecies of “*Ca*. Lachnocurva vaginae”, mgCSTs 20-25 by metagenomic subspecies of the genus *Gardnerella*, and mgCST 27 contains samples without a predominant metagenomic subspecies.

### Vaginal mgCSTs and Demographics

#### Race and Age

Race information was available for 1,441 samples from 858 women. Most women identified as either Black (71%) or White (20%), and the remainder identified as Asian (6.3%), Hispanic (2.2%), or other (<1%) (**Table 1 SUBJECT DEMOGRAPHICS**). Age was also reported for 1,623 samples from 897 individuals and ranged from 15-45 years old. After adjusting for between-cohort heterogeneity, certain races and age categories were associated with mgCSTs (**Figure 2**). The vaginal microbiomes of Black women were more likely to be classified as *Gardnerella* mgCST 22 (p = 0.0006) and least likely to be in *L. crispatus* mgCST 1 (p = 0.005) as compared with microbiomes for other races (**Table S2 STATS SUMMARY**). Microbiomes classified as mgCST 6 were more likely to be from White women than other races (p = 0.002). *L. iners* mgCST 12 was most common among Hispanic women (p=0.0001), and *L. iners* mgCSTs 10 and 14 were absent in Asian women (**Figure 2c**). MgCSTs predominated by “*Ca.* Lachnocurva vaginae” (mgCSTs 17-19) were also not observed in Asian women, consistent with previous reports on that species (**Figure 2c**) [11]. In mgCST 27, women were less likely to be Black (p=0.01) and more likely to be in the oldest age category (41-45, p = 0.04) as compared with other mgCSTs.

**Figure 2.**
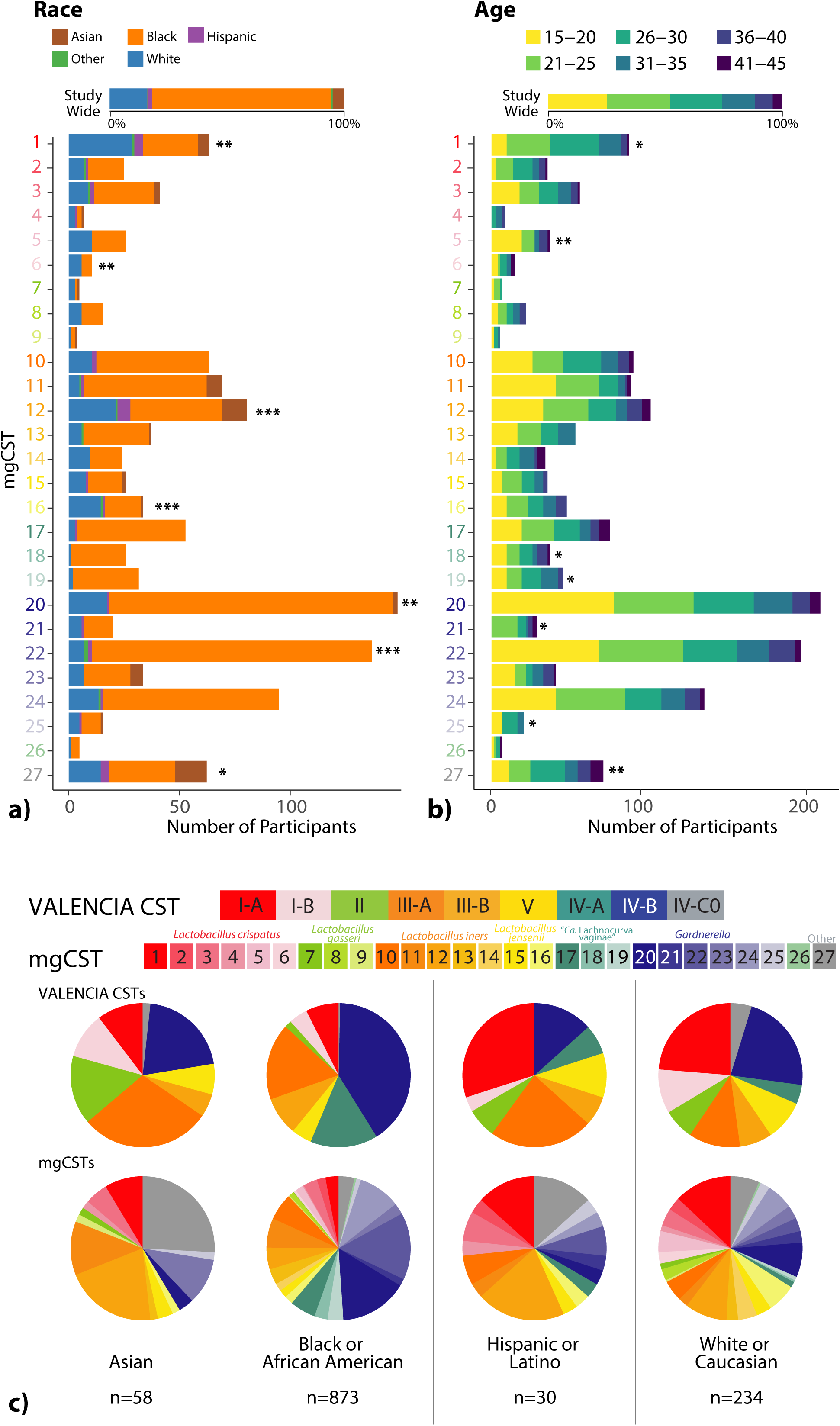
The distribution of (a) race (n=1,441 samples) and (b) age (n=1,623 samples) categories across mgCSTs. Within-mgCST distribution is compared to study-wide distribution (*: p < 0.05, **: p < 0.01, ***: p < 0.001). (c) The distribution of mgCSTs across race.

#### Nugent Scores and Vaginal pH

Of the 968 women for which Nugent scores were available, 48% had low Nugent scores (0-3), 20% had intermediate scores (4-6), and 32% had high scores (7-10) (**Table 1**). Vaginal pH was also available for 979 women and of these 31% had low pH < 4.5, and 69% had high pH ≥ 4.5 (**Table 1**). Both Nugent score and vaginal pH were associated with mgCSTs after adjusting for between-cohort heterogeneity (**Figure 3**). Of all *L. crispatus* mgCSTs, mgCST 2 had the most representation of different Nugent categories, with 61%, 14%, and 25% of samples having low, intermediate, or high Nugent scores, respectively (**Figure 3a**). Communities predominant in “*Ca.* Lachnocurva vaginae” mgCSTs 17, 18, and 19 had the highest percentages of high Nugent scores (7-10), (94%, 96%, and 87% of samples, respectively); and these mgCSTs were also associated with high vaginal pH (p = 6.3 e^-7^, **Figure 3b**). Notably, intermediate Nugent scores were common among *Gardnerella* predominated mgCSTs, especially in mgCSTs 25 (69% of samples).

**Figure 3.**
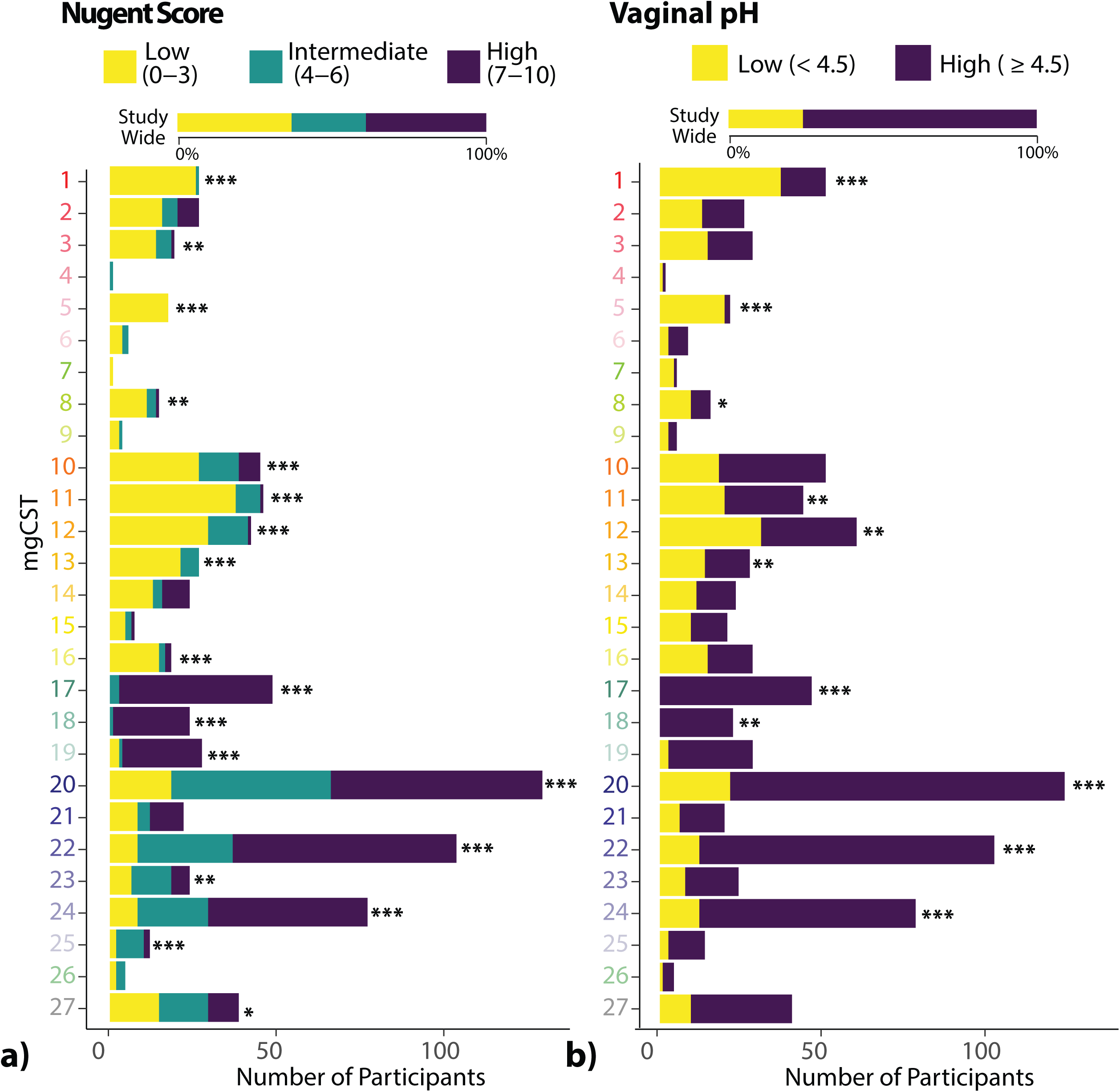
The distribution of (a) Nugent score (n=968), and (b) vaginal pH (n=979) categories. Within-mgCST distribution is compared to study-wide distribution (*: p < 0.05, **: p < 0.01, ***: p < 0.001).

#### Amsel-BV and Vaginal Symptoms

Of 627 women, each with a sample associated with clinical examination data (n=607 from LSVF cohort, n=20 from HMP cohort), the proportion of positive Amsel-BV diagnoses (including both asymptomatic and symptomatic Amsel-BV) was 46%. Twelve percent of Amsel-BV cases were symptomatic. Diagnosis of Amsel-BV was associated with mgCSTs (**Figure 4a**). There were no Amsel-BV diagnoses in mgCSTs predominated by *L. crispatus, L. jensenii*, or *L. gasseri*. *L. iners* predominated mgCSTs 10-13 were negatively associated Amsel-BV diagnoses (p = 9.6e^-4^) but contained some positive Amsel-BV diagnoses in mgCSTs 10, 11, and 13 (11%, 15%, 18% of women, respectively) (**Figure 4a** and **Table S2 STATS SUMMARY**). *L. iners* mgCST 12 contained only a single (asymptomatic) positive Amsel-BV diagnosis out of 39 women. Women with “*Ca.* Lachnocurva vaginae” mgCSTs 17-19 were more likely to have been diagnosed with Amsel-BV (87%, 88%, and 89%, respectively, p = 1.8e^-5^). *Gardnerella* predominated mgCSTs 20, 22, and 24 also had significantly more positive Amsel-BV diagnoses than the study-wide proportion (69%, 73%, and 66%, respectively, p = 1.5e^-3^), while 75% of *Gardnerella* predominated mgCST 23 samples were Amsel-BV negative (p=0.09). MgCST 24 contained significantly more symptomatic cases than expected (26% of 43 individuals, p=0.008, **Figure 4b, Table S2 STATS SUMMARY**). Though not statistically significant, *“Ca.* Lachnocurva vaginae” mgCST 19 also may have a higher-than-expected proportion of symptomatic Amsel-BV cases (17.4%).

**Figure 4.**
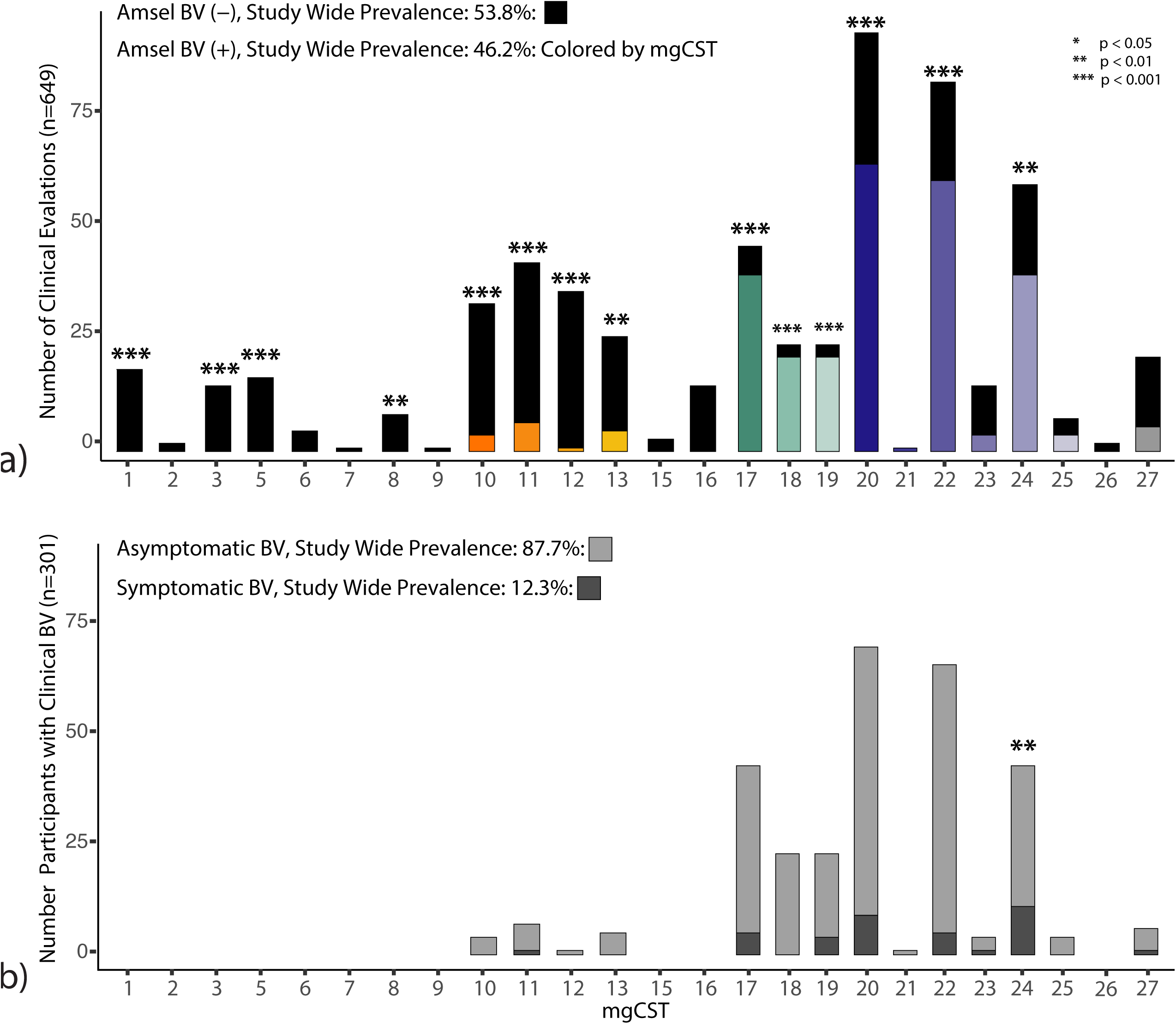
Clinically diagnosed Amsel bacterial vaginosis (a) and symptomatic Amsel bacterial vaginosis (b) are associated with mgCSTs (*: p < 0.05, **: p < 0.01, ***: p < 0.001).

### Functional potential of mgCSTs and metagenomic subspecies

#### L. crispatus *mgCSTs differ by species diversity, stability, and the potential to produce D-lactic acid*

*L. crispatus* is known to produce both L- and D-lactic acid, which acidifies the vaginal environment and confers protective properties [4, 10, 31, 32]. VIRGO identified two L- and two D-lactate dehydrogenase genes in *L. crispatus*. All genes were present in *L. crispatus* mgCSTs except for mgCST 2. Samples in mgCST 2 were missing a D-lactate dehydrogenase gene (V1806611) that has 96.1% identity to a functionally validated ortholog, P30901.2 (**Figure 5a**) [33]. The other D-lactate dehydrogenase, V1891370, is found in all *L. crispatus* mgCSTs but only 82.4% identical to P30901.2. It contains a 55 aa insertion after V101 (position in P30901.2) and a point mutation at position 218 (D218Y) located within a NAD binding site domain. The absence of V1806611 may have functional consequences for microbiomes in mgCST 2. Additionally, samples in mgCST 2 have fewer estimated numbers of *L. crispatus* strains compared to other mgCSTs (**Figure 5b**). Thus, it is likely that an *L. crispatus* strain (or strains) containing V180661 is absent from mgCST 2 samples. Interestingly, the median vaginal pH in mgCST 2 was 4.7, while in mgCST 1 it is 4.0 (1^st^-3^rd^ quartile: 3.8-4.2, **Figure 5c**). Correspondingly, mgCST 2 samples contained a higher Shannon’s H index than mgCST 1 (**Figure 5d**). All samples in mgCST 2 contained genes from “*Ca.* Lachnocurva vaginae”, *Finegoldia magna*, *Peptoniphilus harei*, *P. lacrimalis*, *Prevotella timonensis, P. disiens*, *P. buccalis,* and *Propionibacterium,* albeit at low relative abundances (<1%). We hypothesized that the observed heterogeneity in the compositions of mgCST 2 might result in lower microbiome stability than mgCST 1. Using longitudinal data from the UMB-HMP study, Yue-Clayton θ of daily bacterial composition data over 10 weeks was calculated as an estimate of community stability. Compared to mgCST 1, mgCST 2 samples were indeed significantly less stable (t = 4.073, df = 47.942, p-value < 0.001, **Figure 5e**).

**Figure 5.**
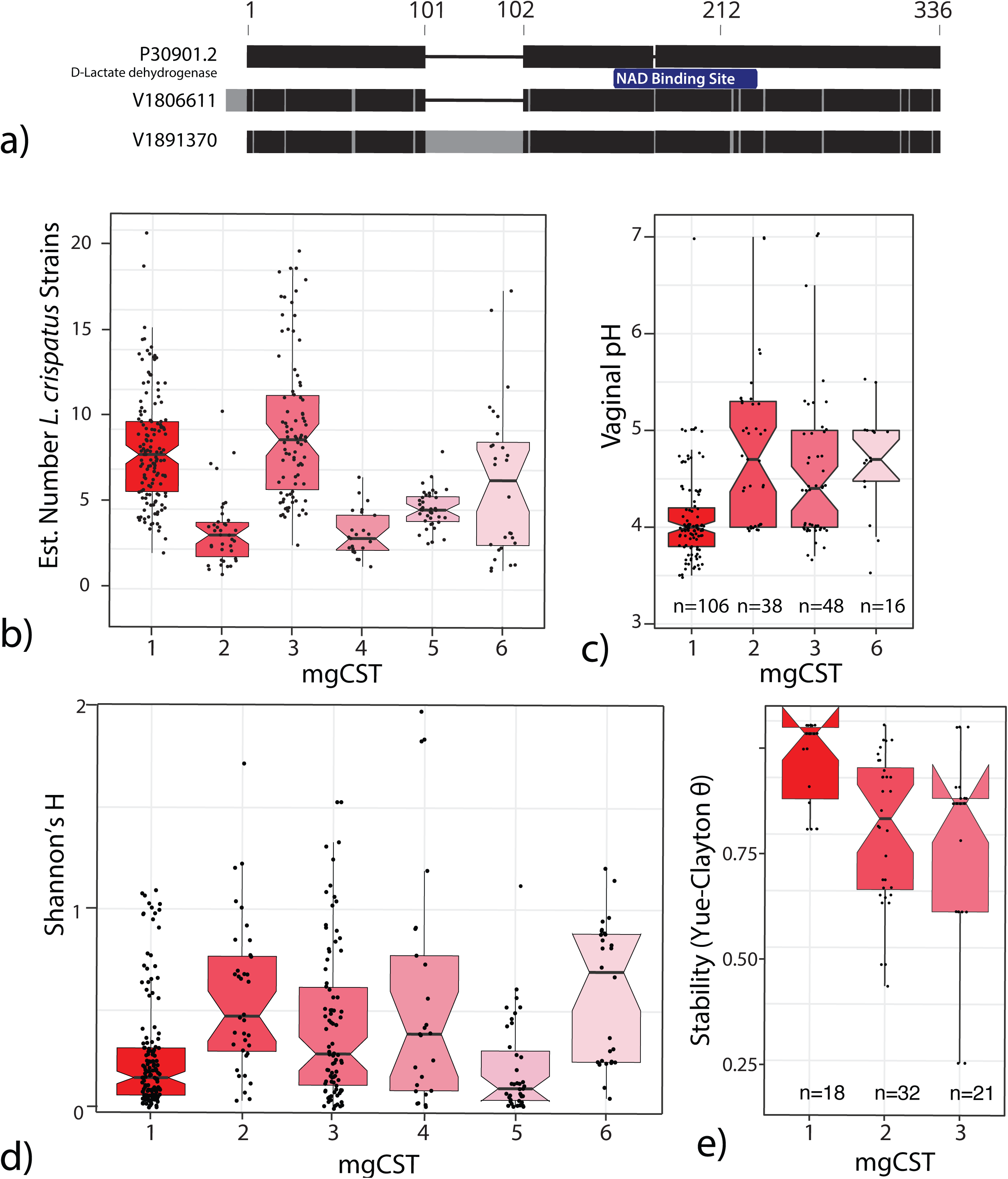
a) D-lactate dehydrogenase orthologs in VIRGO compared to reference P30901.2. b) MgCST 2 contains fewer estimated strains of *L. crispatus.* c) On average, vaginal pH is higher in mgCST 2. d) Shannon’s H is higher in mgCST 2 than mgCST 1 or 3. e) Microbiome stability is lower in mgCST 2.

#### L. iners *metagenomic subspecies are associated with Amsel-BV diagnoses*

The role of *L. iners* in the vaginal microbiome is not fully understood because it is implicated in both healthy and diseased states [34]. *L. iners* is represented by six mgSs. Predominance by *L. iners* mgSs 4 did not define a mgCST (**Figure 1**). Instead, *L. iners* mgSs 4 was present in relatively lower abundances (median: 1.2%, IQR: 1.9%) in 257 microbiomes from BV-like mgCSTs including *“Ca.* Lachnocurva vaginae” mgCSTs 16, 17, and 18, and *Gardnerella* mgCSTs 19 and 24. Seventy percent of samples containing *L. iners* mgSs 4 were positive Amsel-BV cases which is significantly greater than the proportion of cases harboring any *L. iners* mgSs (45.8%, p=1.1^e-6^, **Figure 6a**). Conversely, *L. iners* mgSs 3 was associated with negative Amsel-BV diagnoses (92% Amsel-BV negative, p=1.6^e-9^).

**Figure 6.**
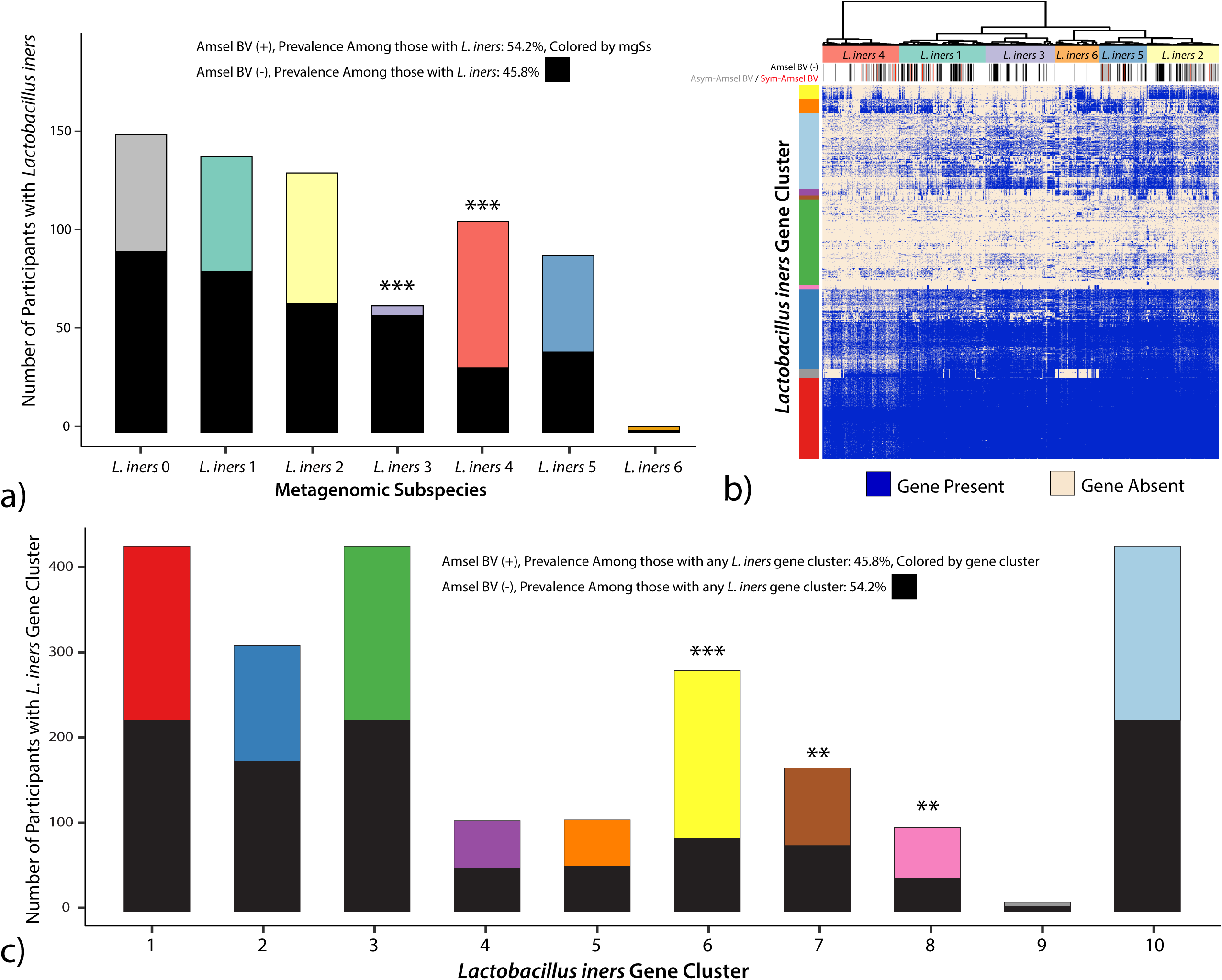
a) Clinically diagnosed Amsel-BV is associated with *L. iners* metagenomic subspecies (mgSs). b) Gene clusters present in *L. iners* mgSs. c) *L. iners* gene clusters 6 (yellow), 7 (brown) and 8 (pink) are associated with positive Amsel-BV diagnosis. Gene cluster 2 (dark blue) is associated with negative Amsel-BV diagnoses. (*: p < 0.05, **: p < 0.01, ***: p < 0.001).

We next evaluated if *L. iners* genes were associated with Amsel-BV. Most samples in *L. iners* mgSs 4 contained genes from cluster 6 (yellow gene cluster, **Figure 6b**). There were significantly more positive Amsel-BV diagnoses among subjects containing *L. iners* gene cluster 6 (69.4%, p=2.1^e-15^), 7 (53.9%, p=0.004), or 8 (60.2%, p=0.036) compared to samples containing any other *L. iners* gene cluster (45.8%, **Figure 6c**). Gene products unique to *L. iners* gene cluster 6 had significant similarity to virulence factors that could contribute to *L. iners* ability to thrive in dynamic vaginal states. Such factors include serine/threonine-protein kinases (STPKs), SHIRT domains known as “periscope proteins” which regulate bacterial cell surface interactions related to host colonization [35], CRISPR-*cas,* β-lactamase and multidrug resistance (MATE), and bacterocin exporters (**Table S3**). Gene products in cluster 7 included ParM, which plays a vital role in plasmid segregation, pre-protein translocation and membrane anchoring (SecA, SecY, sortase), defense mechanism beta-lytic metallopeptidase, and mucin-binding and internalin proteins. In *Listeria monocytogenes*, internalin A mediates adhesion to epithelial cells and host cell invasion [36]. Phage-like proteins in gene group 8 suggest the presence of mobile elements. The presence of the highly-conserved *L. iners* pore-forming cytolysin, inerolysin [37], did not differ by mgSs.

#### *Diversity of* Gardnerella *genomospecies is associated with an increase in virulence factors*

As previously mentioned, positive Amsel-BV diagnoses were common in *Gardnerella* mgCSTs 20, 22, and 24, while mgCST 23 contained more negative Amsel-BV diagnoses. Symptomatic BV cases were more common in mgCST 24 than in all other mgCSTs. By mapping the available genomes of various *Gardnerella* genomospecies [38] to VIRGO, we determined that each *Gardnerella* mgSs consists of a unique combination of *Gardnerella* genomospecies (**Figure 7a**). Compared to other *Gardnerella* mgCSTs, mgCSTs 20-22 contain a greater number of *Gardnerella* genomospecies than mgCSTs 23-25. MgCST 24 samples are predominated by *Gardnerella* mgSs 4 and largely consists of *G. swidsinkii* and *G. vaginalis* genes. This suggests the diversity and types of *Gardnerella* genomospecies may be important determinants of the pathogenicity of mgCSTs. For example, there are more gene variants of common *Gardnerella* virulence factors like sialidase and vaginolysin in samples with more genomospecies (**Figure 7b**).

**Figure 7.**
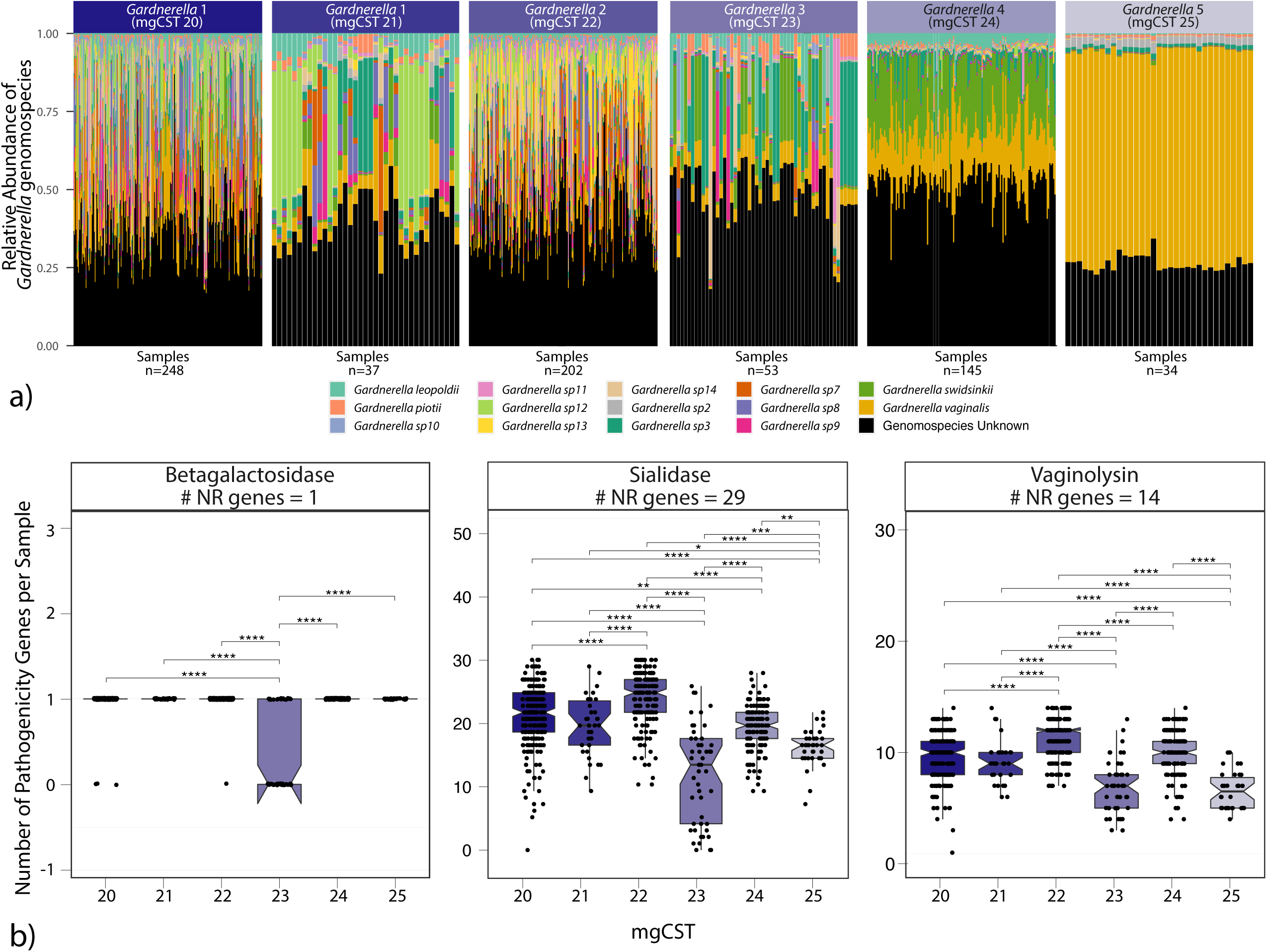
a) The distribution of *Gardnerella* genomospecies across *Gardnerella* mgSs. b) known pathogenicity genes are differently distributed across *Gardnerella* mgCSTs.

### Automated Classification of mgCSTs using Random Forest Models

Random forest models were built for each of the 135 mgSs identified and used to perform mgSs assignments (**see Methods**). The misclassification error for mgSs assignment ranged from 0-30% (**Supplemental Figure 2 mgSs Misclassification Error**). The error estimates for most major vaginal taxa were near or less than 10%, with *L. gasseri* having the lowest (2.2%). *L. iners* consistently provided higher misclassification error estimates (20%) regardless of attempts to fine-tune the model and was likely the result of high genetic heterogeneity within *L. iners* mgSs. Following assignment of mgSs, mgCSTs were assigned using the nearest centroid classification method, as previously used for vaginal taxonomy-based community state type assignments [11]. The mean classification error was 9.6%, with some mgCSTs classified more accurately than others (**Supplemental Figure 3 mgCST Misclassification Error)**. The mgCST classifier is packaged into an R script, is available at https://github.com/ravel-lab/mgCST-classifier and uses direct outputs from VIRGO.

## DISCUSSION

Recent findings that motivated development of mgCST classification are that multiple strains of the same species are commonly observed in the vaginal microbiome [29], and that samples can be clustered into metagenomic subspecies defined by unique strain combinations represented by species-specific gene sets, and thus unique sets of functions. These critical observations led us to conceptualize a vaginal microbiomes classification based on their mgSs compositions and abundance, and thus defined by both species’ composition and functions, *i.e.*, metagenomic community state types. MgCSTs describe vaginal microbiomes through a new lens, one that includes both compositional and functional dimensions.

*L. iners-*predominated vaginal microbiota have been associated with increased risks of experiencing bacterial vaginosis (BV) [39, 40]. Longitudinal observational prospective studies support this conclusion and present several critical findings: 1) *L. iners* is often detected at low to medium abundances during episodes of BV, and *L. iners* commonly dominate the vaginal microbiota after metronidazole treatment for BV and, 2) *L. iners* predominated vaginal microbiota are more prevalent prior to incidence of BV [41, 42]. We observed the frequency of *L. iners* predominated vaginal microbiota was high in Black and Hispanic women (31.4% and 36.1%, respectively), both of whom experience a disproportionate prevalence of BV in the US, with reported rates of 33.2% and 30.7%, respectively (compared to 22.7% and 11.1% in White and Asian women) [43]. Interestingly, *L. iners* predominated vaginal microbiota were even more frequent in North American Asian women in this study, as was shown previously by Ravel *et al.* [1], yet these *L. iners* predominated vaginal microbiota are not associated with higher risk of BV in these women [1]. MgCST classification provides insight into this contradiction to prevailing dogma regarding *L. iners* and increased risk of BV. We noted the absence of *L. iners* mgSs 4 in Asian women, and that *L. iners* mgSs 4 is associated with Amsel-BV, while *L. iners* mgSs 3 (predominates mgCST 12) was significantly negatively related to BV and was most frequently observed in Asian women. This is the first evidence of genetically distinct combinations of *L. iners* strains (mgSs) in healthy versus BV-associated states of the vaginal microbiome. This critical finding points to the possibility of beneficial properties associated with some *L. iners-* dominated microbiomes that had not been evidenced previously. Our analyses also identified a specific set of *L. iners* genes associated with positive Amsel-BV diagnoses. Macklaim *et al.* 2018 reported marked differences in *L. iners* gene expression between two control patients versus two diagnosed with BV, including increased CRISPR-associated proteins gene expression in BV samples [44]. However, our mgSs analysis of *L. iners* indicates that it is not simply alterations in gene expression of a common gene pool that differentiates BV from non-BV microbiomes, but *L. iners* mgSs that also differ. Microbiomes from women with positive BV diagnoses were enriched for host immune response evasion and host-colonization functions. For example, serine/threonine-protein kinases (STPKs) contribute to resistance from phagocytosis by macrophage, invasion of host cells including epithelia and keratinocytes, antibiotic resistance, disruption of the NF-κB signaling pathway, and mucin binding [45]. Bacterial attachment to host cells (clue cells) is a hallmark of high Nugent scores (a bacterial morphology-based definition of bacterial vaginosis) and a criterion in Amsel-BV diagnoses [21, 25]. Attachment of *L. iners* to epithelial cells may look like clue cells and this could lead to morphological misdiagnosis of BV. These features may make *L. iners* mgSs 4 more difficult to displace in the vaginal environment and could contribute to the common observation of *L. iners* following antibiotic treatment [46]. Interestingly, just like *L. iners* mgSs 4, mgSs of *“Ca.* Lachnocurva vaginae” were strongly associated with Amsel-BV and were also not found in the vaginal microbiomes of Asian women in this study. Together, these observations may be evidence of selective pressures by the host environment or niche specialization by vaginal bacteria. Sources of selective pressure could relate to host-provided nutrient availability (*e.g.*, mucus glycan composition), the host innate and adaptive immune system, the circulation of other species’ mgSs in a population, or any such combination.

Several distinct mgCSTs associate strongly with Amsel-BV. Critically, these data support the need for an improved definition of BV and the importance of a personalized approach to treatment. “*Ca.* Lachnocurva vaginae” predominated mgCSTs were strongly associated with asymptomatic Amsel-BV and contained more high Nugent scores than other mgCSTs. Conversely, intermediate Nugent scores were most prevalent in *Gardnerella* predominated mgCSTs, and only three of these six mgCSTs were associated with Amsel-BV, which suggests that not all *Gardnerella-*dominated microbiomes are related to Amsel-BV. *Gardnerella* contains vast genomic diversity, supporting a split into different genomospecies [38, 47, 48]. Because different genomospecies can co-exist, it is likely that *Gardnerella* predominated mgSs represent unique combinations of genomospecies and strains of these genomospecies. Greater *Gardnerella* genomospecies diversity is associated with positive Amsel-BV diagnoses in studies using qPCR or transcriptomic data to define *Gardnerella* species [47, 49-51]. Our data corroborate these reports and further indicate in mgCSTs with higher numbers of *Gardnerella* genomospecies that there are more gene variants coding for virulence factors like cholesterol-dependent pore-forming cytotoxin vaginolysin and neuraminidase sialidase present, thus expanding functional diversity and potentially explaining the association with positive Amsel-BV diagnoses [52-54]. Enumeration of *Gardnerella* genomospecies may prove to be an important diagnostic of certain “types” of Amsel-BV and could inform treatment options. For example, it is possible that harboring more *Gardnerella* genomospecies may predict BV recurrence following metronidazole treatment, suggesting the need for a different approach to treatment. Alternatively, some *Gardnerella* genomospecies may be important and novel targets of therapy.

In the clinic, antibiotic treatment is recommended for BV diagnosis (generally a point-of-care test) only when the patient reports symptoms, which is estimated to occur in fewer than half of women with BV [24, 55, 56]. In research settings, both symptomatic and asymptomatic Amsel-BV can be evaluated. Indeed, in the observational research studies included in this analysis where Amsel criteria were evaluated along with whether participants reported symptoms or not, symptomatic Amsel-BV accounted for only 12% of Amsel-BV cases and 30% of these were in mgCST 24 (dominated primarily by *Gardnerella swidsinkii* and *G. vaginalis*). We hypothesize that the inadequacy of currently recommended BV treatment may be due to the heterogeneity in the genetic make-up of the microbiota associated with BV as revealed by mgCSTs. MgCSTs reduce this heterogeneity resulting in more precise estimates of risk. Furthermore, these findings highlight the potential importance of developing specialized treatments that target “types” of BV.

The mgCST framework can also be used to identify vaginal microbiomes that are associated with positive health outcomes. For example, mgCSTs predominated by different *L. crispatus* mgSs varied in their association with low Nugent scores, the number of *L. crispatus* strains present, and the longitudinal stability of communities. The vaginal microbiome can be dynamic [57-59]. Shifts from *Lactobacillus* to non-*Lactobacillus* predominated microbiota can increase the risk of infection following exposure to a pathogen. Our study identified *L. crispatus* mgCSTs with variable stability, suggesting that not all *L. crispatus* predominated microbiomes are functionally similar and may be differently permissive to infection. Those found to be associated with higher stability may reduce the window of opportunity for pathogens to invade. Microbiome stability may be related to both the diversity of other non-*Lactobacillus* members of the microbiome and/or the number of *L. crispatus* strains present. In any case, our study shows that there is a range of protective abilities even among *L. crispatus* predominated communities. This information could be critical in selecting and assembling strains of *L. crispatus* to design novel live biotherapeutics products aimed to restore an optimal vaginal microenvironment.

It is unclear what factors contribute to vaginal strain assemblages and what rules define their biology and ecology. However, such assemblages can now be detected and further characterized using the concepts of mgSs and mgCSTs presented here. The use of metagenomic sequencing and mgSs and mgCSTs will contribute to a much-needed functional understanding of the role of the vaginal microbiome in reproductive health outcomes. Our findings support the hypothesis that genetic and functional differences between vaginal microbiomes, including those that may look compositionally similar, are critical considerations in vaginal health [7]. To aid in further exploration, we also provide a validated classifier for both mgSs and mgCSTs at https://github.com/ravel-lab/mgCST-classifier/blob/main/README.md.

## CONCLUSION

MgCSTs reveal differences between vaginal microbiome both compositionally and functionally, and thus more finely describe the vaginal microbiome. Associations between mgCSTs and bacterial vaginosis highlight the multi-faceted aspects of the condition and call for new and expanded definitions. Further, we provide tools for the classification of mgSs and mgCST that have potential for use and harmonization of analytical strategies in future studies.

## DATA AVAILABILITY

The classifiers are available to accompany VIRGO output at https://github.com/ravel-lab/mgCST-classifier.

## METHODS

### Study cohorts

Raw metagenomic data from 1,890 vaginal samples were used in this study (**Supplemental File 6**). This included publicly available metagenomes including those used in the construction of the vaginal non-redundant gene database, VIRGO (virgo.igs.umaryland.edu, n=342) [29], the University of Maryland Baltimore Human Microbiome Project (UMB-HMP, n=677, PRJNA208535, PRJNA575586, PRJNA797778)[41], the National Institutes of Health Human Microbiome Project (NIH HMP, n=174, phs000228) [60], metagenomes from Li *et al.* [61] (n=44, PRJEB24147), the Longitudinal Study of Vaginal Flora and Incident STI (LSVF, n=653, dbGaP project phs002367) [24]. All samples in LSVF (n=653) and some in UMB-HMP (n=20) had clinical diagnosis information about Amsel-BV. Amsel-BV was diagnosed based on the presence of 3 out of 4 Amsel’s criteria [21] and symptomatic Amsel-BV was diagnosed when a woman reported symptoms upon questioning [56]. At the time of these studies, gender identity information was not collected. We know all women responded to recruiting materials which included “women” or “woman”. In addition, individuals are referred to as women in previous publications, thus we refer here to individuals as “woman” or “women” to maintain consistency.

### Sequence Processing and Bioinformatics

Host reads were removed from all metagenomic sequencing data using BMTagger and the GRCh38 reference genome, and reads were quality filtered using trimmomatic (v0.38, sliding window size 4bp, Q15, minimum read length:75bp) [62]. Metagenomic sequence reads were mapped to VIRGO using bowtie (v1; parameters: -p 16 -l 25 --fullref --chunkmbs 512 --best --strata -m 20 --suppress 2,4,5,6,7,8), producing a taxonomic and gene annotation for each read. Samples with fewer than 100,000 mapped reads were removed from the analysis (n=59). The number of reads mapped to a gene was multiplied by the read length (150 bp) and divided by the gene length to produce a coverage value for each gene. Conserved domain and motif searches were performed with CD-SEARCH and the Conserved Domain Database (CDD), using an e-value threshold of 10^-4^. The taxonomic composition table generated using VIRGO were run through the vaginal CST classifier VALENCIA [11].

### Metagenomic Subspecies

For each species, a presence/absence matrix was constructed from a metagenome which included all genes with at least 0.5X coverage after normalizing for gene length. Metagenomic subspecies were generated for species present (>75% estimated median number of genes encoded in reference genomes from the Genome Taxonomy Database [63], see **Table S4 GENOME SIZES**) in >20 samples using binary gene counts and hierarchical clustering with Ward linkage of sample Jaccard distances calculated using the vegdist function from the vegan package (v2.5-5) [64] in R (v. 3.5.2). Clusters were defined using the dynamic hybrid tree cut method (v.1.62-1) and minClusterSize = 2 [65]. Clusters were tested for associations with low species coverage using logistic regression in which the mgSs was the binary outcome, the log_10_-transformed coverage of the species was the predictor, and subject ID was used as a nested random effect which accounted for multiple samples from the same subject and variations due to different source studies. Heatmaps of gene presence/absence were constructed for each species using the gplots package heatmap.2 function [66] (**Supplemental File 4**).

### Metagenomic CSTs

Using gene abundance information (normalized by gene length and sequencing depth), we estimated the proportion of vaginal species in each sample. For species that were sub-divided into mgSs, the mgSs proportion in a sample was equal to the proportion of the species in that sample. When a species was present in a sample but with too few genes present to constitute a mgSs (<75% estimated median number of genes encoded in reference genomes), it was labeled as “mgSs 0”. Samples in the resulting compositional table were hierarchically clustered using Jensen-Shannon distances. Clusters were defined using the dynamic hybrid tree cut method (v.1.62-1) [65]. A heatmap for metagenomic CSTs was produced using the gplots package heatmap.2 function (**Figure 1**) [66]. For participants in the HMP cohort who contributed longitudinal samples, the Yue-Clayton theta was measured to define microbiota stability for each subject [67]. Average per-subject stability thetas were plotted for each mgCST.

### Estimating the number of *L. crispatus* strains

The number of *L. crispatus* strains in a mgSs was estimated using a pangenome accumulation curve which was generated by mapping the gene contents of publicly available isolate genome sequences (**Supplemental File 5**) to VIRGO (blastn, threshold: 90% identity, 70% coverage). Bootstrap (n=100) combinations of N (N=1 to 61) isolates were selected and the number of unique *L. crispatus* Vaginal Orthologous Groups [VOGs; provided in the VIRGO output[29] encoded in their genomes was determined. An exponential curve relating the number of isolates to the number of VOGs detected was then fit to the resulting data and produced the equation: Y=2057N^0.14^ where Y is the number of *L. crispatus* VOGs detected, and N is the estimated number of strains. This equation was then used to estimate the number of *L. crispatus* strain’s detected in a metagenome based on the observed number of *L. crispatus* VOGs in each metagenome.

### Statistical analysis of the association between mgCST and age, race, Nugent score, vaginal pH, and BV

For those samples with race, age category, Nugent score category, vaginal pH category, or Amsel-BV diagnoses information (**TABLE**), the Cochran-Mantel-Haenszel Chi-Squared Test (CMH test, “mantelhaen.test” from the samplesizeCMH R package, v 0.0.0, github.com/pegeler/samplesizeCMH) was used to determine associations with mgCSTs while accounting for source study (the confounding variable). The CMH test evaluates associations between two binary variables (*i.e.*, “mgCST X or not” and “high Nugent score or not”). Tests were done at the subject level; if a subject had more than one sample and both samples were the same mgCST, only one sample was used, but if the mgCSTs differed, the samples were included in each.

#### Construction of the random forests for mgSs classification

We constructed random forests for classification of mgSs using the R package randomForestSRC v2.12.1R [68]. For mgSs, a random forest was built for each species (n=28) where the training data contained presence/absence values of genes. Gene presence was defined as above for mgSs. We implemented random forest classification analysis with all predictors included in a single model. For each mgSs random forest, predictors were all genes in a species. Ten-fold cross-validation (90% of data as training, 10% as testing) was performed wherein each training set was used to build and tune a random forest model using tune “tune.rfsrc”. A random forest model using optimal parameters was then used to predict mgSs classifications for the test set and out-of-bag error estimates (misclassification error) are reported. The overall misclassification error is the average misclassification error from each fold and the “correct” assignment is based on original hierarchical clustering assignment. The final models included all data and the optimal tuning parameters determined for that species.

#### Construction of the a nearest centroid classifier for mgCSTs

Using mgCSTs as defined above, reference centroids were produced using the mean relative abundances of each mgSs in a mgCST. For classification, the similarity of a sample to the reference centroids is determined using Yue-Clayton’s θ [67]. Ten-fold cross validation was applied wherein each training set was used to build “reference” centroids and each test set was used for assignment. The misclassification error was determined by subtracting the number of correct assignments (based on original hierarchical clustering assignment) divided by the total number of assignments from 1. The overall misclassification error is the average of misclassification error from each fold.

#### Running the mgCST classifier

The required inputs are direct outputs from VIRGO [29] and include the taxonomic abundance table (“summary.Abundance.txt”) and gene abundance table (“summary.NR.abundance.txt”). It is *imperative* that taxonomic and gene column headings match those output by VIRGO. The expected output is a count table with samples as rows, taxa as columns, and counts normalized by gene length as values. Additional columns indicate the sample mgCST classification and the Yue-Clayton similarity score for all 26 mgCSTs. A heatmap is also produced showing taxon relative abundances in samples, where samples are labeled with assigned mgCSTs Substantial differences may indicate either an incongruence in taxonomic or gene names or the need for an additional mgCST. The classifier is contained in an R script, which is available at https://github.com/ravel-lab/mgCST-classifier.

All bioinformatic and statistical analyses are available in R Markdown notebooks (**Supplemental File 7 mgCST_paper_bioinformatics.Rmd** and **Supplemental File 8 mgCST_paper_stats.Rmd**)

## SUPPLEMENTAL FIGURE LEGENDS

**Supplemental Figure 1.**
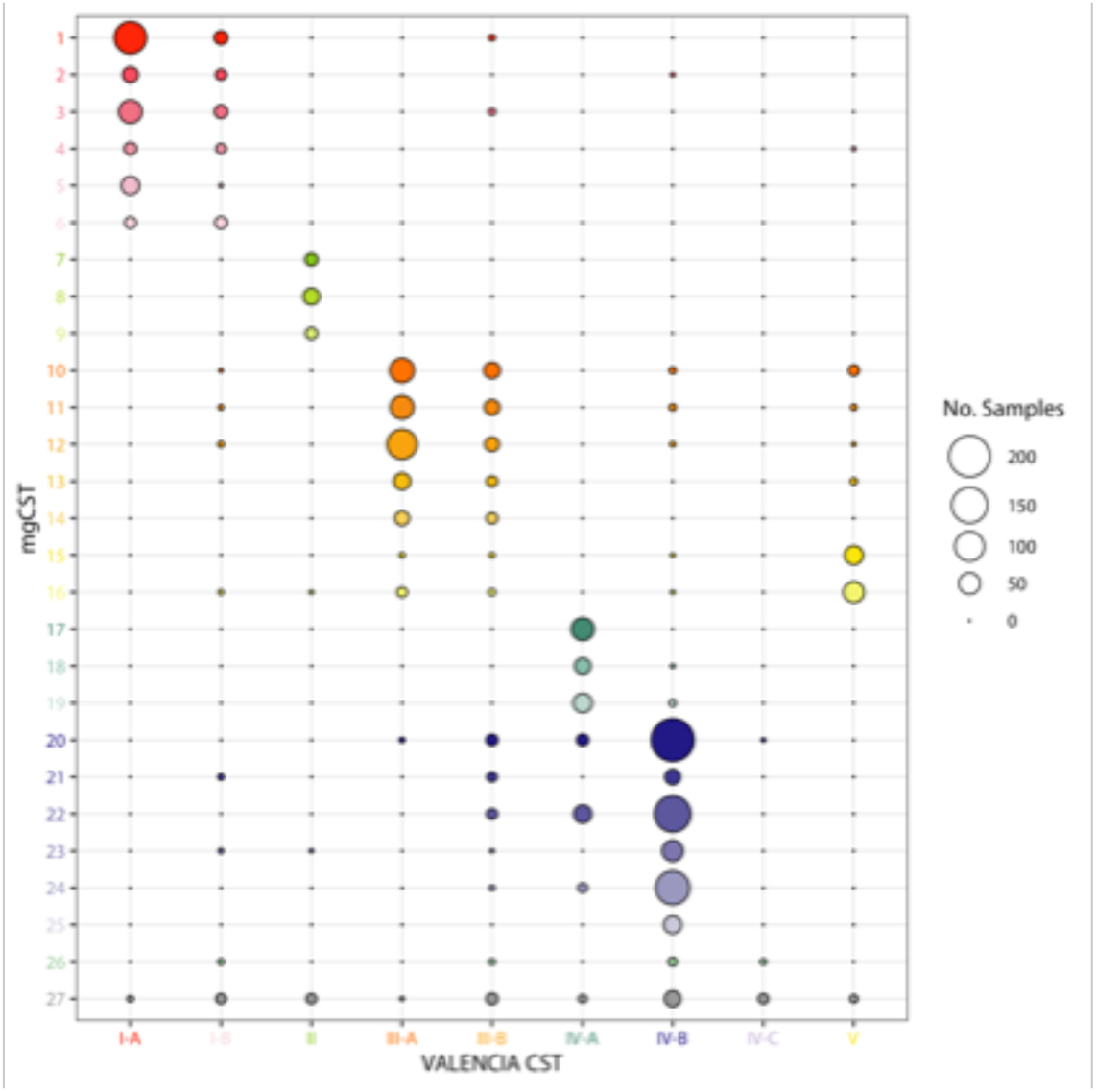
Metagenomic CSTs correspond to marker gene-based CSTs primarily through predominant taxon. Dominance by mgSs is not captured through marker-based CSTs.

**Supplemental Figure 2.**
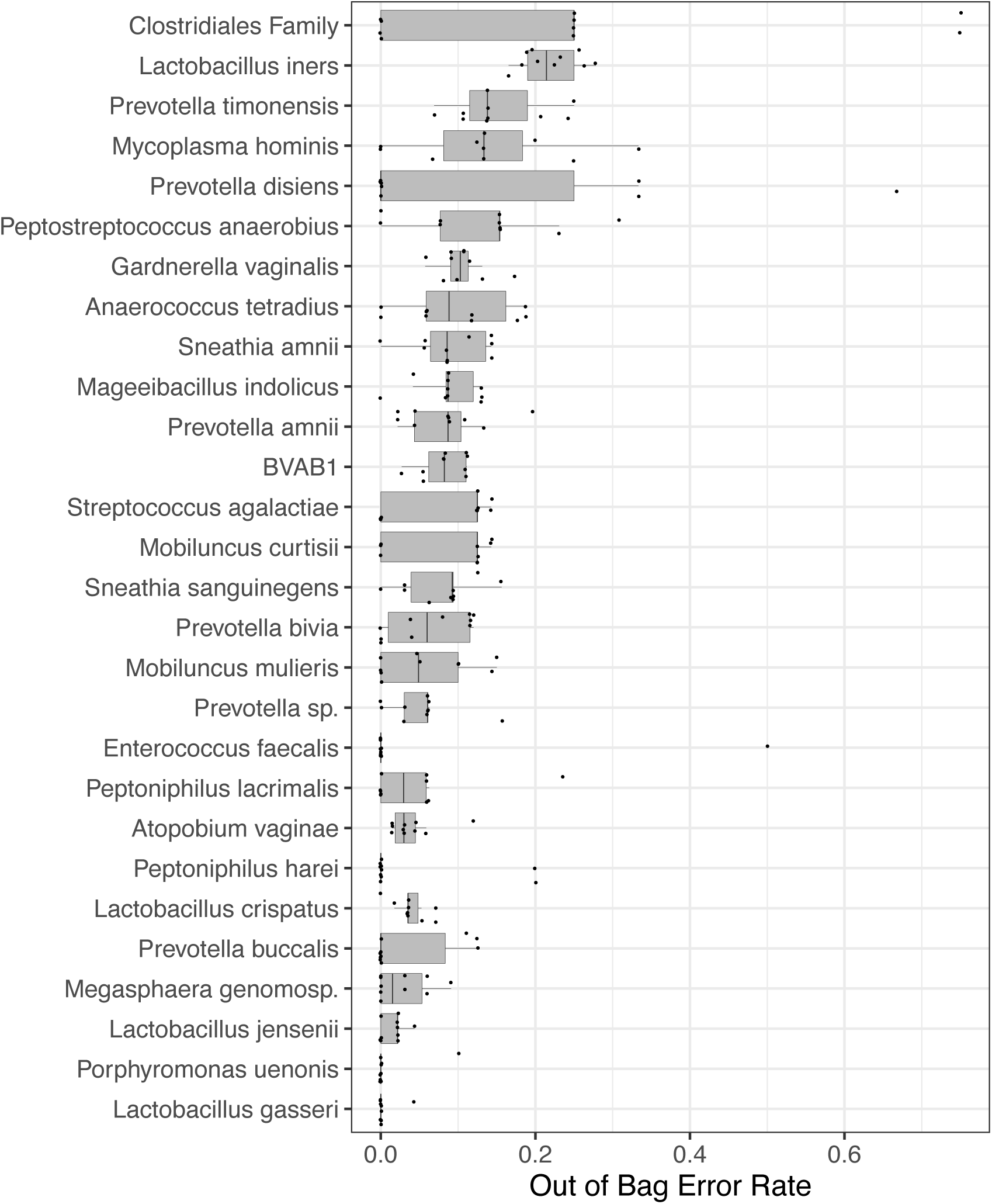
Random forest misclassification error estimates from 10-fold cross-validation for each of the species that contained metagenomic subspecies.

**Supplemental Figure 3.**
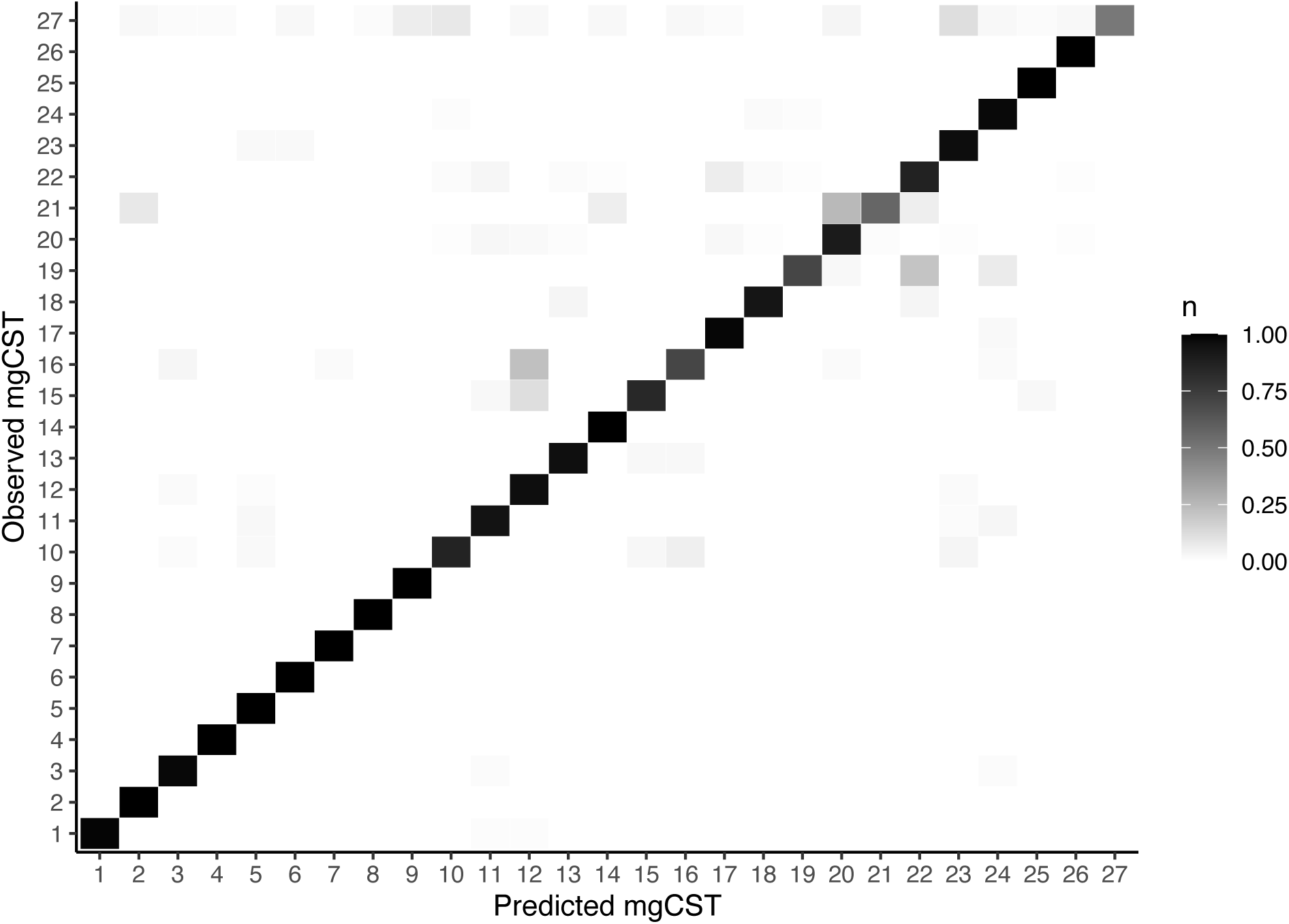
Confusion matrix and classification error estimates from 10-fold cross-validation of a nearest-centroid classifier for mgCSTs.

Supplemental File 4. Gene presence and absence heatmaps for all metagenomic subspecies.

Supplemental File 5. Gene contents of publicly available isolate metagenomes.

Supplemental Table 6. Metadata and source location for all metagenomes in this study. For Amsel-BV: clinBV.asymp=Clinical diagnosis of Amsel-BV, no symptoms reported by patient; noBV=No diagnosis of Amsel-BV after evaluation; clinBV.symp=Clinical diagnosis of Amsel-BV, symptoms reported by patient; NA=No clinical evaluation

Supplemental_File_7_mgCST_paper_bioinformatics.Rmd. Rmarkdown notebook with code used to build metagenomic subspecies and metagenomic community state types.

Supplemental_File_8_mgCST_paper_stats.Rmd. Rmarkdown notebook with code for performing all analyses and generating all figures in this manuscript.

## REFERENCES

1. Ravel J, Gajer P, Abdo Z, Schneider GM, Koenig SSK, McCulle SL, et al.: Vaginal microbiome of reproductive-age women. In: Proc Natl Acad Sci USA. vol. 108 Suppl 1; 2011: 4680-7.

2. O’Hanlon DE, Come RA, Moench TR. Vaginal pH measured in vivo: lactobacilli determine pH and lactic acid concentration. Bmc Microbiol. 2019;19(1):13; doi: 10.1186/s12866-019-1388-8.

3. Gong Z, Luna Y, Yu P, Fan H. Lactobacilli inactivate Chlamydia trachomatis through lactic acid but not H2O2. PLoS One. 2014;9(9):e107758; doi: 10.1371/journal.pone.0107758.

4. Witkin SS, Mendes-Soares H, Linhares IM, Jayaram A, Ledger WJ, Forney LJ. Influence of vaginal bacteria and D- and L-lactic acid isomers on vaginal extracellular matrix metalloproteinase inducer: implications for protection against upper genital tract infections. mBio. 2013;4(4); doi: 10.1128/mBio.00460-13.

5. Ravel J, Brotman RM. Translating the vaginal microbiome: gaps and challenges. Genome Med. 2016;8(1):35; doi: 10.1186/s13073-016-0291-2.

6. Amabebe E, Anumba DOC. The Vaginal Microenvironment: The Physiologic Role of Lactobacilli. Front Med (Lausanne). 2018;5:181; doi: 10.3389/fmed.2018.00181.

7. Ma B, Forney LJ, Ravel J. Vaginal microbiome: rethinking health and disease. Annu Rev Microbiol. 2012;66:371–89; doi: 10.1146/annurev-micro-092611-150157.

8. Boskey ER, Cone RA, Whaley KJ, Moench TR: Origins of vaginal acidity: high D/L lactate ratio is consistent with bacteria being the primary source. In: Hum Reprod. vol. 16: Oxford University Press; 2001: 1809–13.

9. Nunn KL, Wang YY, Harit D, Humphrys MS, Ma B, Cone R, et al. Enhanced Trapping of HIV-1 by Human Cervicovaginal Mucus Is Associated with Lactobacillus crispatus-Dominant Microbiota. mBio. 2015;6(5):e01084–15; doi: 10.1128/mBio.01084-15.

10. Edwards VL, Smith SB, McComb EJ, Tamarelle J, Ma B, Humphrys MS, et al. The Cervicovaginal Microbiota-Host Interaction Modulates Chlamydia trachomatis Infection. mBio. 2019;10(4); doi: 10.1128/mBio.01548-19.

11. France MT, Ma B, Gajer P, Brown S, Humphrys MS, Holm JB, et al. VALENCIA: a nearest centroid classification method for vaginal microbial communities based on composition. Microbiome. 2020;8(1):166; doi: 10.1186/s40168-020-00934-6.

12. Brotman RM, Bradford LL, Conrad M, Gajer P, Ault K, Peralta L, et al. Association between Trichomonas vaginalis and vaginal bacterial community composition among reproductive-age women. Sexually transmitted diseases. 2012;39(10):807–12; doi: 10.1097/OLQ.0b013e3182631c79.

13. Mehta SD, Donovan B, Weber KM, Cohen M, Ravel J, Gajer P, et al. The vaginal microbiota over an 8- to 10-year period in a cohort of HIV-infected and HIV-uninfected women. PLoS One. 2015;10(2):e0116894; doi: 10.1371/journal.pone.0116894.

14. Dunlop AL, Satten GA, Hu YJ, Knight AK, Hill CC, Wright ML, et al. Vaginal Microbiome Composition in Early Pregnancy and Risk of Spontaneous Preterm and Early Term Birth Among African American Women. Front Cell Infect Microbiol. 2021;11:641005; doi: 10.3389/fcimb.2021.641005.

15. Price JT, Vwalika B, Hobbs M, Nelson JAE, Stringer EM, Zou F, et al. Highly diverse anaerobe-predominant vaginal microbiota among HIV-infected pregnant women in Zambia. PLoS One. 2019;14(10):e0223128; doi: 10.1371/journal.pone.0223128.

16. Gosmann C, Anahtar MN, Handley SA, Farcasanu M, Abu-Ali G, Bowman BA, et al. Lactobacillus-Deficient Cervicovaginal Bacterial Communities Are Associated with Increased HIV Acquisition in Young South African Women. Immunity. 2017;46(1):29–37; doi: 10.1016/j.immuni.2016.12.013.

17. Atashili J, Poole C, Ndumbe PM, Adimora AA, Smith JS. Bacterial vaginosis and HIV acquisition: a meta-analysis of published studies. AIDS. 2008;22(12):1493–501; doi: 10.1097/QAD.0b013e3283021a37.

18. Elovitz MA, Gajer P, Riis V, Brown AG, Humphrys MS, Holm JB, et al. Cervicovaginal microbiota and local immune response modulate the risk of spontaneous preterm delivery. Nat Commun. 2019;10(1):1305; doi: 10.1038/s41467-019-09285-9.

19. Borgdorff H, Tsivtsivadze E, Verhelst R, Marzorati M, Jurriaans S, Ndayisaba GF, et al. Lactobacillus-dominated cervicovaginal microbiota associated with reduced HIV/STI prevalence and genital HIV viral load in African women. ISME J. 2014;8(9):1781–93; doi: 10.1038/ismej.2014.26.

20. Tamarelle J, Thiebaut ACM, de Barbeyrac B, Bebear C, Ravel J, Delarocque-Astagneau E. The vaginal microbiota and its association with human papillomavirus, Chlamydia trachomatis, Neisseria gonorrhoeae and Mycoplasma genitalium infections: a systematic review and meta-analysis. Clin Microbiol Infect. 2019;25(1):35–47; doi: 10.1016/j.cmi.2018.04.019.

21. Amsel R, Totten PA, Spiegel CA, Chen KC, Eschenbach D, Holmes KK. Nonspecific vaginitis: diagnostic criteria and microbial and epidemiologic associations. The American journal of medicine. 1983;74(1):14–22.

22. Scharbo-Dehaan M, Anderson DG. The CDC 2002 guidelines for the treatment of sexually transmitted diseases: implications for women’s health care. J Midwifery Womens Health. 2003;48(2):96–104; doi: 10.1016/s1526-9523(02)00416-6.

23. Bilardi JE, Walker S, Temple-Smith M, McNair R, Mooney-Somers J, Bellhouse C, et al. The burden of bacterial vaginosis: women’s experience of the physical, emotional, sexual and social impact of living with recurrent bacterial vaginosis. PLoS One. 2013;8(9):e74378; doi: 10.1371/journal.pone.0074378.

24. Klebanoff MA, Schwebke JR, Zhang J, Nansel TR, Yu KF, Andrews WW. Vulvovaginal symptoms in women with bacterial vaginosis. Obstet Gynecol. 2004;104(2):267–72; doi: 10.1097/01.AOG.0000134783.98382.b0.

25. Nugent RP, Krohn MA, Hillier SL. Reliability of diagnosing bacterial vaginosis is improved by a standardized method of gram stain interpretation. Journal of clinical microbiology. 1991;29(2):297–301.

26. McKinnon LR, Achilles SL, Bradshaw CS, Burgener A, Crucitti T, Fredricks DN, et al. The Evolving Facets of Bacterial Vaginosis: Implications for HIV Transmission. AIDS Res Hum Retroviruses. 2019;35(3):219–28; doi: 10.1089/AID.2018.0304.

27. Sela U, Euler CW, Correa da Rosa J, Fischetti VA. Strains of bacterial species induce a greatly varied acute adaptive immune response: The contribution of the accessory genome. PLoS Pathog. 2018;14(1):e1006726; doi: 10.1371/journal.ppat.1006726.

28. Douillard FP, Ribbera A, Kant R, Pietila TE, Jarvinen HM, Messing M, et al. Comparative genomic and functional analysis of 100 Lactobacillus rhamnosus strains and their comparison with strain GG. PLoS Genet. 2013;9(8):e1003683; doi: 10.1371/journal.pgen.1003683.

29. Ma B, France MT, Crabtree J, Holm JB, Humphrys MS, Brotman RM, et al. A comprehensive non-redundant gene catalog reveals extensive within-community intraspecies diversity in the human vagina. Nat Commun. 2020;11(1):940; doi: 10.1038/s41467-020-14677-3.

30. Tortelli BA, Lewis AL, Fay JC. The structure and diversity of strain-level variation in vaginal bacteria. Microb Genom. 2021;7(3); doi: 10.1099/mgen.0.000543.

31. O’Hanlon DE, Moench TR, Cone RA. Vaginal pH and microbicidal lactic acid when lactobacilli dominate the microbiota. PLoS One. 2013;8(11):e80074; doi: 10.1371/journal.pone.0080074.

32. Tachedjian G, Aldunate M, Bradshaw CS, Cone RA. The role of lactic acid production by probiotic Lactobacillus species in vaginal health. Res Microbiol. 2017;168(9-10):782–92; doi: 10.1016/j.resmic.2017.04.001.

33. Kochhar S, Hottinger H, Chuard N, Taylor PG, Atkinson T, Scawen MD, et al. Cloning and overexpression of Lactobacillus helveticus D-lactate dehydrogenase gene in Escherichia coli. Eur J Biochem. 1992;208(3):799–805; doi: 10.1111/j.1432-1033.1992.tb17250.x.

34. Petrova MI, Reid G, Vaneechoutte M, Lebeer S. Lactobacillus iners: Friend or Foe? Trends Microbiol. 2017;25(3):182–91; doi: 10.1016/j.tim.2016.11.007.

35. Whelan F, Lafita A, Gilburt J, Degut C, Griffiths SC, Jenkins HT, et al. Periscope Proteins are variable-length regulators of bacterial cell surface interactions. Proc Natl Acad Sci U S A. 2021;118(23); doi: 10.1073/pnas.2101349118.

36. Gaillard JL, Berche P, Frehel C, Gouin E, Cossart P. Entry of L. monocytogenes into cells is mediated by internalin, a repeat protein reminiscent of surface antigens from gram-positive cocci. Cell. 1991;65(7):1127–41; doi: 10.1016/0092-8674(91)90009-n.

37. Rampersaud R, Planet PJ, Randis TM, Kulkarni R, Aguilar JL, Lehrer RI, et al. Inerolysin, a cholesterol-dependent cytolysin produced by Lactobacillus iners. J Bacteriol. 2011;193(5):1034–41; doi: 10.1128/JB.00694-10.

38. Vaneechoutte M, Guschin A, Van Simaey L, Gansemans Y, Van Nieuwerburgh F, Cools P. Emended description of Gardnerella vaginalis and description of Gardnerella leopoldii sp. nov., Gardnerella piotii sp. nov. and Gardnerella swidsinskii sp. nov., with delineation of 13 genomic species within the genus Gardnerella. Int J Syst Evol Microbiol. 2019;69(3):679–87; doi: 10.1099/ijsem.0.003200.

39. Muzny CA, Blanchard E, Taylor CM, Aaron KJ, Talluri R, Griswold ME, et al. Identification of Key Bacteria Involved in the Induction of Incident Bacterial Vaginosis: A Prospective Study. J Infect Dis. 2018;218(6):966–78; doi: 10.1093/infdis/jiy243.

40. Verstraelen H, Verhelst R, Claeys G, De Backer E, Temmerman M, Vaneechoutte M. Longitudinal analysis of the vaginal microflora in pregnancy suggests that L. crispatus promotes the stability of the normal vaginal microflora and that L. gasseri and/or L. iners are more conducive to the occurrence of abnormal vaginal microflora. BMC Microbiol. 2009;9:116; doi: 10.1186/1471-2180-9-116.

41. Ravel J, Brotman RM, Gajer P, Ma B, Nandy M, Fadrosh DW, et al.: Daily temporal dynamics of vaginal microbiota before, during and after episodes of bacterial vaginosis. In: Microbiome. vol. 1: BioMed Central; 2013: 29.

42. Ferris MJ, Norori J, Zozaya-Hinchliffe M, Martin DH: Cultivation-Independent Analysis of Changes in Bacterial Vaginosis Flora Following Metronidazole Treatment. In: Journal of Clinical Microbiology. vol. 45; 2007: 1016–8.

43. Peebles K, Velloza J, Balkus JE, McClelland RS, Barnabas RV. High Global Burden and Costs of Bacterial Vaginosis: A Systematic Review and Meta-Analysis. Sex Transm Dis. 2019;46(5):304–11; doi: 10.1097/OLQ.0000000000000972.

44. Macklaim JM, Fernandes AD, Di Bella JM, Hammond J-A, Reid G, Gloor GB: Comparative meta-RNA-seq of the vaginal microbiota and differential expression by Lactobacillus iners in health and dysbiosis. In: Microbiome. vol. 1; 2013: 12.

45. Canova MJ, Molle V. Bacterial serine/threonine protein kinases in host-pathogen interactions. J Biol Chem. 2014;289(14):9473–9; doi: 10.1074/jbc.R113.529917.

46. Tamarelle J, Ma B, Gajer P, Humphrys MS, Terplan M, Mark KS, et al. Nonoptimal Vaginal Microbiota After Azithromycin Treatment for Chlamydia trachomatis Infection. J Infect Dis. 2020;221(4):627–35; doi: 10.1093/infdis/jiz499.

47. Potter RF, Burnham CD, Dantas G. In Silico Analysis of Gardnerella Genomospecies Detected in the Setting of Bacterial Vaginosis. Clin Chem. 2019;65(11):1375–87; doi: 10.1373/clinchem.2019.305474.

48. Ksiezarek M, Ugarcina-Perovic S, Rocha J, Grosso F, Peixe L. Long-term stability of the urogenital microbiota of asymptomatic European women. Bmc Microbiol. 2021;21(1):64; doi: 10.1186/s12866-021-02123-3.

49. Turner E, Sobel JD, Akins RA. Prognosis of recurrent bacterial vaginosis based on longitudinal changes in abundance of Lactobacillus and specific species of Gardnerella. PLoS One. 2021;16(8):e0256445; doi: 10.1371/journal.pone.0256445.

50. Zozaya-Hinchliffe M, Lillis R, Martin DH, Ferris MJ: Quantitative PCR Assessments of Bacterial Species in Women with and without Bacterial Vaginosis. In: Journal of Clinical Microbiology. vol. 48; 2010: 1812–9.

51. Janulaitiene M, Paliulyte V, Grinceviciene S, Zakareviciene J, Vladisauskiene A, Marcinkute A, et al. Prevalence and distribution of Gardnerella vaginalis subgroups in women with and without bacterial vaginosis. BMC Infect Dis. 2017;17(1):394; doi: 10.1186/s12879-017-2501-y.

52. Gelber SE, Aguilar JL, Lewis KL, Ratner AJ. Functional and phylogenetic characterization of Vaginolysin, the human-specific cytolysin from Gardnerella vaginalis. J Bacteriol. 2008;190(11):3896–903; doi: 10.1128/JB.01965-07.

53. Pleckaityte M, Janulaitiene M, Lasickiene R, Zvirbliene A. Genetic and biochemical diversity of Gardnerella vaginalis strains isolated from women with bacterial vaginosis. FEMS Immunol Med Microbiol. 2012;65(1):69–77; doi: 10.1111/j.1574-695X.2012.00940.x.

54. Yeoman CJ, Yildirim S, Thomas SM, Durkin AS, Torralba M, Sutton G, et al. Comparative genomics of Gardnerella vaginalis strains reveals substantial differences in metabolic and virulence potential. PLoS One. 2010;5(8):e12411; doi: 10.1371/journal.pone.0012411.

55. Koumans EH, Sternberg M, Bruce C, McQuillan G, Kendrick J, Sutton M, et al. The prevalence of bacterial vaginosis in the United States, 2001-2004; associations with symptoms, sexual behaviors, and reproductive health. Sex Transm Dis. 2007;34(11):864–9; doi: 10.1097/OLQ.0b013e318074e565.

56. Workowski KA, Bachmann LH, Chan PA, Johnston CM, Muzny CA, Park I, et al. Sexually Transmitted Infections Treatment Guidelines, 2021. MMWR Recomm Rep. 2021;70(4):1–187; doi: 10.15585/mmwr.rr7004a1.

57. Brotman RM, Ravel J, Cone RA, Zenilman JM. Rapid fluctuation of the vaginal microbiota measured by Gram stain analysis. Sex Transm Infect. 2010;86(4):297–302; doi: 10.1136/sti.2009.040592.

58. Gajer P, Brotman RM, Bai G, Sakamoto J, Schütte UME, Zhong X, et al.: Temporal Dynamics of the Human Vaginal Microbiota. In: Sci Transl Med. vol. 4: American Association for the Advancement of Science; 2012: 132ra52–ra52.

59. Munoz A, Hayward MR, Bloom SM, Rocafort M, Ngcapu S, Mafunda NA, et al. Modeling the temporal dynamics of cervicovaginal microbiota identifies targets that may promote reproductive health. Microbiome. 2021;9(1):163; doi: 10.1186/s40168-021-01096-9.

60. Gevers D, Knight R, Petrosino JF, Huang K, McGuire AL, Birren BW, et al. The Human Microbiome Project: a community resource for the healthy human microbiome. 2012.

61. Li F, Chen C, Wei W, Wang Z, Dai J, Hao L, et al. The metagenome of the female upper reproductive tract. Gigascience. 2018;7(10); doi: 10.1093/gigascience/giy107.

62. Bolger AM, Lohse M, Usadel B. Trimmomatic: a flexible trimmer for Illumina sequence data. Bioinformatics. 2014;30(15):2114–20; doi: 10.1093/bioinformatics/btu170.

63. Parks DH, Chuvochina M, Waite DW, Rinke C, Skarshewski A, Chaumeil PA, et al. A standardized bacterial taxonomy based on genome phylogeny substantially revises the tree of life. Nat Biotechnol. 2018;36(10):996–1004; doi: 10.1038/nbt.4229.

64. Oksanen J, Kindt R, Legendre P, O’Hara B, Stevens MHH, Oksanen MJ, et al. The vegan package. Community ecology package. 2007;10:631–7.

65. Langfelder P, Zhang B, Horvath S. Defining clusters from a hierarchical cluster tree: the Dynamic Tree Cut package for R. Bioinformatics. 2008;24(5):719–20; doi: 10.1093/bioinformatics/btm563.

66. Warnes MGR, Bolker B, Bonebakker L, Gentleman R. Package ‘gplots’. Various R Programming Tools for Plotting Data. 2016.

67. Yue JC, Clayton MK. A similarity measure based on species proportions. Communications in Statistics-theory and Methods. 2005;34(11):2123–31.

68. U.B. IHaK. Fast Unified Random Forests for Survival, Regression, and Classification (RF-SRC), R package. 2021.

