## Supplementary figures and images for "High-resolution functional description of vaginal microbiomes in health and disease"

### Supplemental Figure 1

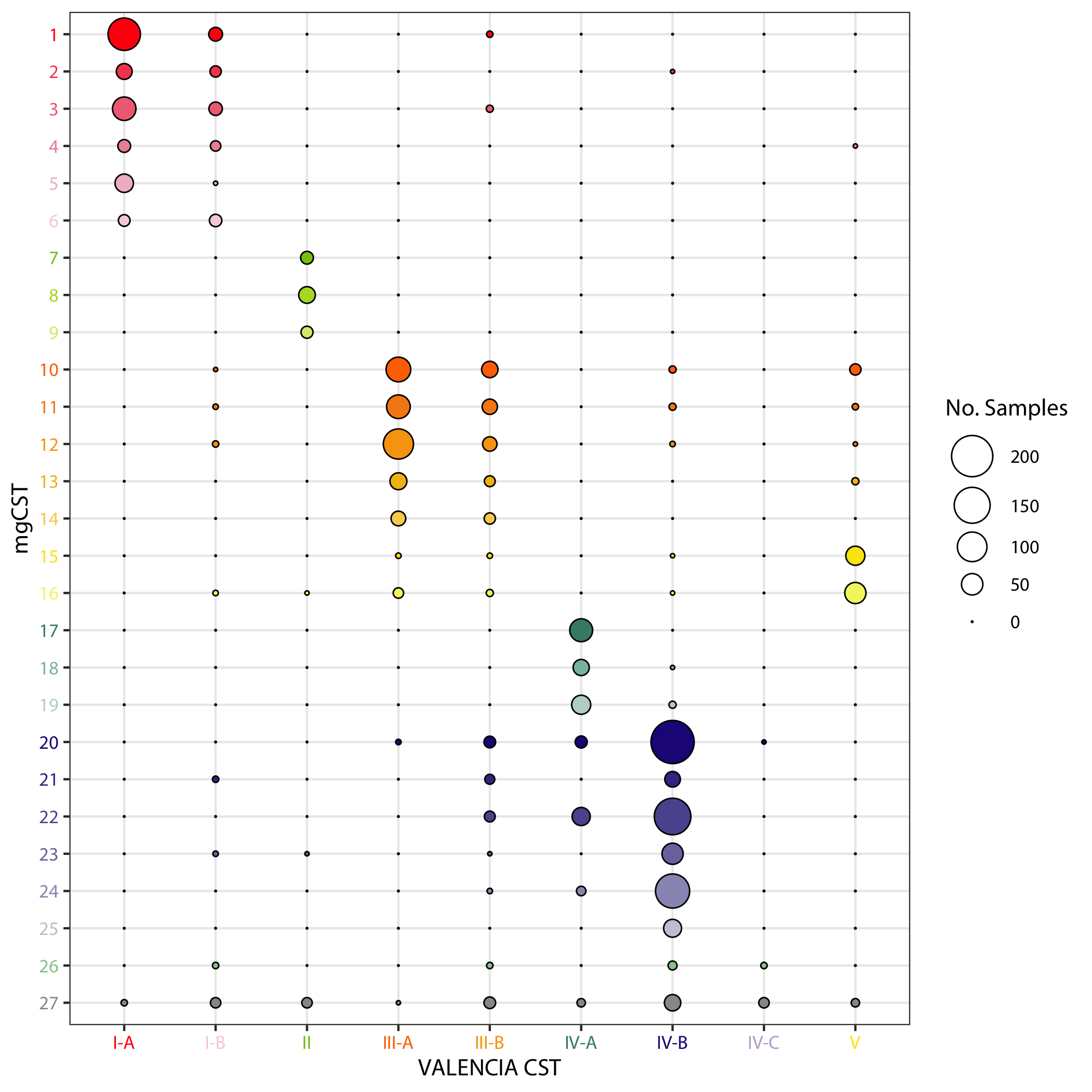

### Supplemental Figure 2

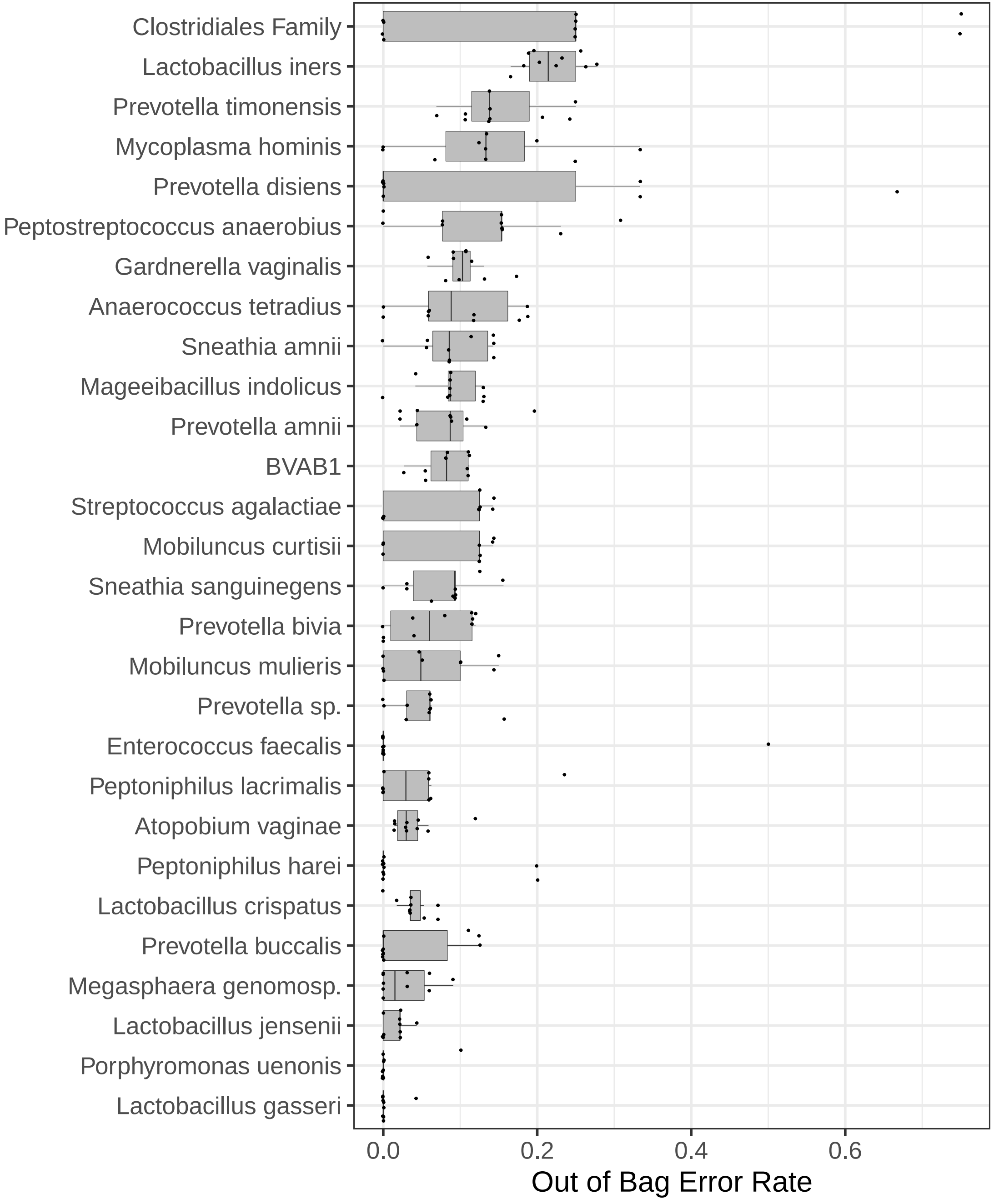

### Supplemental Figure 3

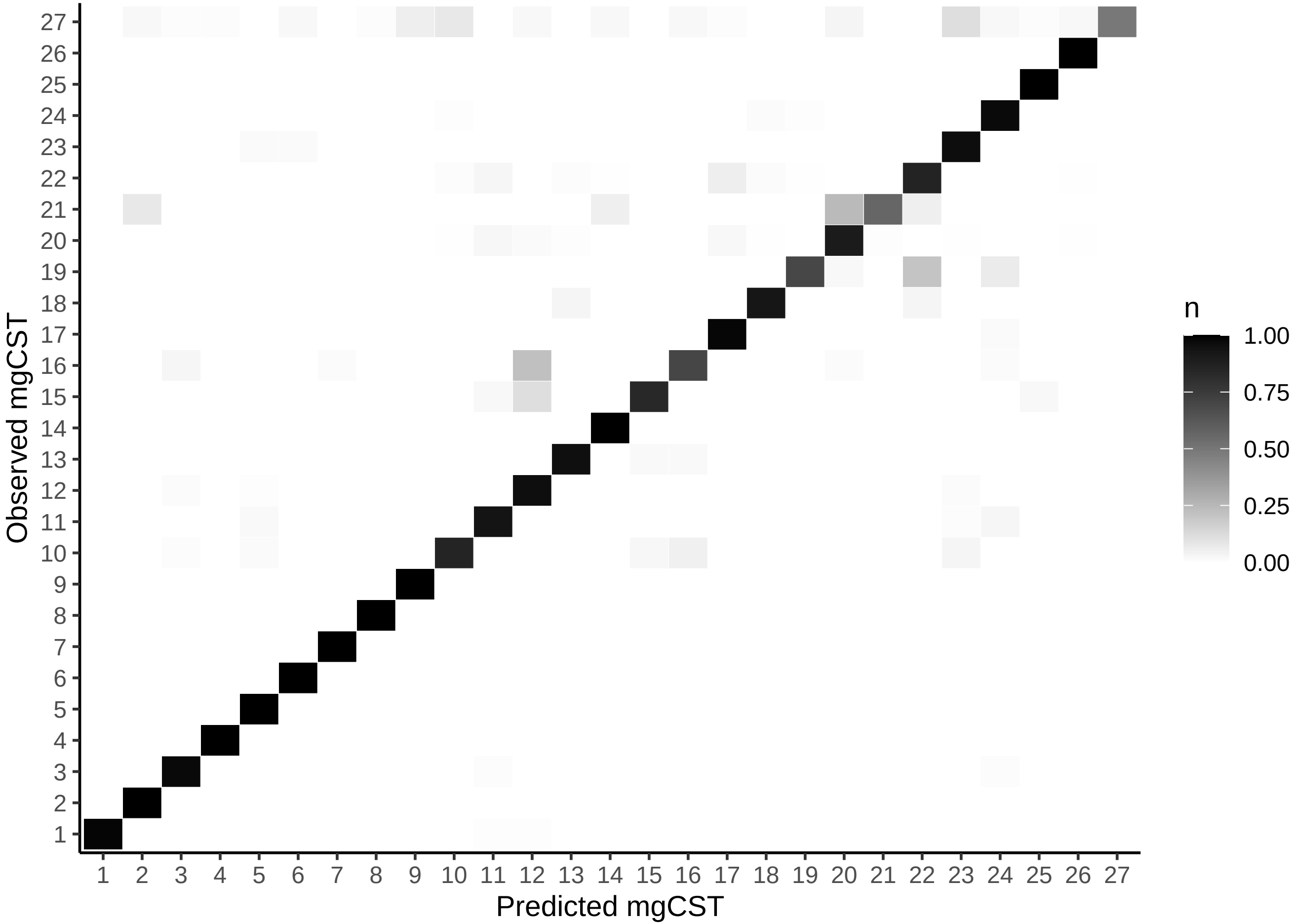
