## Supplemental File 4 for "High-resolution functional description of vaginal microbiomes in health and disease"

**Anaerococcus_tetradius**
**No. Samples=167 No. Genes=2619**

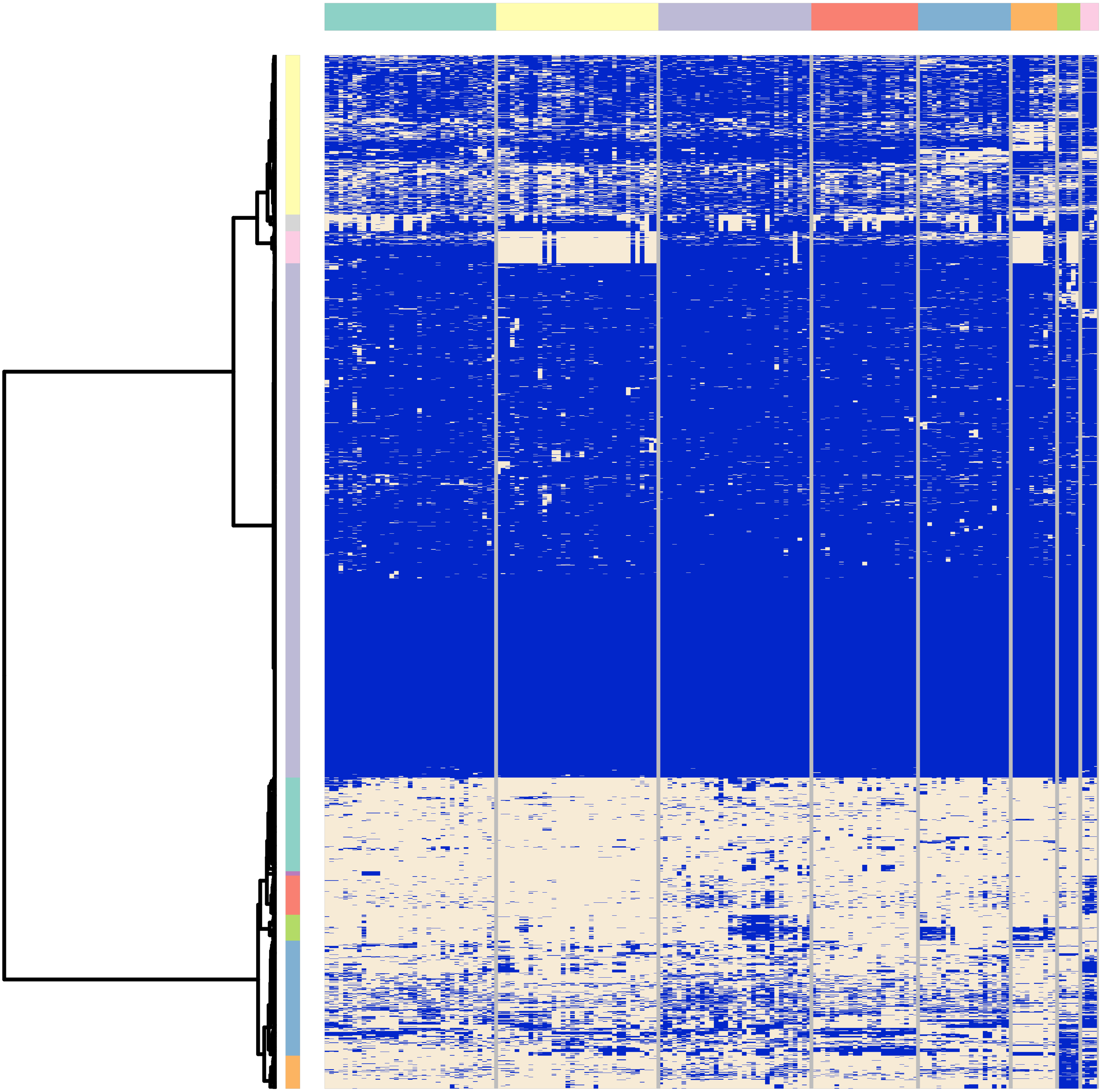

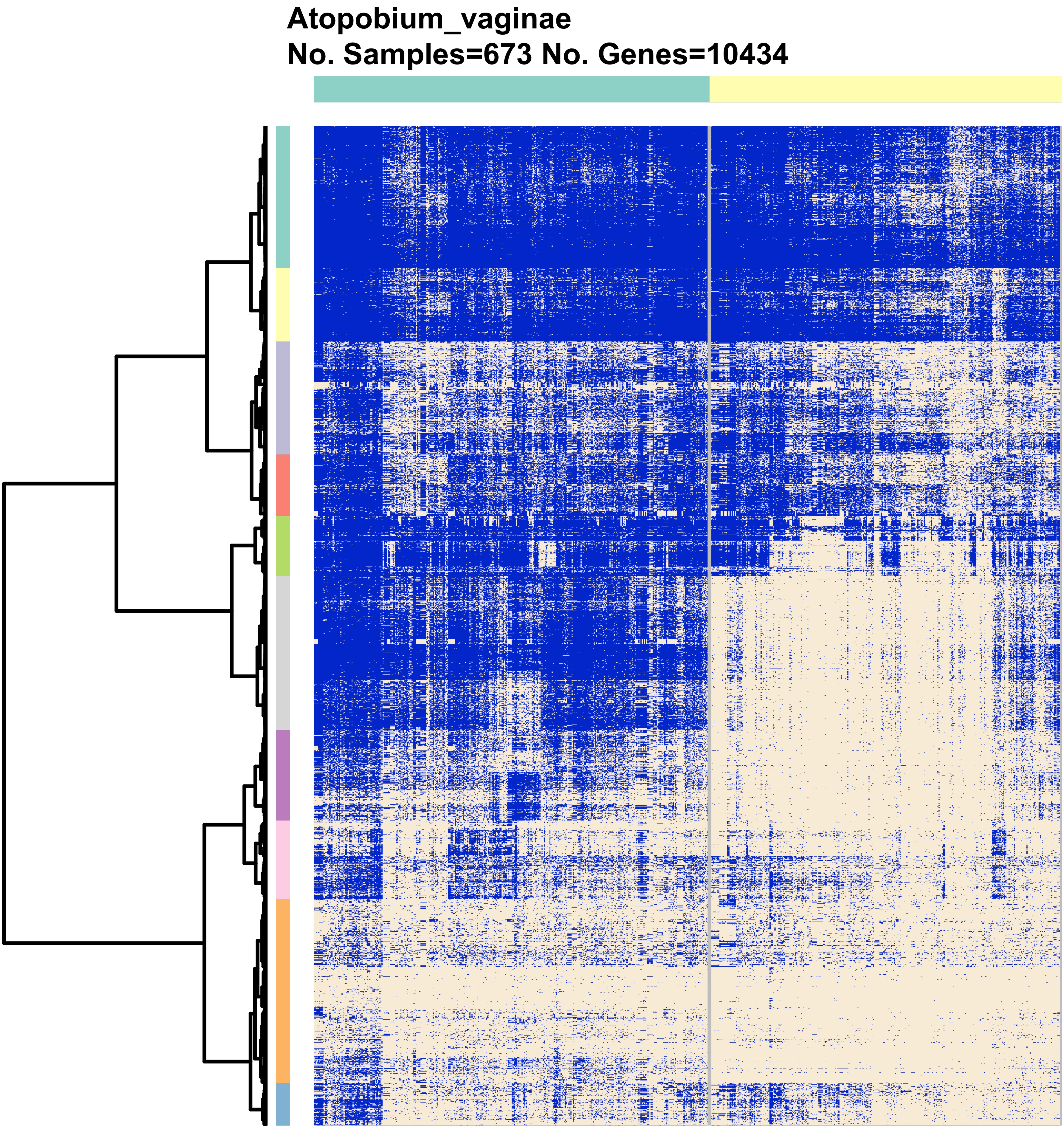
Atopobium_vaginae
No. Samples=673 No. Genes=10434

**BVAB1**
**No. Samples=364 No. Genes=1356**

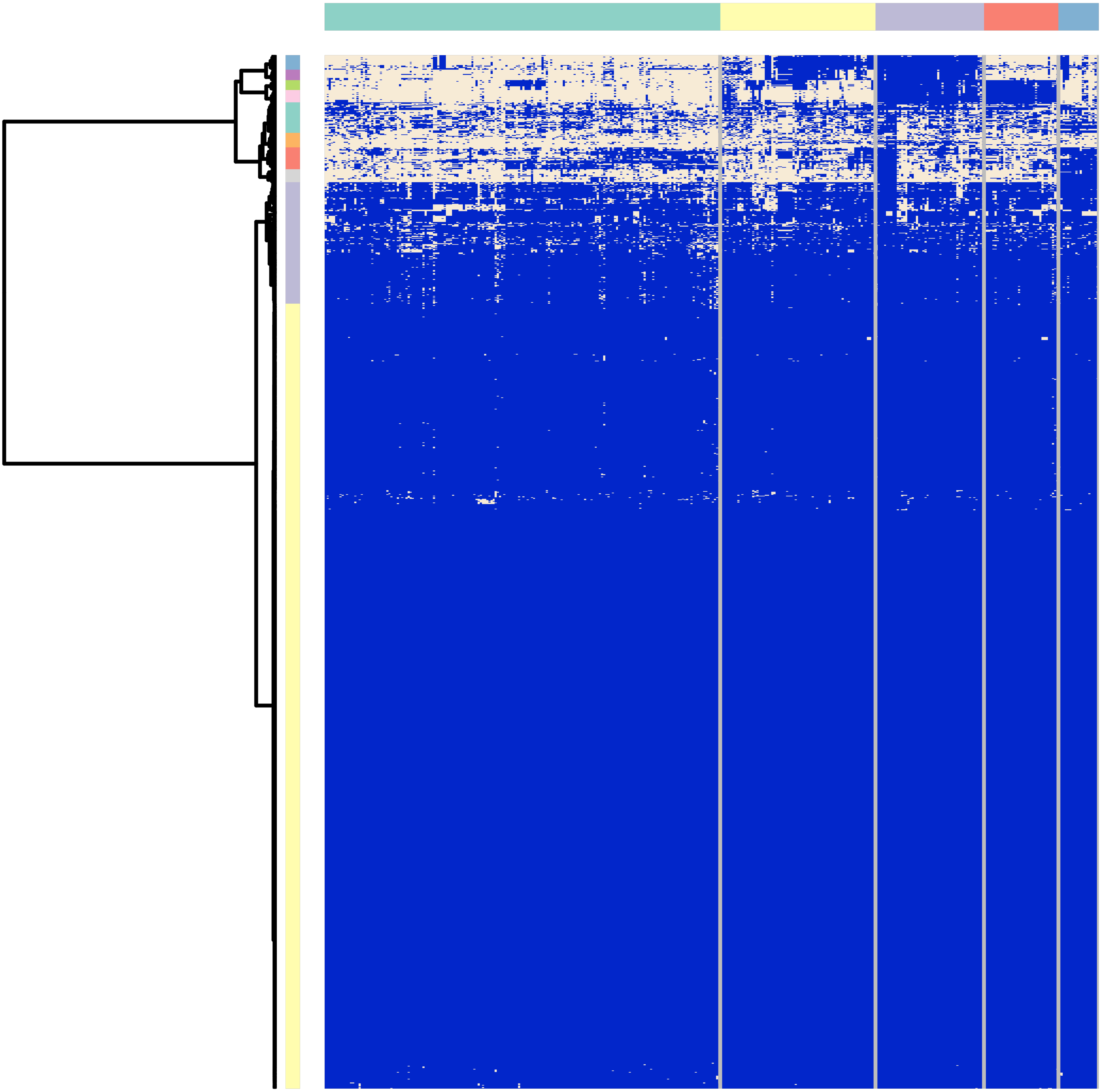

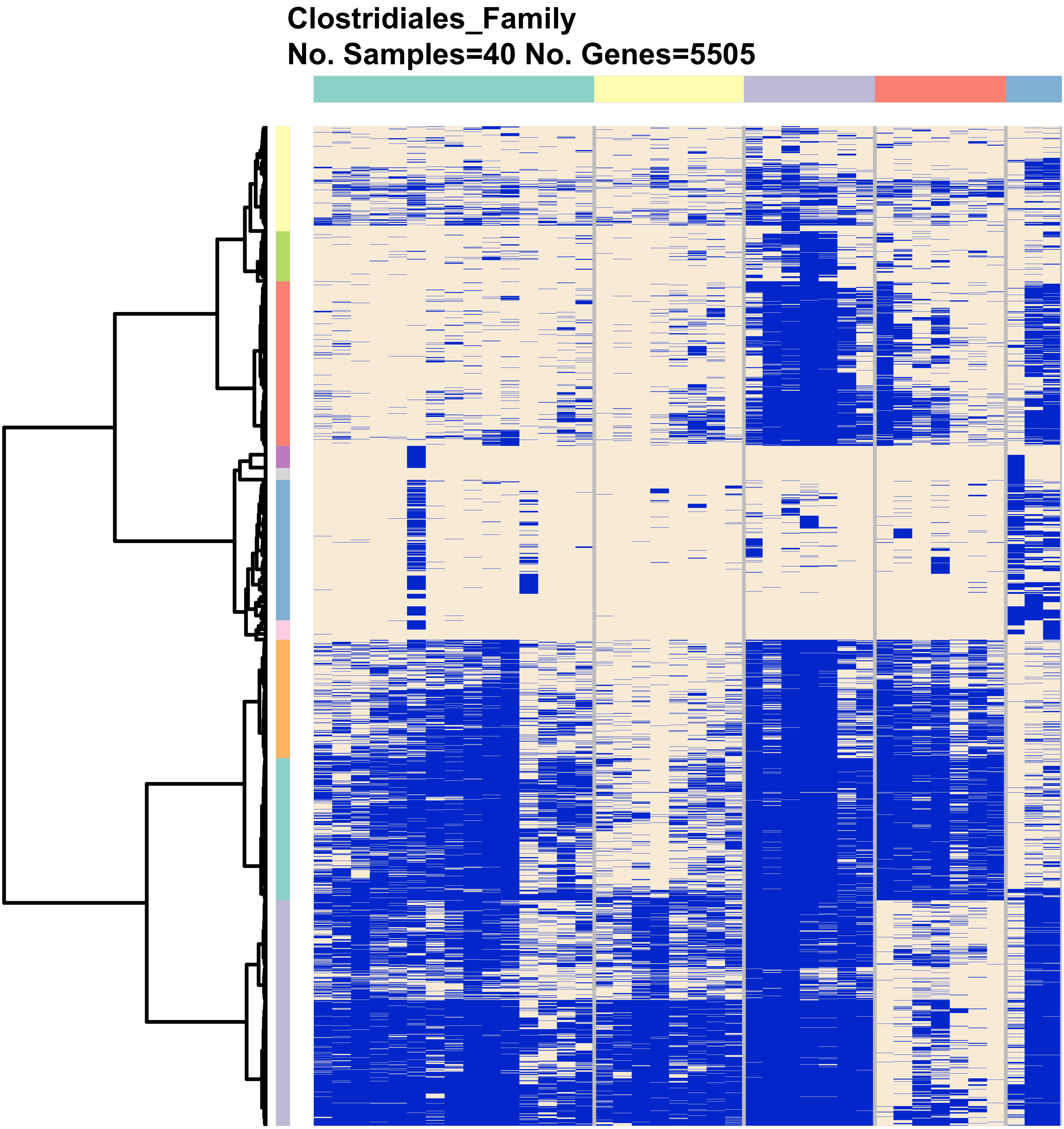
Clostridiales_Family
No. Samples=40 No. Genes=5505

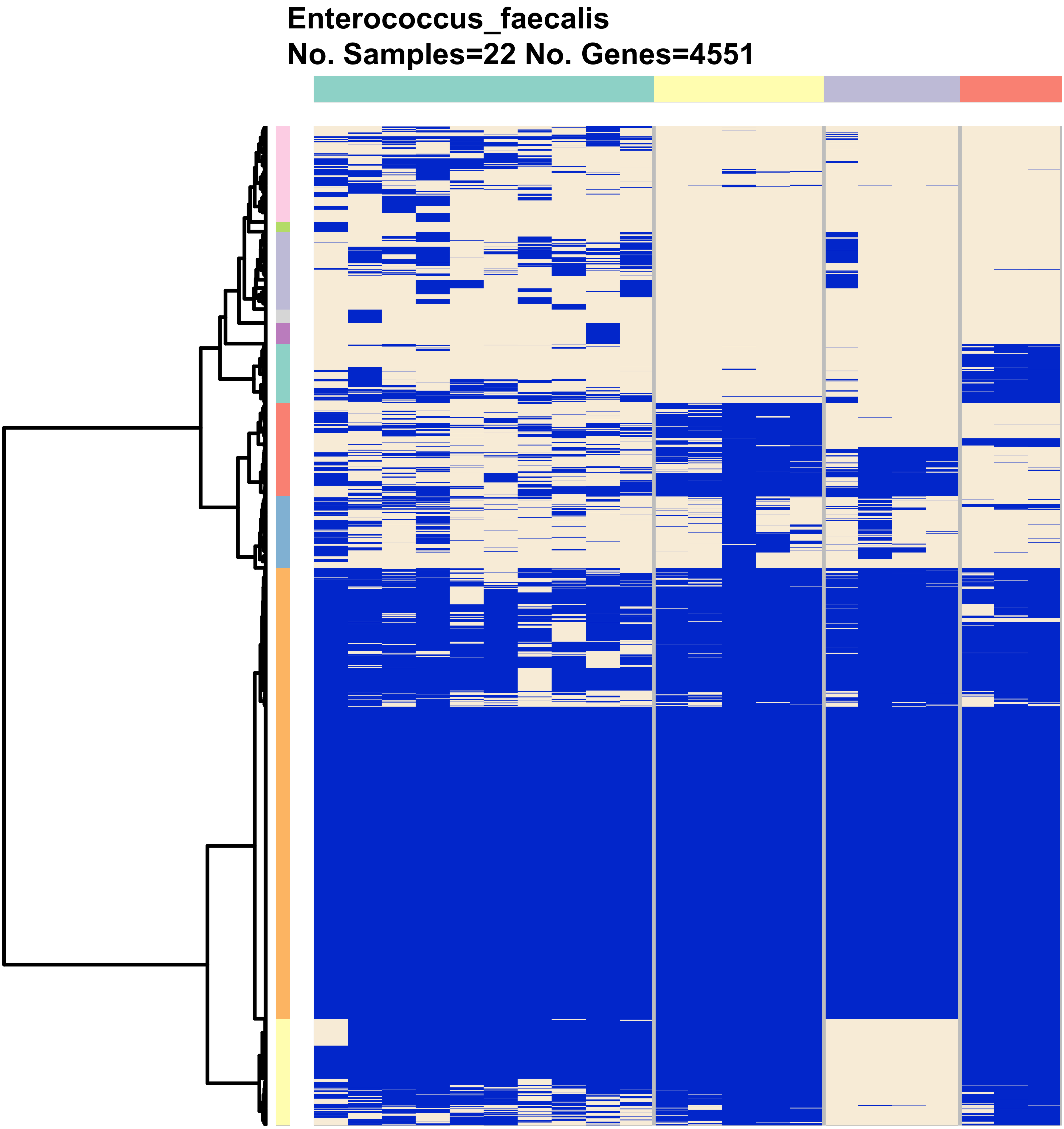
Enterococcus_faecalis
No. Samples=22 No. Genes=4551

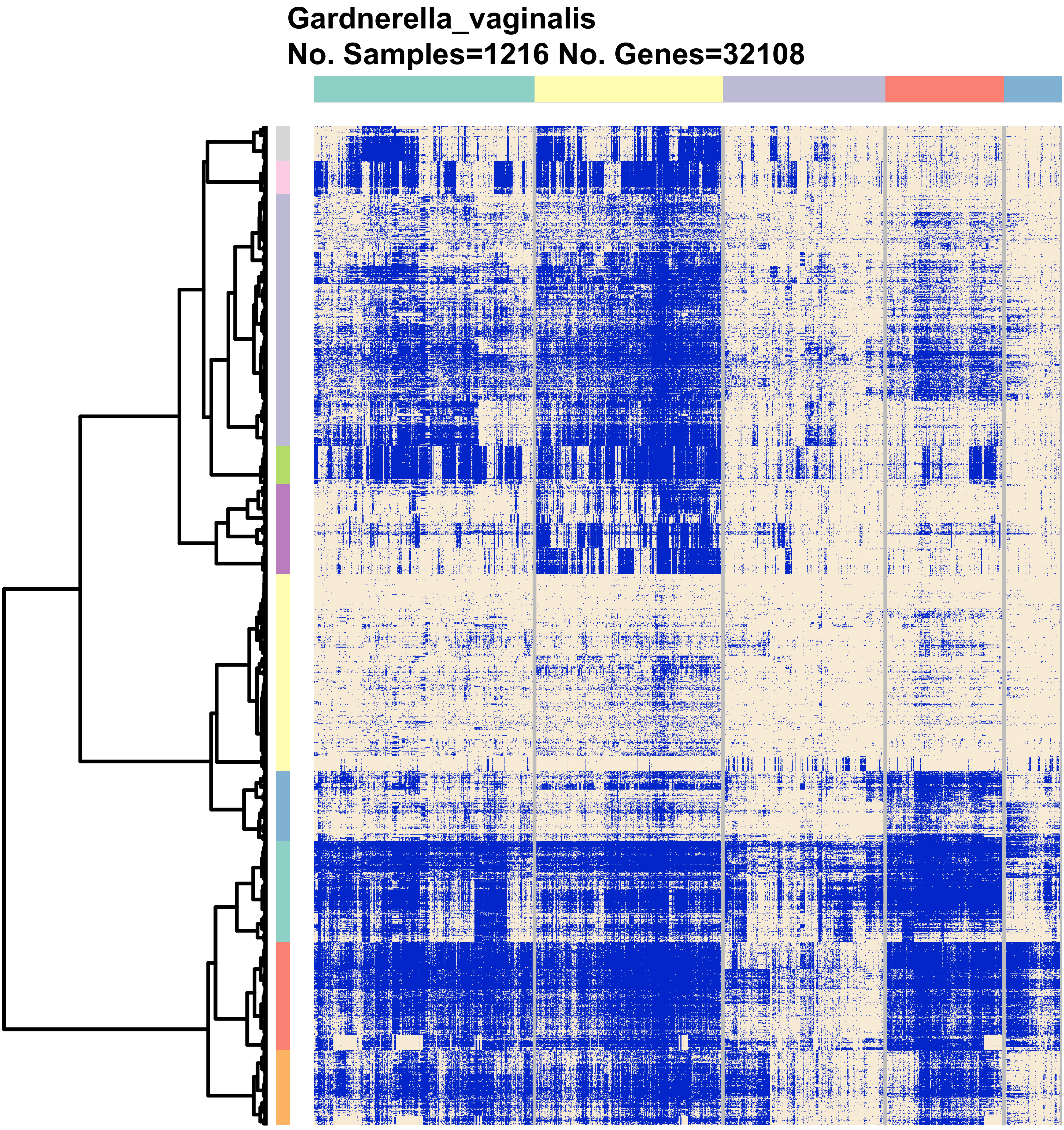
Gardnerella_vaginalis
No. Samples=1216 No. Genes=32108

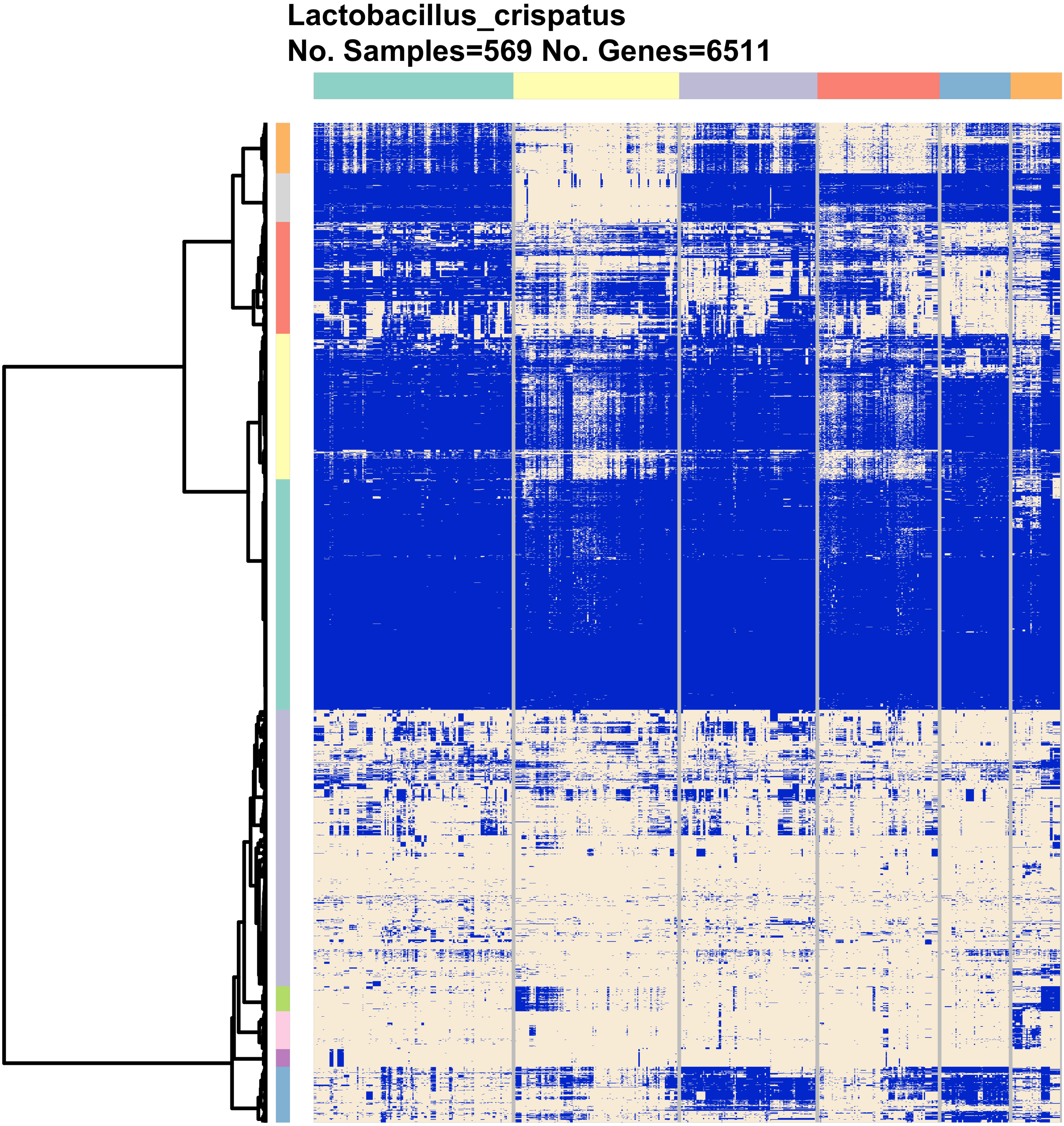
Lactobacillus_crispatus
No. Samples=569 No. Genes=6511

**Lactobacillus_gasseri**
**No. Samples=230 No. Genes=4476**

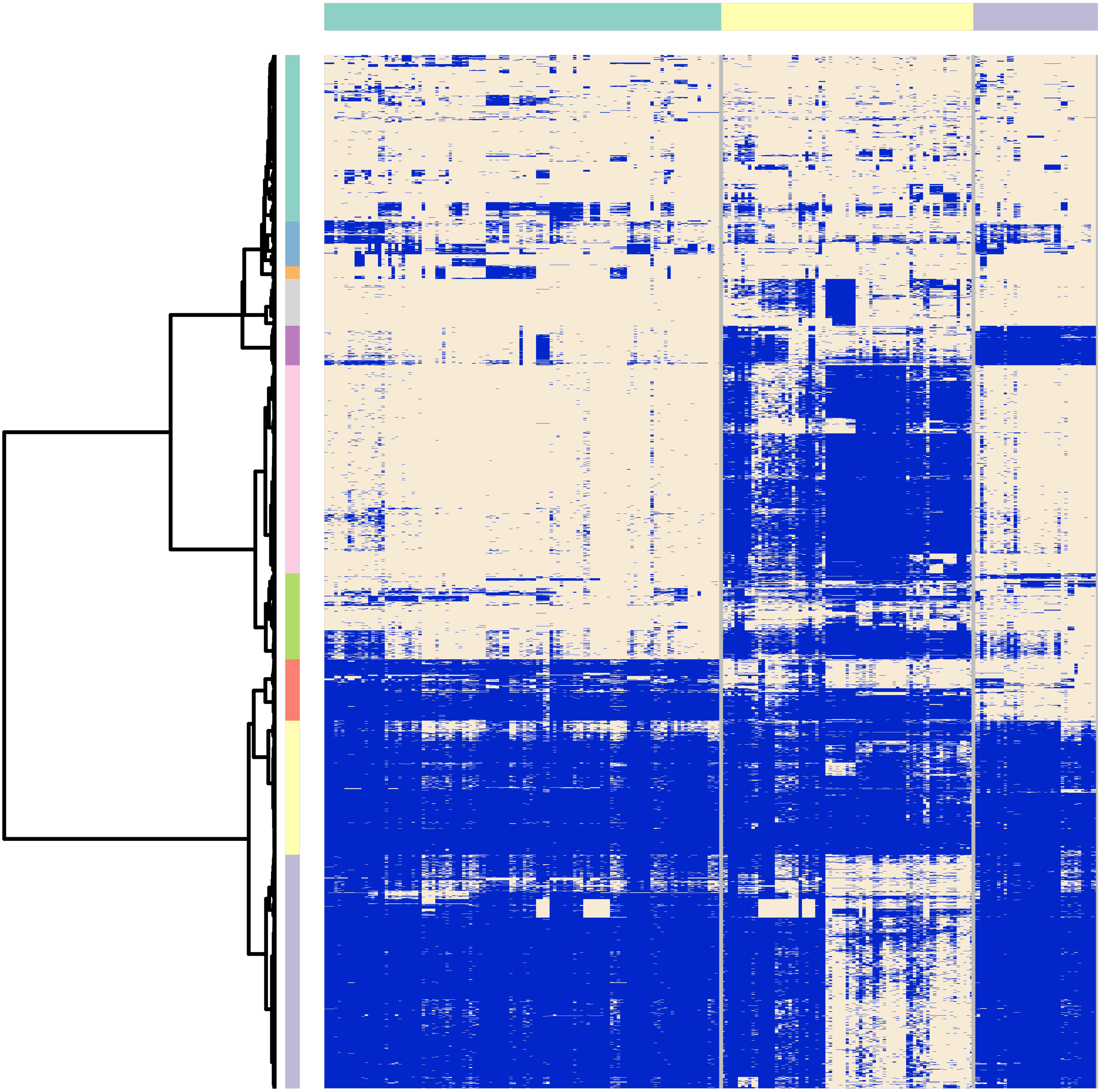

No. Samples= 1329 No. Genes=5027

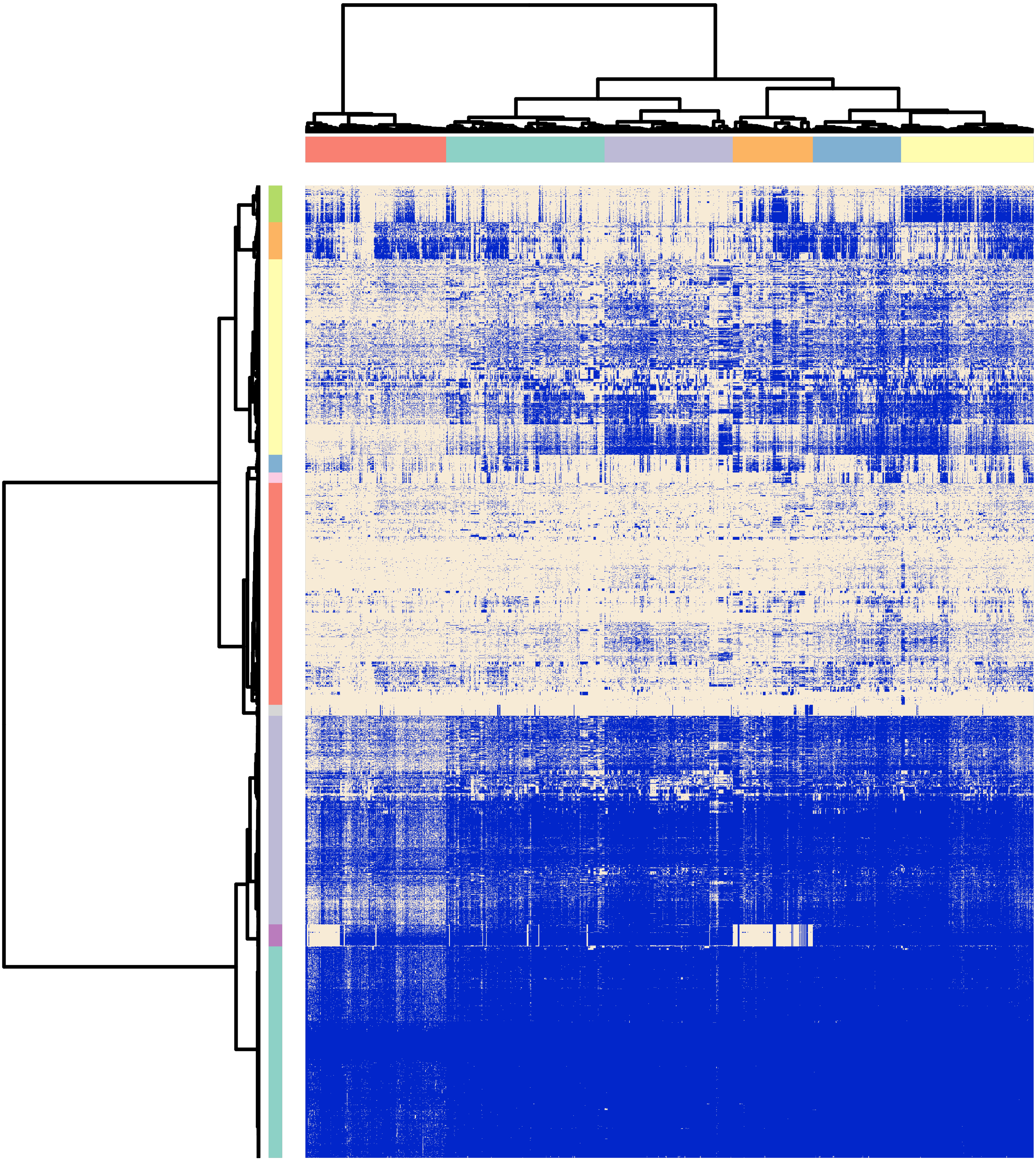

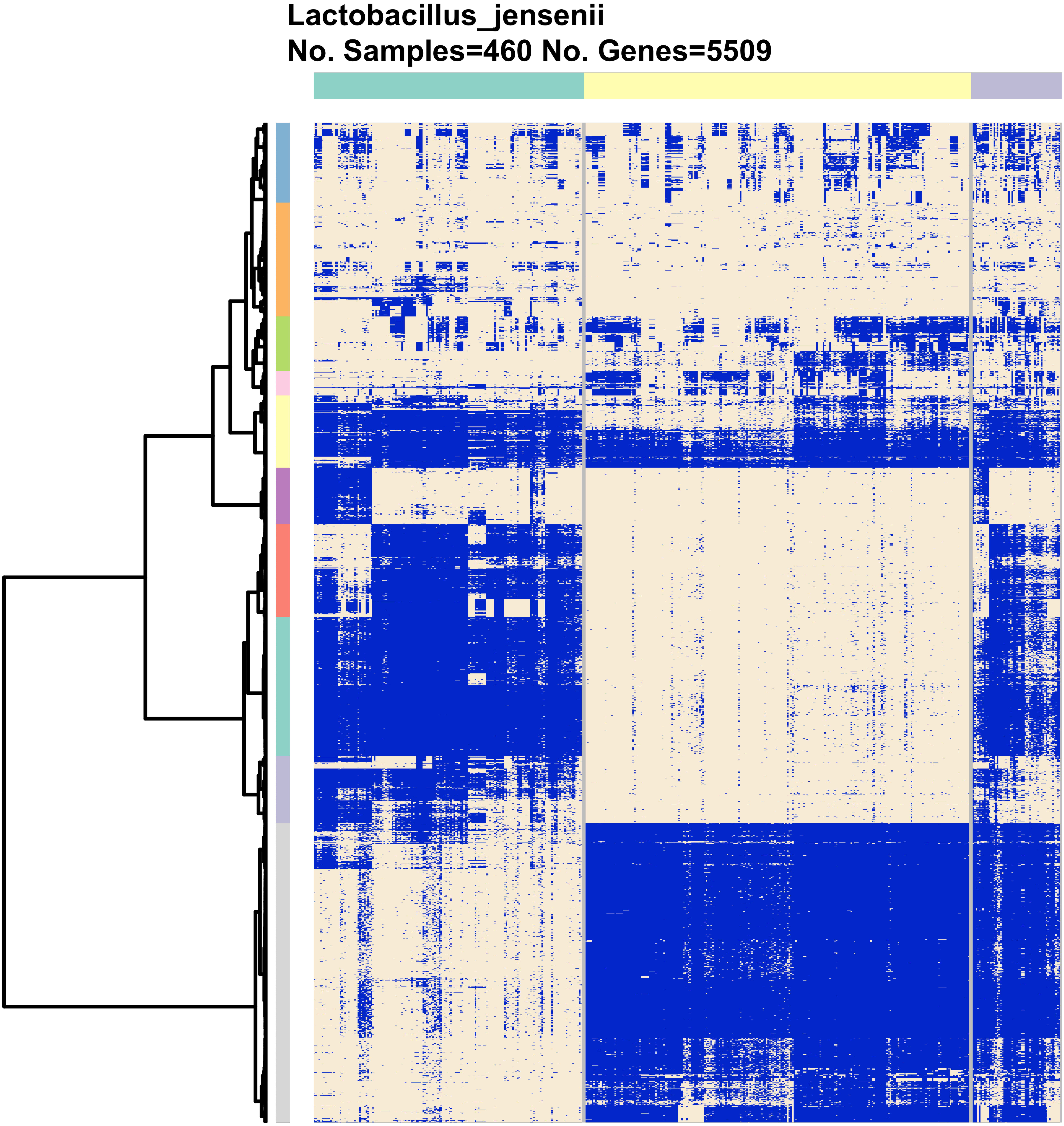
Lactobacillus_jensenii
No. Samples=460 No. Genes=5509

**Mageeibacillus_indolicus**
**No. Samples=232 No. Genes=2672**

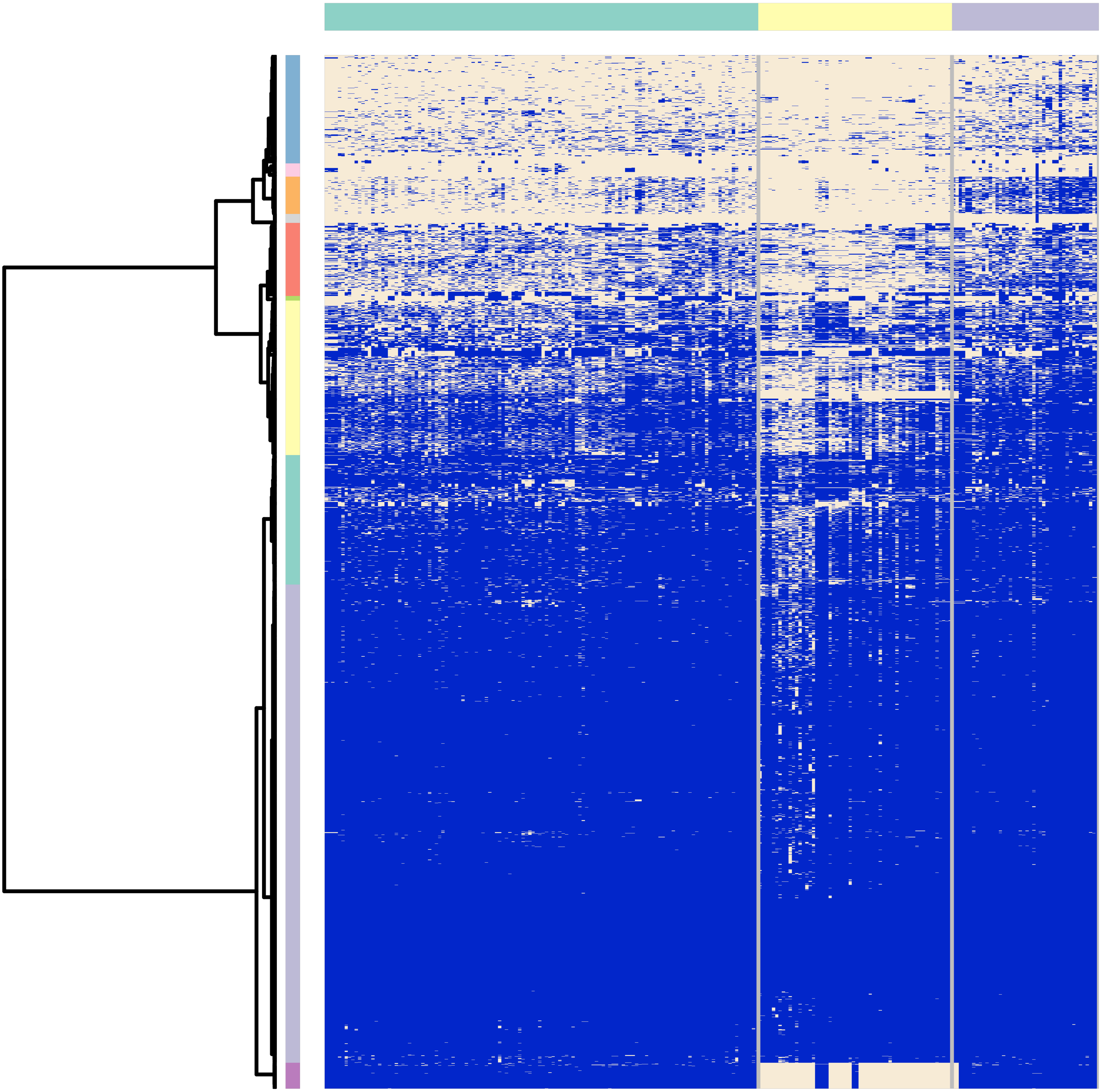

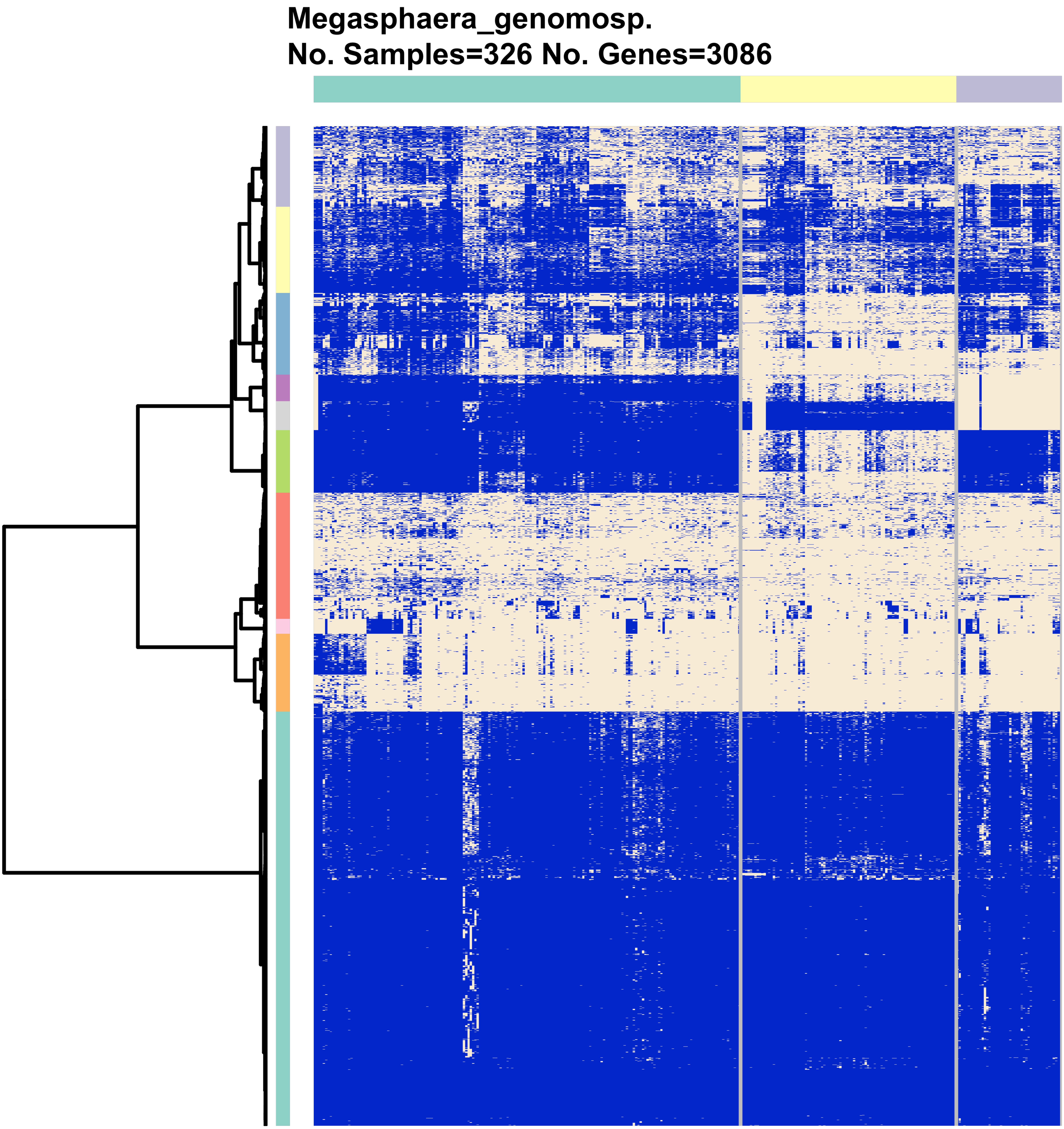
Megasphaera_genomosp.
No. Samples=326 No. Genes=3086

Mobiluncus_curtisii
No. Samples=78 No. Genes=2646

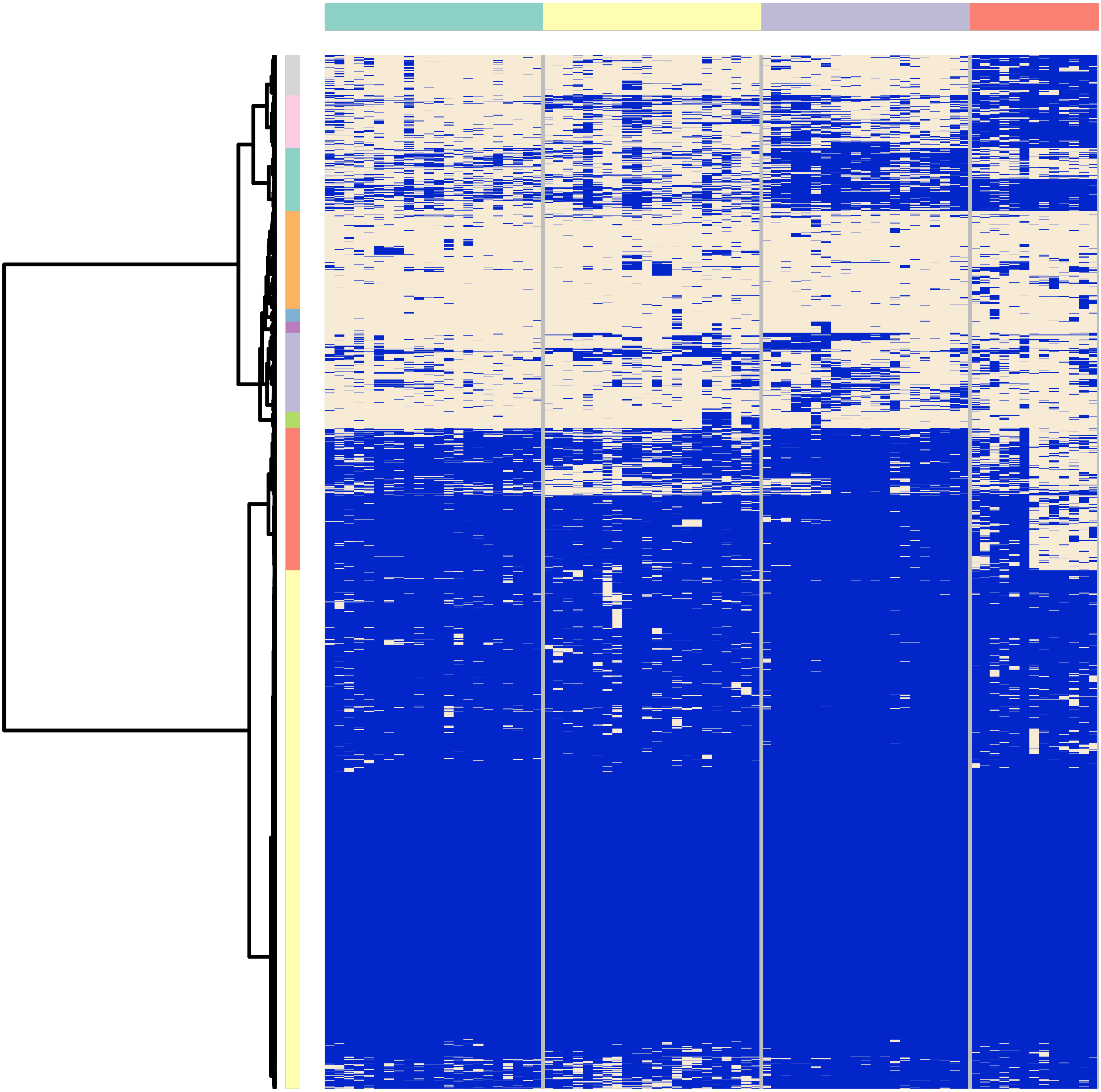

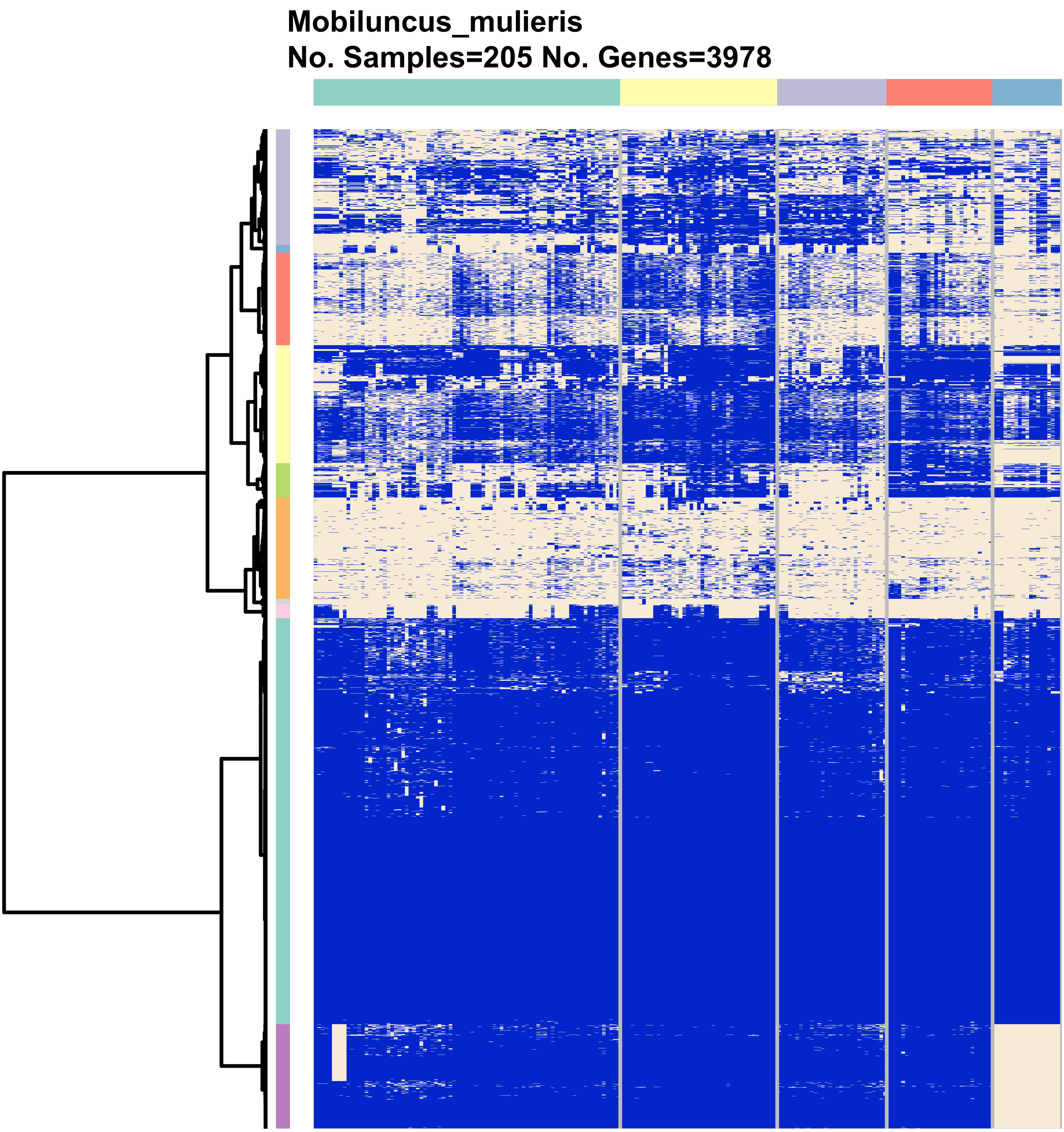

Mobiluncus_mulieris
No. Samples=205 No. Genes=3978

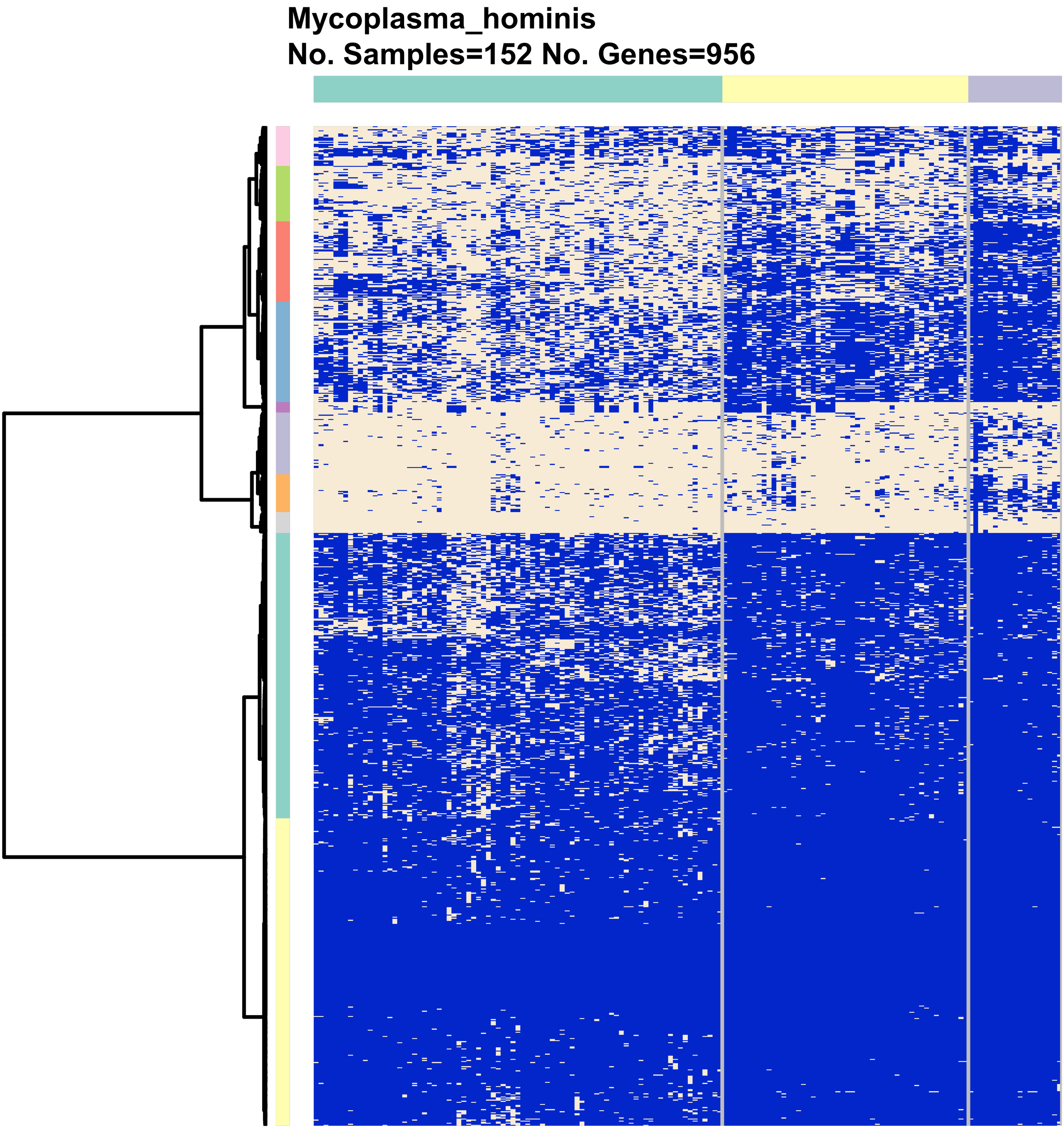
Mycoplasma_hominis
No. Samples=152 No. Genes=956

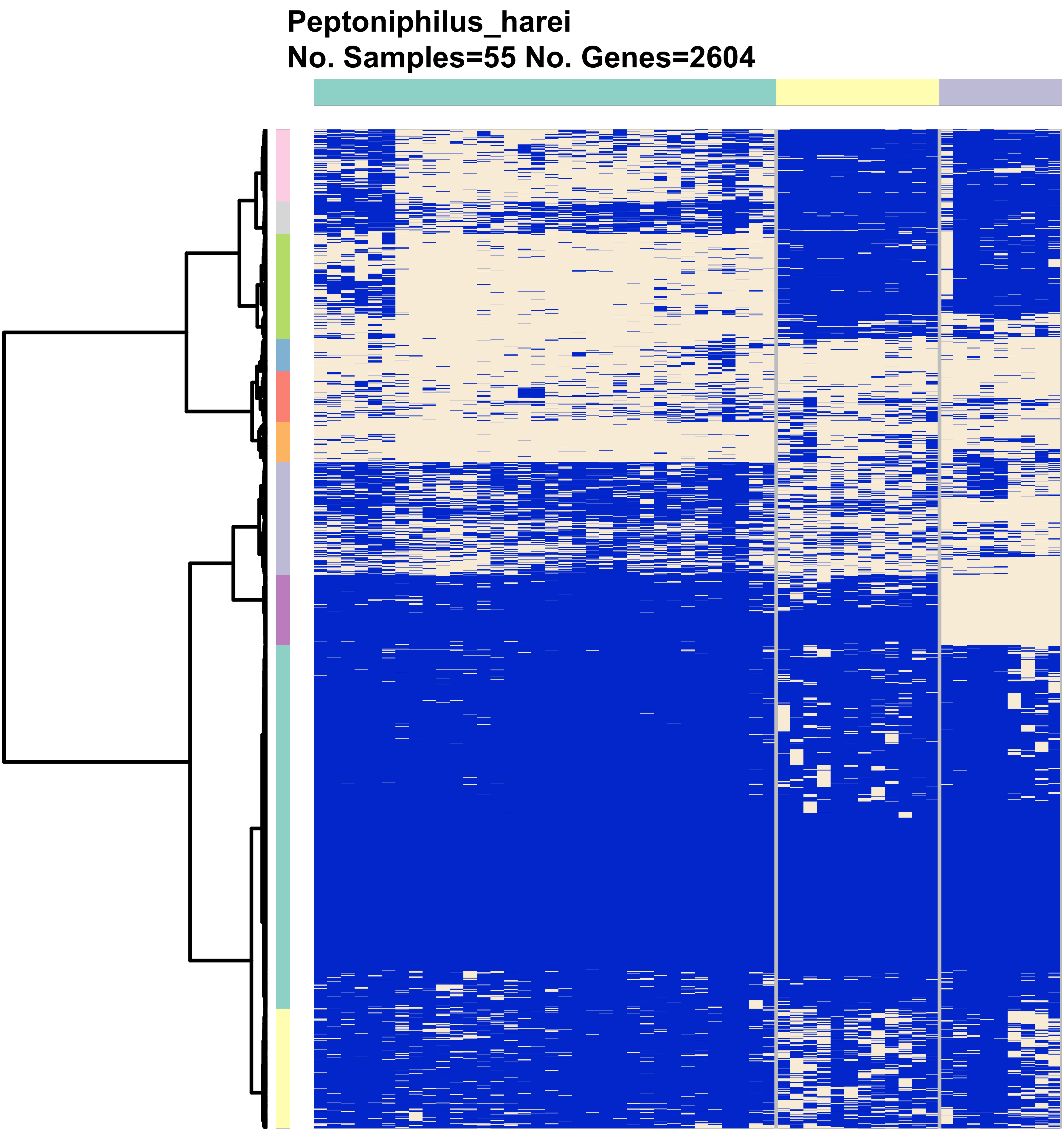
Peptoniphilus_harei
No. Samples=55 No. Genes=2604

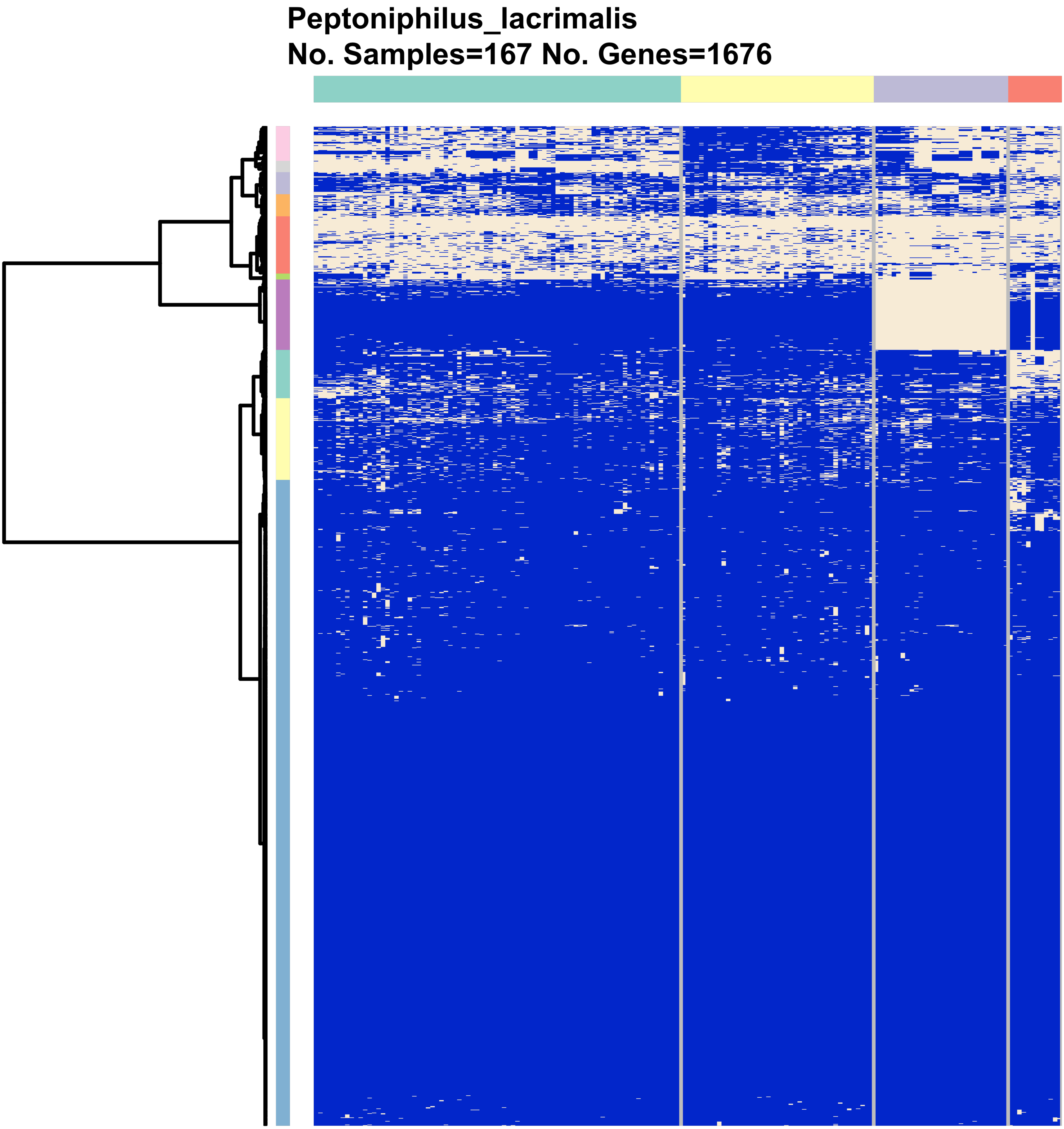
Peptoniphilus_lacrimalis
No. Samples=167 No. Genes=1676

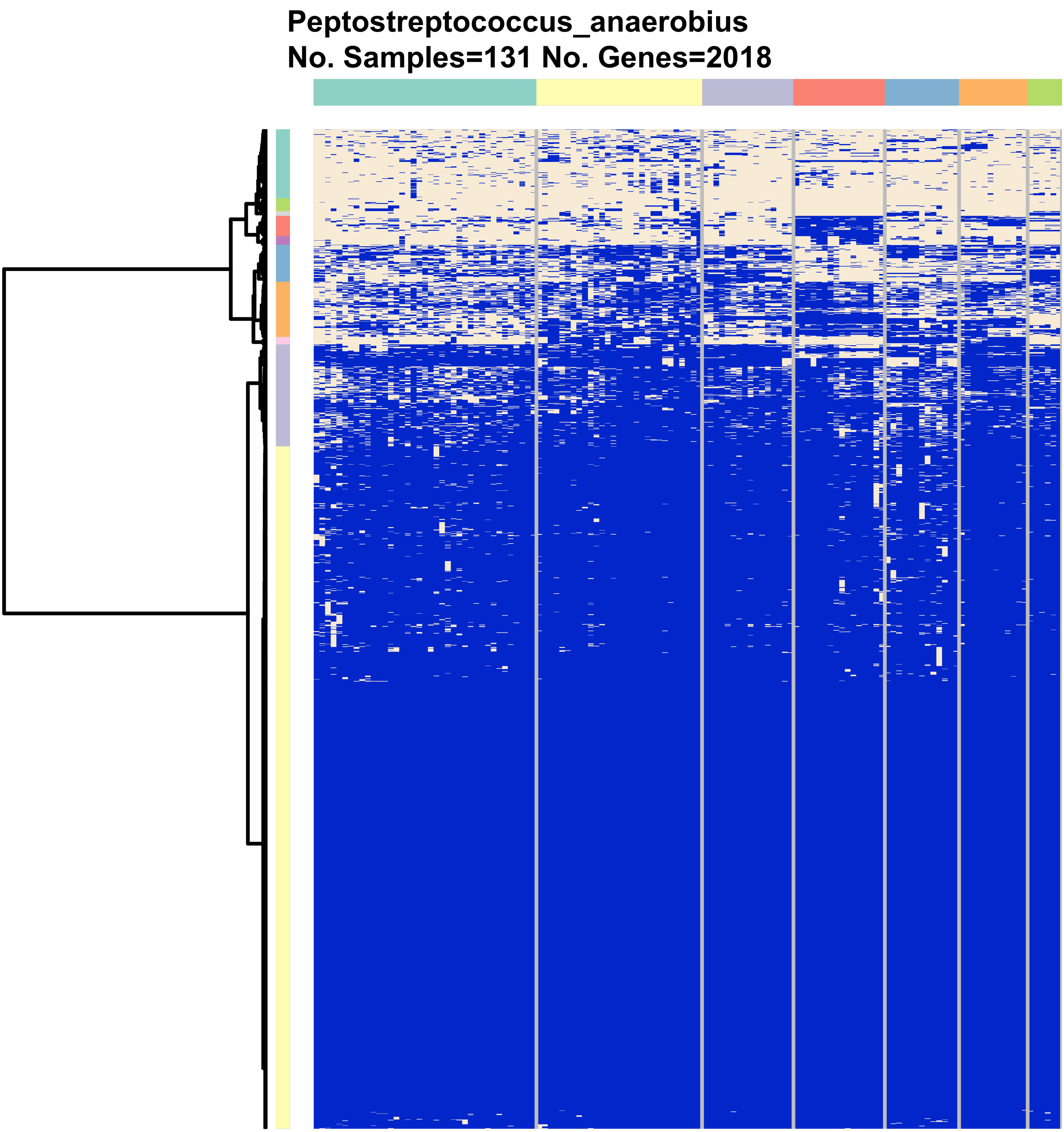
Peptostreptococcus_anaerobius
No. Samples=131 No. Genes=2018

**Porphyromonas_uenonis**
**No. Samples=100 No. Genes=2834**

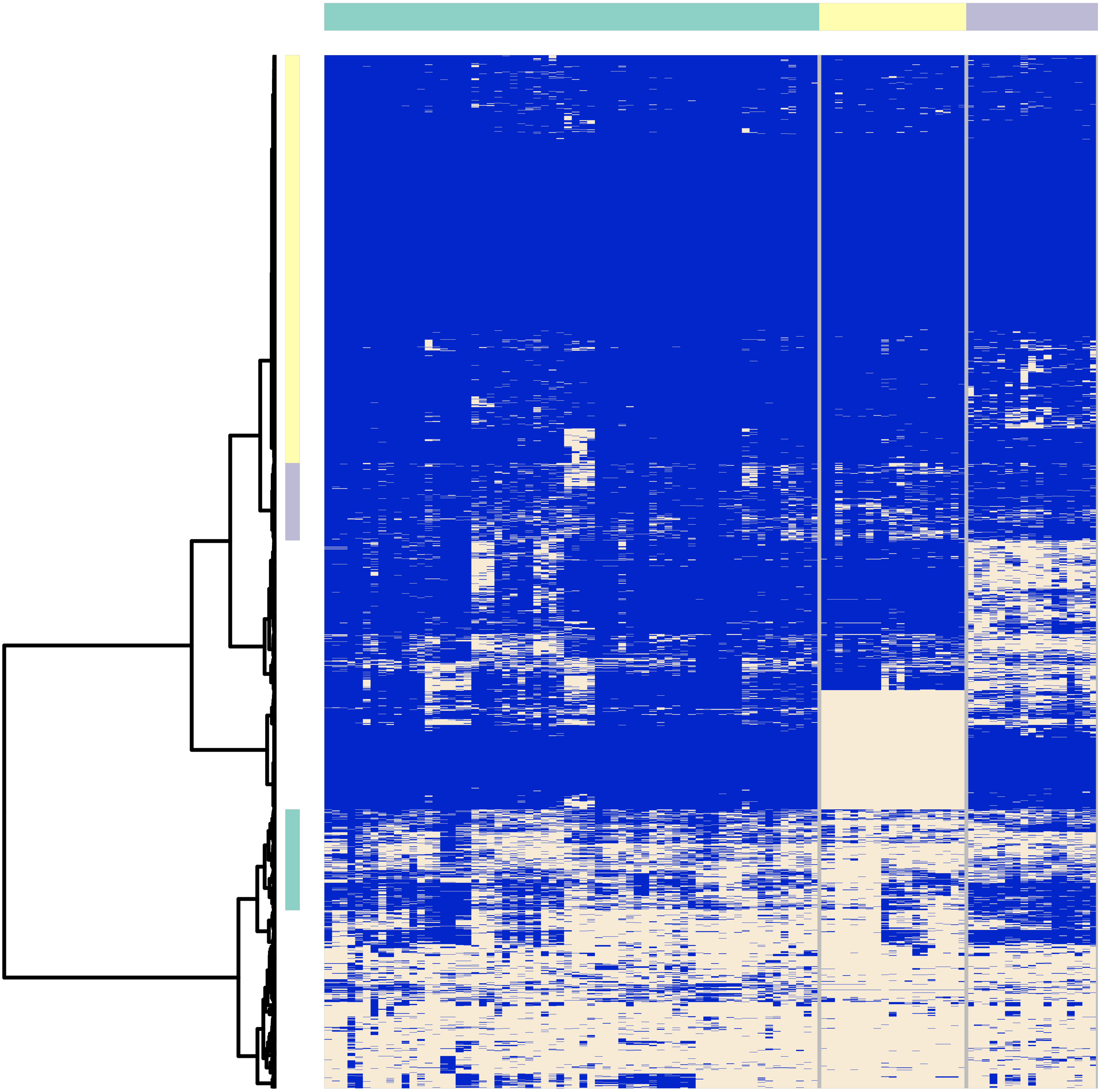

**Prevotella_amnii**
**No. Samples=458 No. Genes=3488**

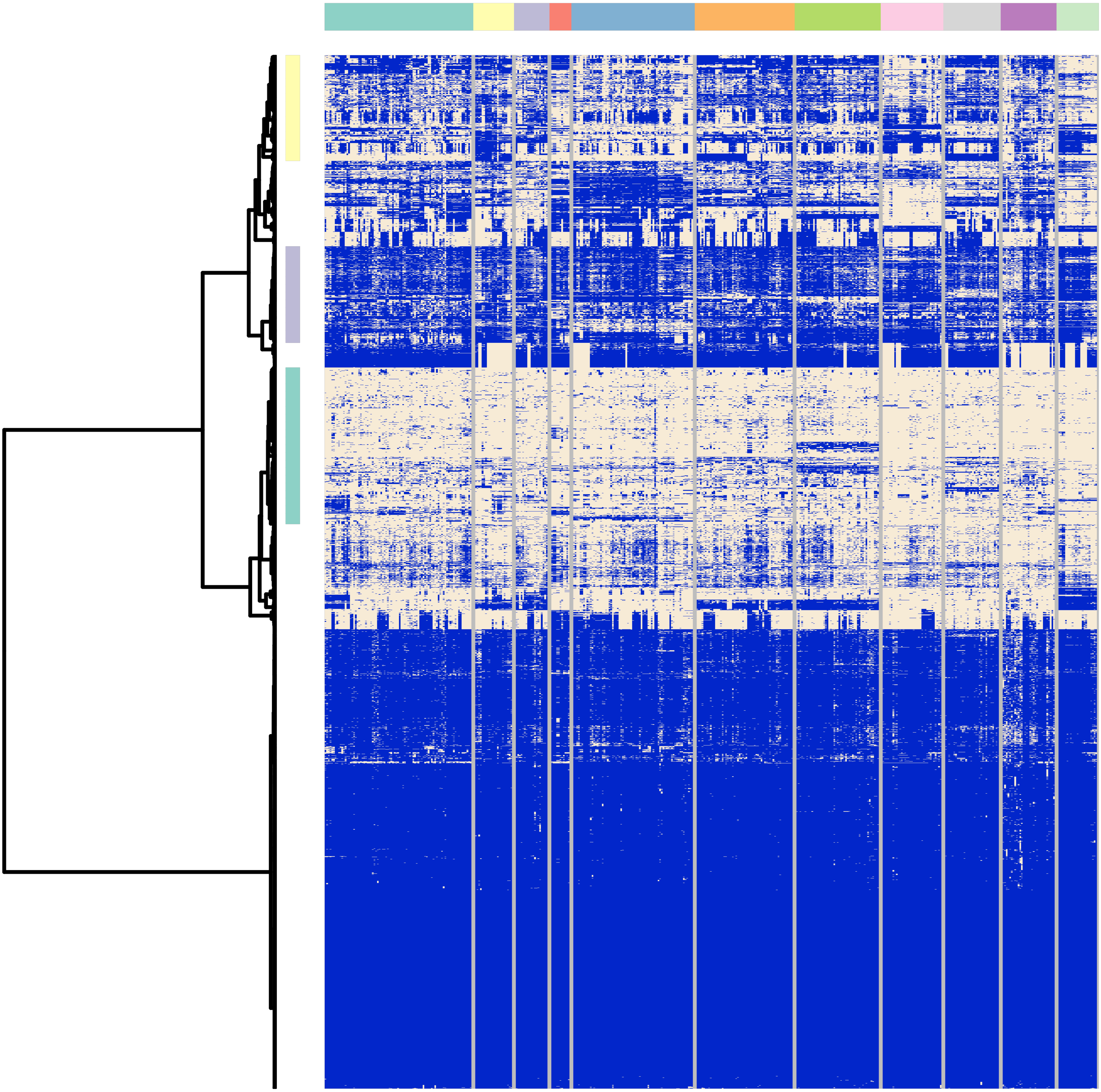

**Prevotella_bivia**
**No. Samples=255 No. Genes=2820**

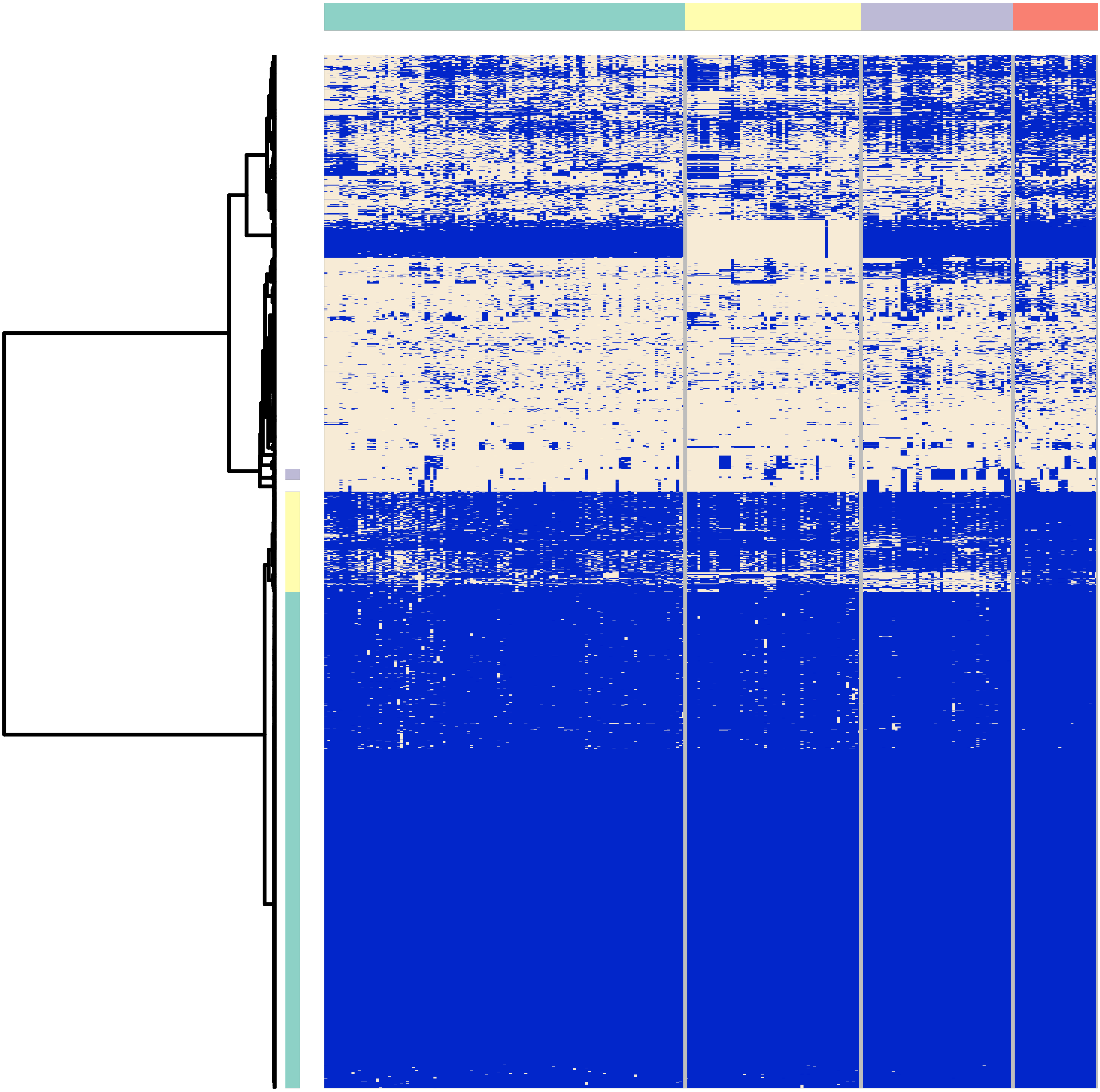

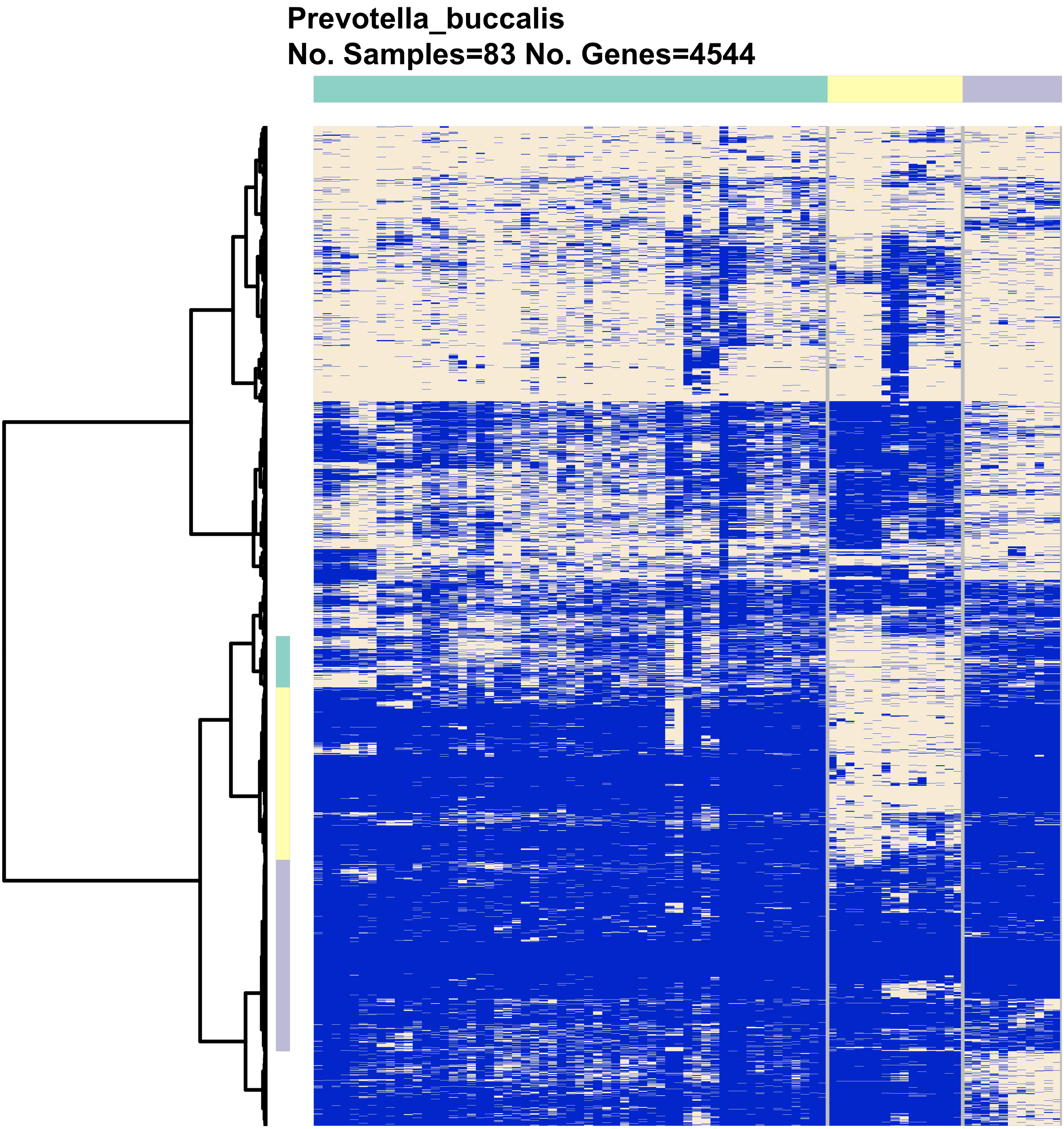
Prevotella_buccalis
No. Samples=83 No. Genes=4544

**Prevotella_disiens**
**No. Samples=29 No. Genes=2550**

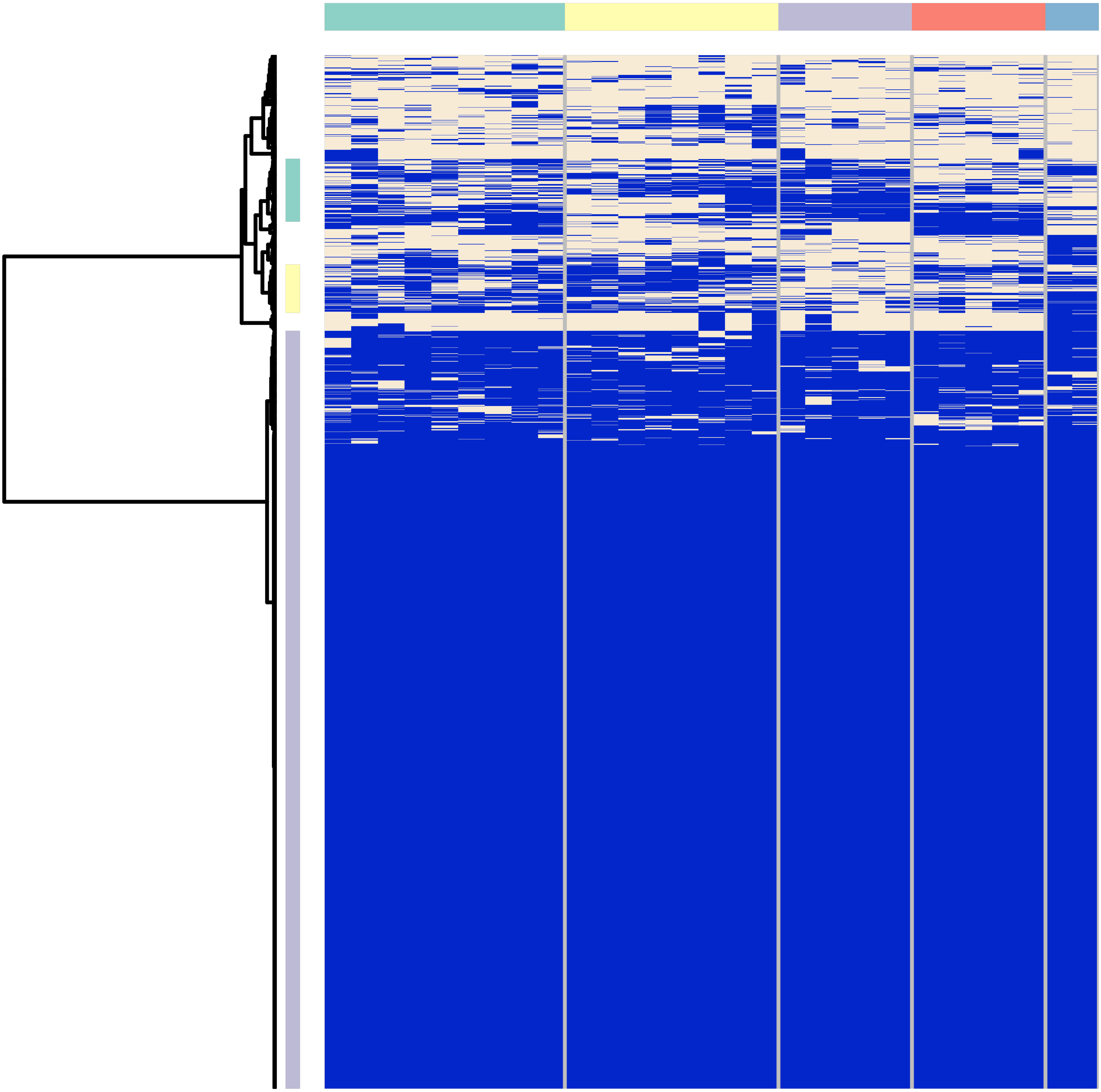

**Prevotella_sp.**
**No. Samples=328 No. Genes=2490**

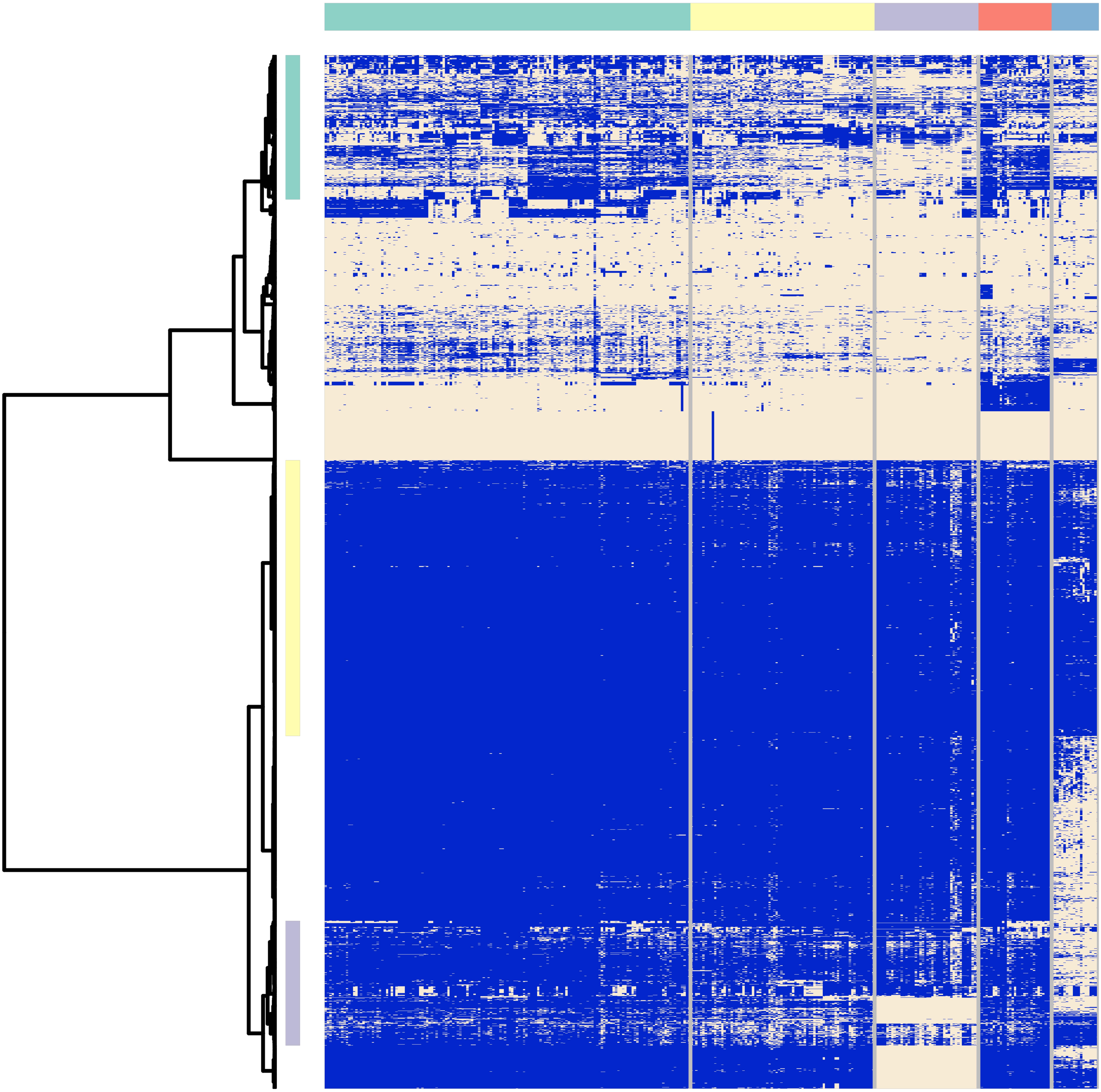

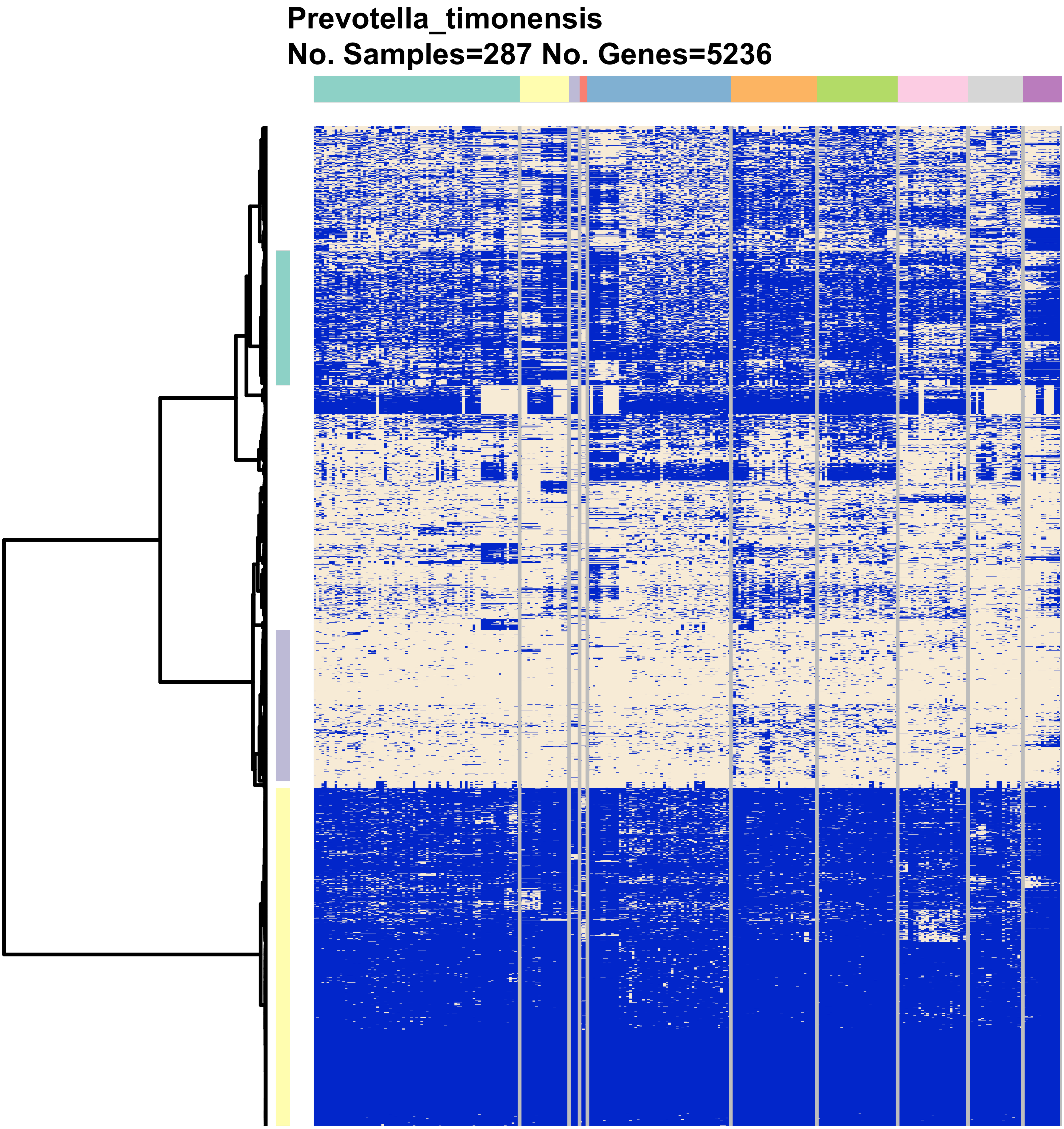
Prevotella_timonensis
No. Samples=287 No. Genes=5236

### Sneathia_amnii
### No. Samples=350 No. Genes=1116

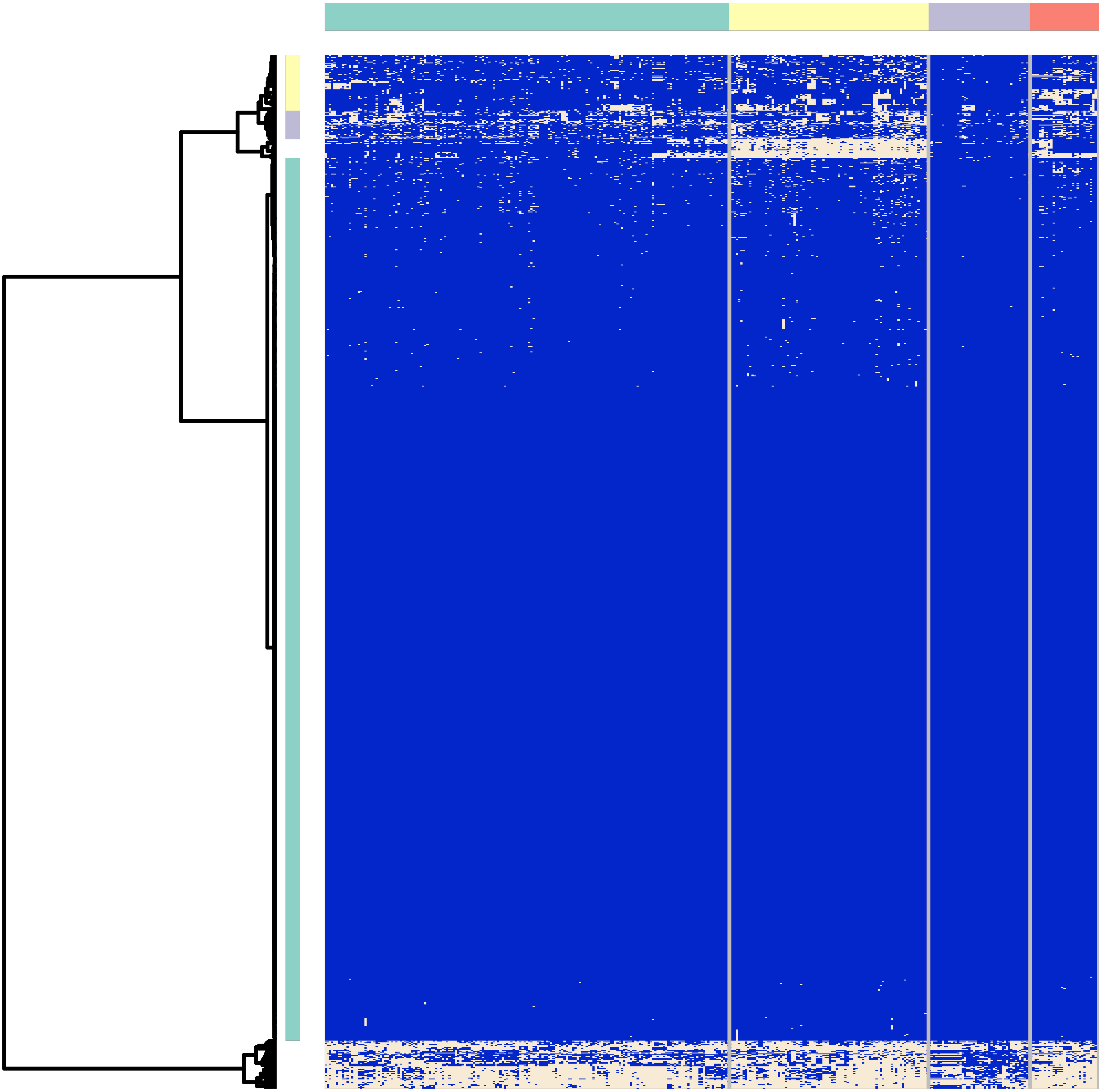

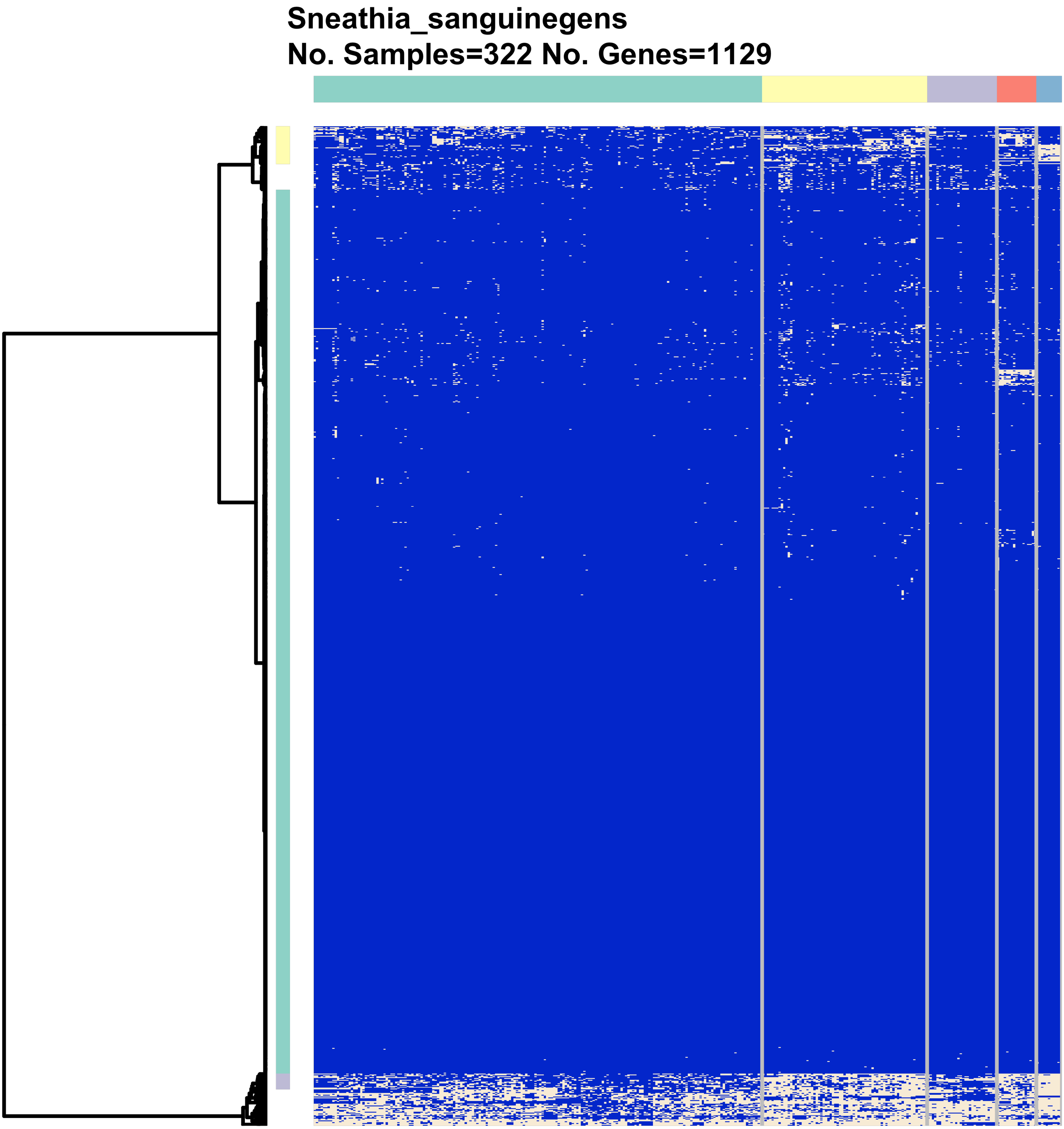
Sneathia_sanguinegens
No. Samples=322 No. Genes=1129

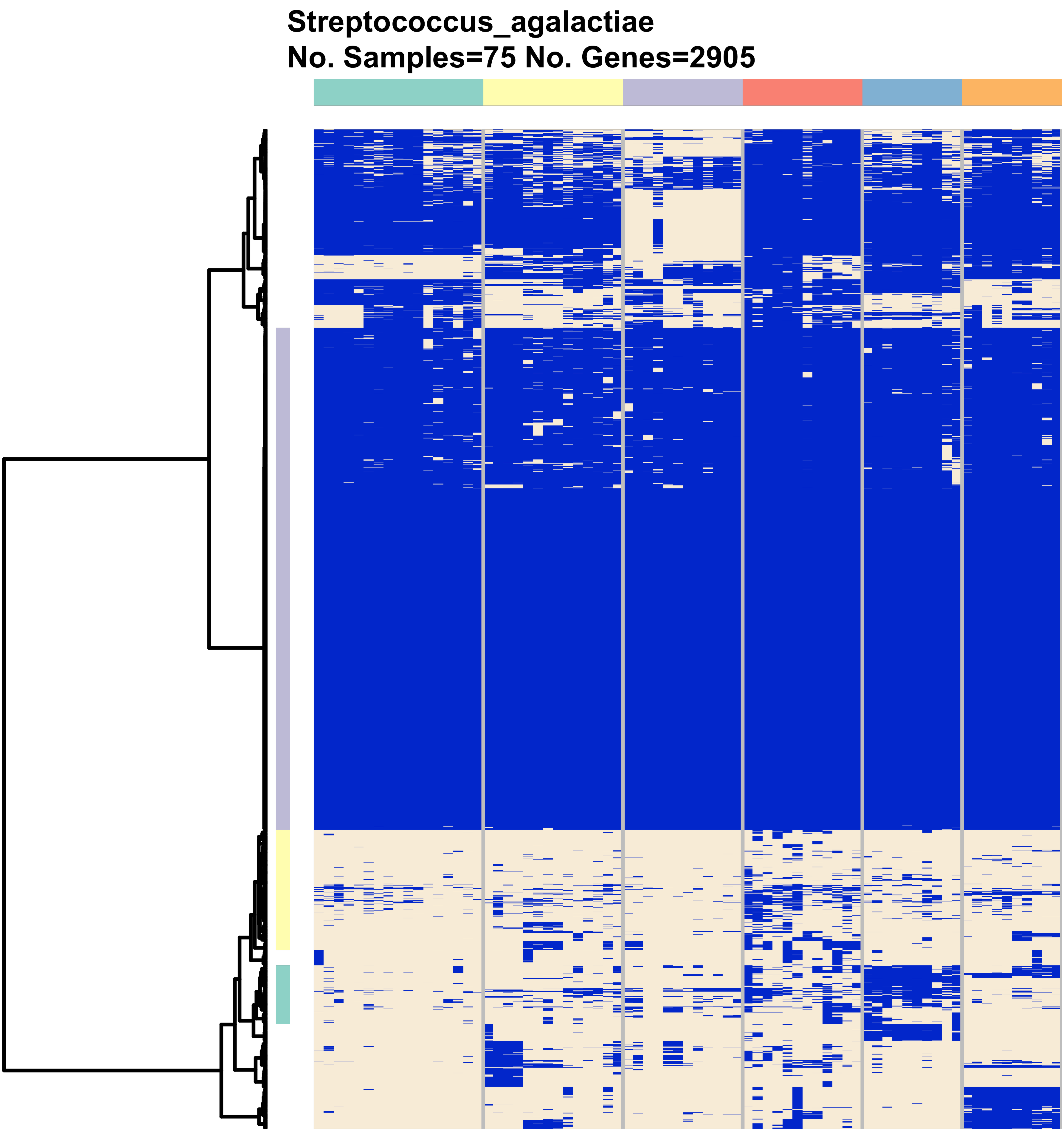
Streptococcus_agalactiae
No. Samples=75 No. Genes=2905
